# Ion beam subcellular tomography

**DOI:** 10.1101/557728

**Authors:** Ahmet F. Coskun, Guojun Han, Shih-Yu Chen, Xavier Rovira Clavé, Sizun Jiang, Christian M. Schürch, Yunhao Bai, Chuck Hitzman, Garry P. Nolan

**Affiliations:** Baxter Laboratory, Department of Microbiology and Immunology, Stanford University School of Medicine, Stanford, CA; Department of Radiology, Molecular Imaging Program at Stanford, Stanford University School of Medicine, Stanford, CA; Department of Chemistry, Stanford University, Stanford, CA; Department of Materials Science and Engineering, Stanford University, Stanford, CA

## Abstract

Multiplexed ion beam imaging (MIBI) has been previously used to profile multiple parameters in two dimensions in single cells within tissue slices. Here, a mathematical and technical framework for three-dimensional subcellular MIBI is presented. We term the approach ion beam tomography (IBT) wherein ion beam images are acquired iteratively across successive, multiple scans and later compiled into a 3D format. For IBT, cells were imaged at 0.2-4 pA ion current across 1,000 axial scans. Consecutive subsets of ion beam images were binned over 3 to 20 slices (above and below) to create a resolved image, wherein binning was incremented one slice at a time to yield an enhanced multi-depth data without loss of depth resolution. Algorithmic deconvolution, tailored for ion beams, was then applied to the transformed ion image series using a hybrid deblurring algorithm and an ion beam current-dependent point-spread function. Three-dimensional processing was implemented by segmentation, mesh, molecular neighborhoods, and association maps. In cultured cancer cells and tissues, IBT enabled accessible visualization of three-dimensional volumetric distributions of genomic regions, RNA transcripts, and protein factors with 65-nm lateral and 5-nm axial resolution. IBT also enabled label-free elemental mapping of cells, allowing “point of source” cellular component measurements not possible for most optical microscopy targets. Detailed multiparameter imaging of subcellular features at near macromolecular resolution should now be made possible by the IBT tools and reagents provided here to open novel venues for interrogating subcellular biology.

---

Multiparameter single-cell phenotyping for nucleic acids and proteins has revealed changes in molecular complexity that occur during development and throughout cancer progression in manners that reveal underlying mechanism and potential therapeutics.^1,2^ Localization within individual cells of biologic constituents with high-dimensional analytic techniques provides further depth to such understanding.^3,4^ Multiplexed ion beam imaging (MIBI) can be employed to spatially visualize multiple (6-50) protein parameters in histology sections,^5,6^. Since multiplexed fluorescence-based profiling methods have revealed identities of cells in tissues and suggested functions of novel cellular neighborhoods^7–9^, IBT should allow us to expand such neighborhood concepts to the subcellular realm and in 3D.

Super-resolution microscopy^10,11^ and 4pi single-molecule microscopy^12^ have been employed for 3D subcellular analysis, but these techniques are currently limited to handful of parameters and can be cumbersome to implement for whole cells. Mass spectroscopy offers unique advantages with its multi-parameter and high-resolution imaging capability, and recent efforts in secondary ion beam spectroscopy (SIMS), specifically OrbiSIMS^13^, has demonstrated the potential of multiplexing by metabolic imaging, but the spatial resolution was limited to 300 nm. To date, primarily lipid profiling has been implemented with OrbiSIMS largely due sensitivity limits of given target analytes. Another implementation of SIMS, NanoSIMS imaging, directly overcomes the optical diffraction limit ^14–16^, but it has not been applied to date for error-reduced whole-cell volumetric imaging.

Here we present ion beam tomography (IBT), a technical, reagent, and mathematical pipeline for analysis of ion beam images acquired from continuous depth scans of single cells. IBT achieves high parameter multiplex detection (7 to potentially 50 markers), subcellular 65-nm lateral resolution, and 5-10 nm axial resolution. To assess spatial resolution capabilities and sensitivity of IBT, we developed a suite of reagents and algorithmic approach to analyze subcellular structures and metabolites in single cells. Specifically, we developed methods to use IBT to measure spatial distributions of nuclear architectural features, including replication forks, and newly synthesized and mature mRNAs in cultured cells. Subcellular tomographic data will enable visualization and quantification of normal and diseased intracellular states and the synthesis of high-dimensional histology data with subcellular 3D mapping will intersect with basic biology and medical practice. Systematic and hierarchical subcellular organization of metabolic progression of single cells were consistent observations from the presented ion beam tomographic information cubes based on the computational association analysis of isotopically tagged replication and transcription sites near chromatin.

## RESULTS & DISCUSSION

### Design of ion beam tomography methods and analysis approaches

Ion beam imaging enables visualization of the elemental composition of a cell specimen on silicon substrates under bombardment with primary ions (**Fig. 1a**). A thin layer of cellular material (<5-10 nm) is etched during each raster scan, a process that generates secondary ions from the etched layer. The released ions can be analyzed by mass spectroscopy to identify endogenous structures or labeled molecules. Two-dimensional (2D) scans of ion distributions have been previously used to produce spatial maps of cell materials at each layer.

**Figure 1.**
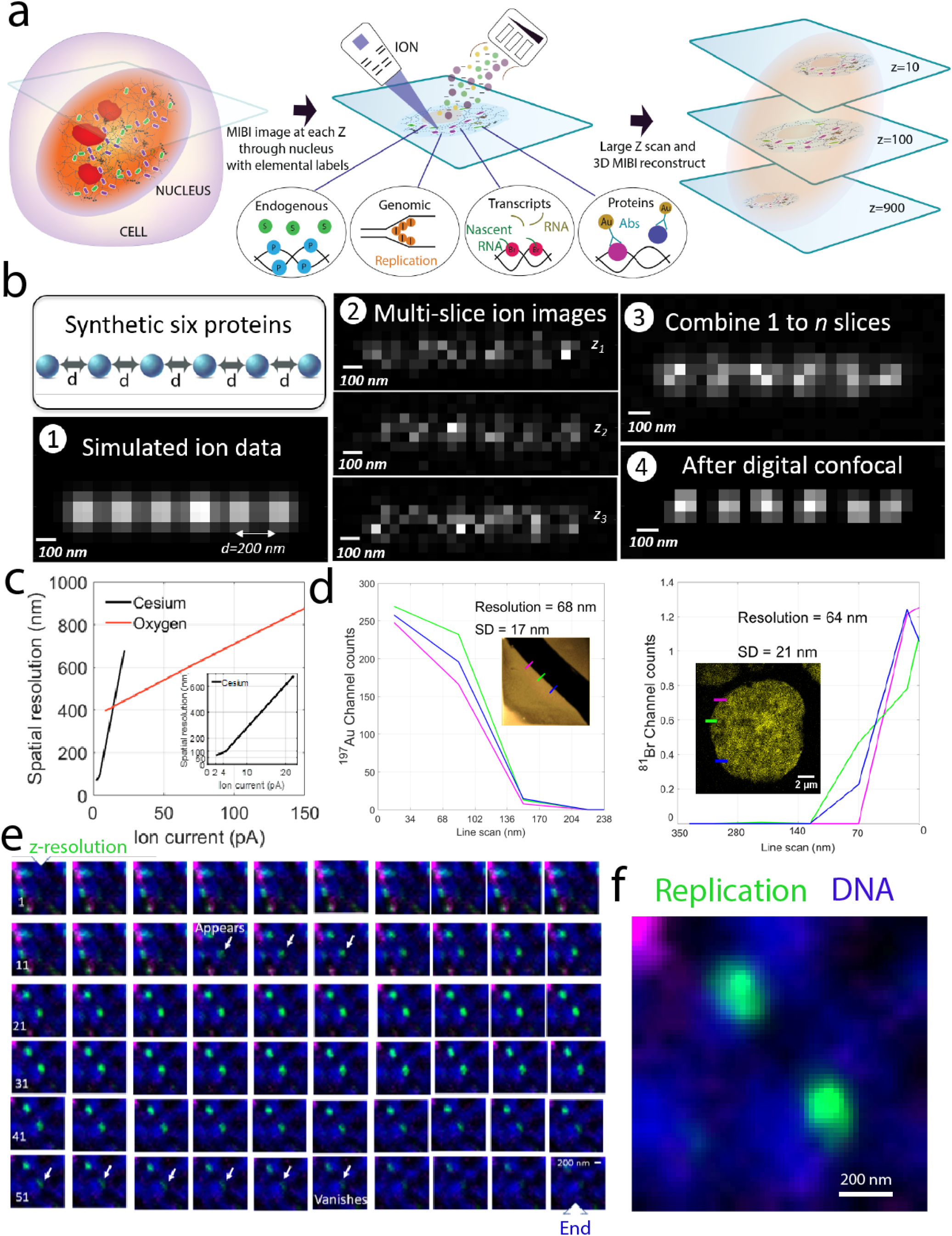
Ion beam image reconstruction framework achieves high 3D resolution with multiplexing. (a) Endogenous markers, genomic regions, transcripts, and protein factors were visualized by metabolic labeling or metal-tagging in ion beam imaging. Continuous ion beam scans of up to 1,000 depth sections were performed across the entire cell. (b) Validation of IBT using synthetic ion beam data.

1. Six simulated protein molecules separated by 200 nm were imaged with 50-nm pixel size and 120-nm ion beam width.
2. The original ion image was contaminated by digital multiplicative noise, providing simulated ion images at three different depths (z_1_, z_2_, and z_3_).
3. Multiple depth images of 1 to *n* slices (with sliding windows of 5 slices) were summed.
4. Binned ion images were digitally deconvolved to deblur the objects, resolving the neighboring pixels. The resultant image agrees well with distribution in the synthetic image, validating the accuracy of reconstructions. (c) Lateral resolution of IBT was evaluated by plotting spatial resolution vs. ion current for cesium and oxygen beams. (d) IBT provides spatial resolution down to 68 nm ± 17 nm (standard deviation) in gold-silicon interface and 64 nm ± 21 nm (standard deviation) in a DNA image of a Jurkat cell. Inset shows high resolution performance of the cesium source. Data presented in these resolution experiments were collected at 0.155 to 4 pA for the cesium beam. (e) To evaluate axial resolution, a 175-nm wide DNA replication pattern (green) that crosses about 40 ion image slices (z sections labeled as 1, 11, 21, 31, 41, and 51 with increasing slice number in the same row) was imaged, demonstrating around 5-nm depth resolution. Blue is the total DNA signal. (f) Zoomed image of distinct replication forks (green) around DNA (blue) at the 36^th^ section. Ion image was interpolated 3× to avoid pixelation issues. Scale bar: 200 nm.

Previous applications of MIBI utilized an oxygen duoplasmatron primary ion beam to generate 2D profiles of tissues at 260 to 500-nm spatial resolution with lanthanide isotopes as mass-based reporters.^5,6^ Here, a different set of halogens and post-transitional and noble elements were used as contrast agents and a cesium ion beam was employed, yielding 2D ion images at sub-100-nm lateral resolution due to the small spot size (<50 nm) of the beam. Iterative scanning of a cell through the Z axis generates multiple 2D images. Each 2D ion image series contains 3D information about the cellular volume since the ion beams extract ions slightly below (out of focus from) the ablated layer. This makes it challenging to register and render images.

We first developed a mathematical framework and a computational method to allow error-minimized 3D ion beam mapping of cellular volumes (**Fig. 1a and Supplementary Fig. 1**). The resultant images were digitally assembled to generate volumetric renders of cells. Synthetic ion beam data were first used to validate the IBT approach (**Fig. 1b**). In the image used for validation, the true object distribution was a synthetic array of six virtual molecules separated by 200 nm and modeled as digitally acquired by a 120-nm ion beam width and 50-nm scanning pixel size on x and y axes (**Fig. 1b, step 1; Supplementary Fig. 2**). Only multiplicative noise is significant in ion beam imaging due to the independent fluctuations of ion pixel values for each frame; it is present as a gamma distribution and was calculated from a sample image acquired at similar ion imaging parameters. The convolved image was then contaminated by multiplicative noise factors. The digitally created images at distinct depths (z_1_, z_2_, and z_3_) were typical of noisy data (**Fig. 1b, step 2**). These multiple depth images were binned over a sliding window of size *n* that was varied between 3 and 20 slices (**Fig. 1b, step 3**). This process was then incremented by one layer at a time. For instance, ion signals from 1 to *n* slices were combined to yield the first transformed image, ion signals from slices 2 to *n*+1 were binned for the second transformed image, and slices 3 to *n*+2 yielded the third transformed ion image without the loss of axial resolution. This rendered an image series of 1,000 slices into a restructured image series of a similar length (slightly shorter, reduced only by the sliding window size). Each binned section was then digitally deblurred by an iterative Lucy-Richardson^17^ deconvolution method (**Fig. 1b, step 4**). The Lucy Richardson algorithm is suitable for experimental point spread function (PSF) and iterative analysis; however, changes in electric current levels due to the ion source quality and parameters of the ion beam imaging system make it challenging to measure a unique PSF for every ion image of interest. To address this problem, a “hybrid” deconvolution algorithm was created by using a blind PSF rather than an experimentally determined PSF. The width of the blind PSF is selected based on the numerical value of the ion current recorded before and after the ion beam scanning. The width of the blind PSF is typically in the 100-200 nm range; for the simulation described, the PSF was 120 nm. The final ion image showed six distinct signatures that agreed well with the synthetic input.

To quantify and validate the spatial resolution of IBT, fabricated metal-coated silicon substrates with a sharp interface of gold (^197^Au) and silicon were imaged at different current levels (**Fig. 1c**). Using the IBT mathematical analysis pipeline, 68-nm lateral resolution was obtained using a cesium source from three edge scans (**Fig. 1d and Supplementary Fig. 3**). Here, the resolution was defined as the distance between 84% and 16% of the maximum ion beam signal. With these settings a presumptive 65-nm resolution image was obtained for ion beam images of a Jurkat cell (**Fig. 1d and Supplementary Fig. 4**). Using the oxygen source, only a 395-nm maximum resolution was obtained from three edge scans at the junction of aluminum and silicon (**Supplementary Fig. 5**). Thus, the resolution depends on ion current levels and aperture size. As expected a cesium beam allowed for higher resolution imaging than an oxygen beam. Therefore, with a scalable resolution from 65 nm to 600 nm, ion beam imaging bridges a gap between super-resolution imaging and wide-field microscopy. The axial depth resolution of the ion beam imaging was evaluated by two approaches. In the first, a single replication site (marked by a 30-min incorporation of 5-^127^iodo-2′-deoxyuridine (^127^I-dU)) was evaluated: A 175-nm wide pattern crossed 40 slices, indicative of 5-nm axial resolution (**Fig. 1e and Supplementary Figs. 6 and 7**). In the second approach, the cell height (∼10 µm) was divided by the total number of scans (1,000), again indicative of 5-10 nm axial resolution (**Supplementary Fig. 8**).

IBT image acquisition settings must be carefully adjusted to obtain accurate reconstructions. Images are acquired at a low current level (<4 pA) for high-resolution analysis, yielding ion beam raw images at sub-100 nm resolution. Pixel size dimensions, Δx and Δy, of these ion beam images must be smaller than the ion beam width of *Δw* (**Supplementary Figs. 9-10**). To reconstruct these finely sampled experimental images, a PSF width between 100 and 200 nm was utilized in the deconvolution process. Another parameter that affects the sensitivity in the ion beam imaging is ion conversion efficiency; this is defined as the ratio of ion incoming signal (*P*_*in*_) to the total secondary ion extracted from the sample (*P*_*out*_) and was calculated to be 3%. The captured ion beam images are typically fuzzy due to the finite ion beam width, imperfect image focusing, and out-of-focus etching. Therefore, multiple depth binning was used to enhance the signal-to-noise ratio (**Supplementary Fig. 11**), and hybrid deconvolution methods were used to transform raw images into high-resolution ion beam image reconstructions.

Optical microscopy often suffers from multi-color aberrations that limit co-localization of labeled regions. IBT benefits from simultaneous etching of the same pixel regions in the channel of interest without any lateral shifts. Corrections are required, however, due to general image offset during depth scans. To enable these corrections, a fiducial pattern is required that allows detection of spatial shifts of each depth image. To provide this pattern, iron (^55^Fe) microparticles with diameter of 2-6 µm were added to cell samples, and the center and edge of the particle was tracked during the ion scans to allow image shift corrections (**Supplementary Fig. 12**). These shift values were then applied to other mass channels creating error-reduced, 3D cellular maps.

Subcellular IBT experiments provide with tomographic, continuous acquisition of multiple depths in a few cells per day. To increase the throughput of IBT approach, a “chained mode” was used to image at high-low currents by alternating image acquisition (**Supplementary Fig. 13**). In this strategy, a high-resolution scan was followed by a low-resolution scan, and this process was repeated until the bottom of the cell was reached. Chained mode preserves lateral resolution at sub-100-nm spatial details, but it loses IBT’s high axial resolution as the low-resolution scans etch through deeper (more than 50 nm) cellular layers. At the expense of spatial resolution, a higher current can image through the entire cell in a shorter timeframe to increase cell numbers evaluated (**Supplementary Fig. 14**).

### Tagging, labeling systems, and post-processing of IBT data

Previous MIBI experiments utilized mostly lanthanides as reporters for tissue-labeling experiments and oxygen or gold ion beam sources (**Fig. 2a**). For IBT, a different set of elemental tags were required to enable a higher contrast levels when imaged by a sub-100 nm spot size of a cesium source. Sensitives were based on the efficiency of secondary ion beam generation (**Fig. 2b and Supplementary Fig. 15**). Previous high-resolution ion beam imaging experiments have relied on metabolic enrichment of ^2^H, ^13^C, ^15^N, and ^18^O to measure turnover rates and to monitor cell division.^14,15^ As a proof-of-concept for the IBT workflow, the following were detected in Nalm6 cells using a cesium beam: ^12^C as a measure of carbon distribution, ^14^N (measured as ^12^C^14^N) as a measure of nitrogen distribution, ^34^S as a measure of total protein, ^31^P as a measure of nucleic acids, ^127^I-dU for labeling of replication sites, and 5-^81^Br-ribouridine (^81^Br-rU) for labeling of newly synthesized transcripts (**Fig. 2c, step 1**). Raw images were improved by multi-slice binning (**Fig. 2c, step 2**), and digital deconvolution of binned images (**Fig. 2c, step 3**). After performing these steps for each ion beam depth image, 3D representations in the form of analog pixels values (equivalent to 3D cell tomograms in terms of chromatin, replication, or transcription) and surface-rendered volumetric reconstructions were created (**Fig. 2c, step 4**).

**Figure 2.**
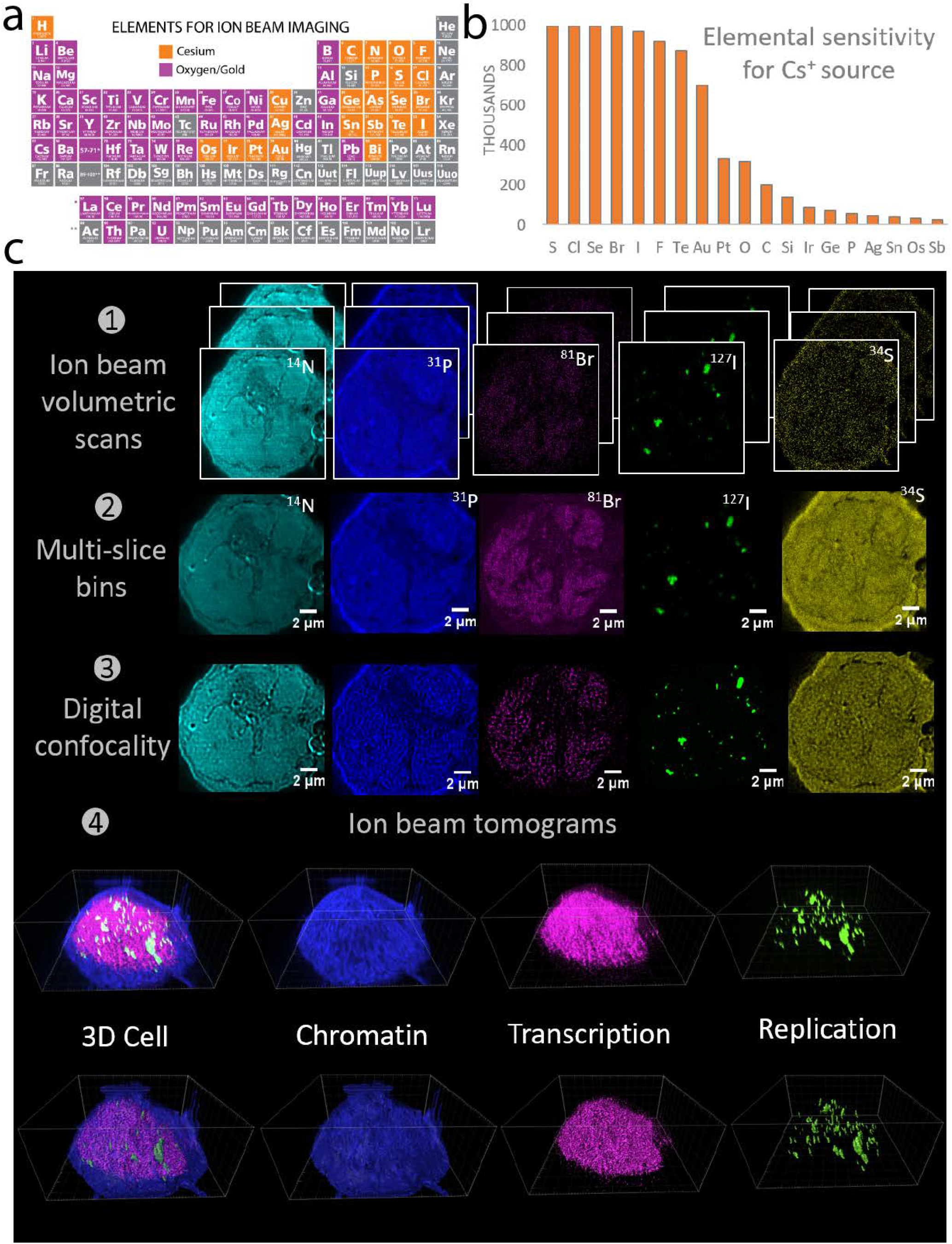
Metabolic tagging and endogenous elemental contrasts enable ion beam tomography. (a) Elements suitable for high-contrast labeling detected with cesium (orange) or oxygen/gold (purple) beams. (b) Sensitivity of indicated elements for cesium source imaging. (c) A single Nalm6 B cell lymphoblast imaged for ^14^N (measured as ^12^C^14^N, cyan), ^31^P for DNA (blue), ^81^Br for newly synthesized transcripts (labeled with ^81^Br-rU, magenta), ^127^I for replication loci (labeled with ^127^I-dU, green), and ^34^S (yellow) for detection of total proteins.

1. A volumetric scan across 1,000 slices was performed by ion beam imaging from top to the bottom of a cell for seven mass channels. Potentially 100 mass channels can be analyzed.
2. Slices were combined with a sliding window of 5 depth sections.
3. Binned ion images were digitally deconvolved to sharpen the images.
4. Ion beam tomograms are shown in raw image format (upper images) and 3D surface renders (lower images) with the three channels (selected out of seven acquired channels) combined as a 3D cell, in the form of chromatin, transcription, and replication tomographic representations.

Ion beam tomograms provide “snapshots” of subcellular structures; they do not provide direct measures of molecular changes over time. For instance, after a 30-minute incubation with ^127^I-dU, cells are fixed and the moment just before fixation is preserved and measured. To enable capture of dynamic information with ion imaging time-dependent labeling can be used during the sample preparation. To analyze replication dynamics, cells were sequentially pulsed with ^127^I-dU and ^81^Br-dU with chase times of 30 minutes or 2 hours, followed by immediate fixation and ion beam tomographic analysis. For transcription and replication co-dynamics analysis, cells were simultaneously pulsed with ^81^Br-rU and with ^127^I-dU for 30 minutes, followed by fixation and ion beam imaging (**Supplementary Fig. 16**).

After application of the ion current-dependent blind PSF to iteratively compute the sharpened image, the final ion images are visualized in the form of segmented objects, meshed voxels, volumetric renders, and molecular neighborhood maps. As in analyses of the 2D data obtained from histology and of the high-parameter single-cell mass cytometry data,^1,5^ analysis of 3D ion beam data requires dimensionality reduction from pixel positions (x, y, and z). Octree analysis provides a direct solution to dimensionality reduction in which voxel arrays are subdivided into spatial volumes by eight neighboring octants.^18^ Iterative Octree reduced dimensions of ion images to only a few digital numbers that represent the cell state as demonstrated for replication dynamics analysis in **Supplementary Fig. 16**. Our lab recently demonstrated the existence of novel organized cell neighborhoods related to tissue function in the immune system.^9^ In the next section here such spatial network concepts were implemented to study chromatin interactions with other molecular types such as proteins in complex tissue samples. To further study subcellular structural organization in ion beam tomographic information cubes, hierarchical clustering, correlation and association maps were generated across subcellular patterns with various metabolic labeling conditions.

### IBT reveals nuclear architecture, metabolites, and dynamics

#### Analysis of chromatin conformation

Chromatin has multiple levels of super-structural complexity from 11-nm DNA-bound mono-nucleosomes to 30-, 120-, and 700-nm fibers^19^. Mono- or di-parameter direct visualization of higher order chromatin details has been performed by super-resolution optical microscopy (STORM, spatial details: 30-100 nm) and enhanced electron microscopy (ChromEM, spatial details: 5-100 nm).^20,21^ Tomographic imaging of chromatin remains challenging, however. Optical microscopy requires many frames per cell and electron microscopy requires physical sectioning of cells; both pose difficulties for 3D registration. In contrast, IBT directly records spatial details of chromatin at sub-100 nm resolution, with multiple parameters, without the need for sectioning or multiple frames per depth.

Chromatin 3D conformation was analyzed by IBT using two different methods. First, replicated chromatin was detected based on I-dU and Br-dU incorporation into the newly synthesized DNA (**Supplementary Fig. 17)**. In this experiment, HeLa cells were cultured with ^127^I-dU and ^81^Br-dU for 24 hours, followed by fixation, and ion beam tomographic imaging across 700 depths. Raw ion images and reconstructed ion tomographic slices are presented for 30^th^, 225^th^, and 550^th^ slices in **Fig. 3a**. ^127^I-dU and ^81^Br-dU ion images should correspond to the *same* chromatin pattern in an individual HeLa cell due to the incorporation of the same base (thymidine) into the DNA. Although spatial distributions of pixels in raw ion images for each depth were significantly dissimilar (structural similarity index, SSIM, value: 0.54 ±SE 0.0037, **Supplementary Fig. 18**) reconstructed ion images were highly correlated (SSIM value: 0.86 ±SE 0.001), providing a validation of the presented mathematical processing methods. A parallel experiment in a Nalm6 cell provided further validation (**Supplementary Fig. 19**). Second, chromatin was identified by analysis of the ^31^P signal. Since all nucleic acids as well as phosphorylated proteins contribute to the endogenous ^31^P signal, the phosphate channel image differs slightly, as expected, from the ^127^I-dU and ^81^Br-dU images (**Fig. 3a**). Even though ^31^P is present in additional molecular entities, this image demonstrates that the endogenous ^31^P channel can be used to identify chromatin architecture without the need for additional metabolic or secondary tagging.

**Figure 3.**
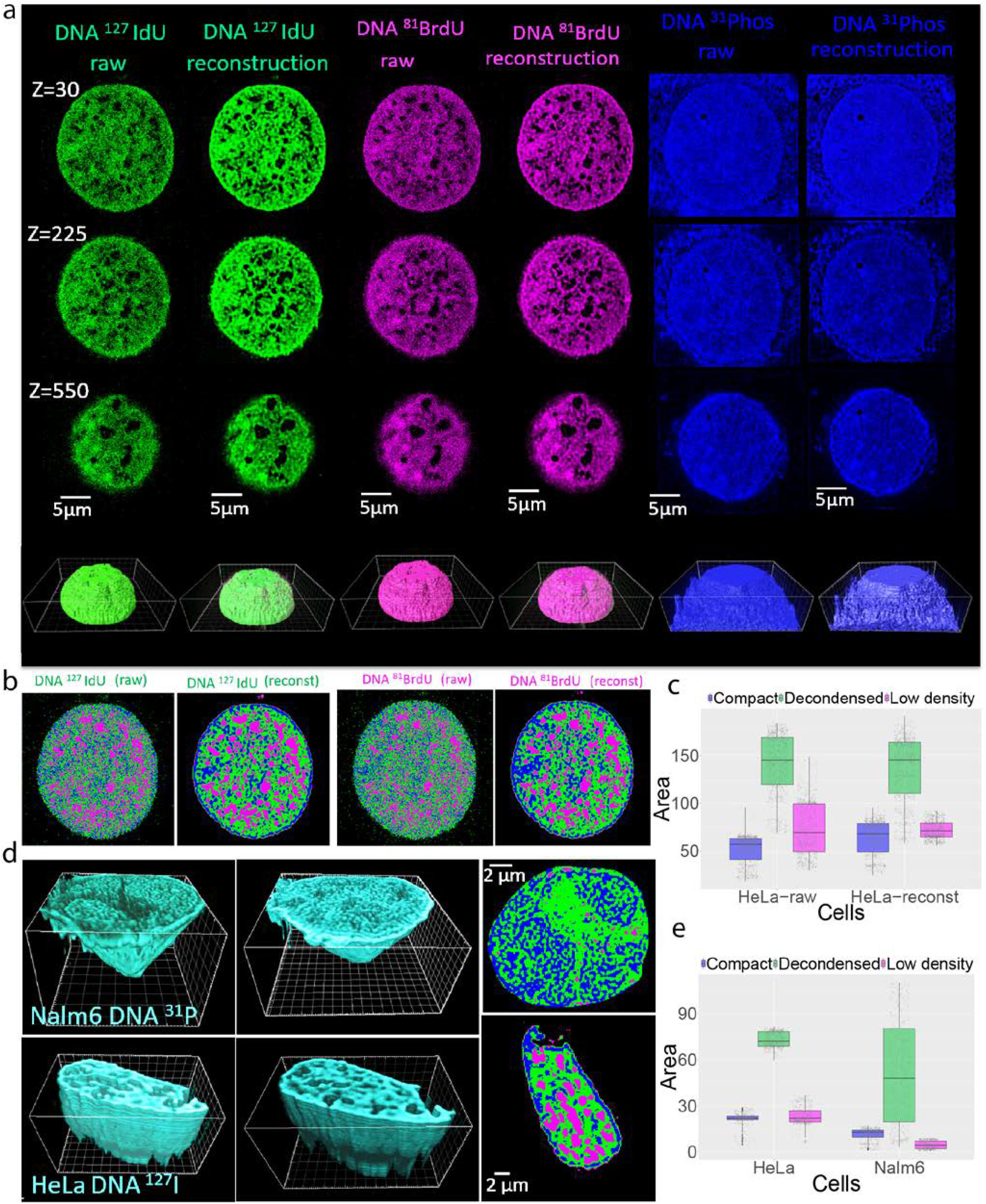
IBT for assay of chromatin states. (a) Thymidine analogues ^127^I-dU and ^81^Br-dU were simultaneously incorporated into cells for 24 hours. This allowed chromatin visualization in a single HeLa cell in two different mass channels, providing two distinct datasets to test our mathematical pipeline. The first column shows typical raw ion images from 30^th^, 225^th^, and 550^th^ depths from the ^127^I-dU channel (green). At the bottom of each column, a 3D render of 700 slices is shown as an ion beam cell tomogram. The second column presents reconstructed ^127^I-dU images; these are sharper and better reveal spatially resolved features of chromatin fibers than the raw images. The third and fourth columns correspond to raw and reconstructed images of the chromatin obtained by analysis of the ^81^Br-dU channel (magenta). Reconstructed images in the second and fourth columns agree well with each other, validating the mathematical analysis. The raw images in the first and third columns show significant spatial differences, even though both I-dU and Br-dU label replicating DNA. The 3D renders in the bottom row show strong chromatin features only in the renders from reconstructed ion images. The renders based on the raw images show random fluctuations due to the noise levels of each ion beam image. The fifth and sixth columns show the ^31^P channel, which is dominated by signal due to phosphate in the DNA backbone of the nucleus. As proteins and RNAs in both cytosol and nucleus contribute to this signal there are differences compared to the ^127^I-dU and ^81^Br-dU channel images. (b) To explore local regions of chromatin, an image segmentation was performed on the raw and reconstructed images of the ^127^I-dU and ^81^Br-dU images. A 225^th^ depth image was partitioned into three colors based on the density of the chromatin using a fuzzy logic segmentation algorithm: compact chromatin (blue), decondensed chromatin (green), and low-density chromatin (magenta). Raw images in both channels failed to resolve fibers, whereas the three colors are present in similar spatial patterns in ^127^I-dU and ^81^Br-dU reconstructed images. Each data point corresponds to a single ion beam depth slice from top to bottom of a single cell (*n*=1,000). (c) A bar chart showing area occupied by chromatin forms in raw and reconstructed images (median values were used). (d) Upper images: Chromatin features in a Nalm6 cell imaged using ^31^P channel. Lower images: HeLa cell imaged using the ^127^I-dU channel, followed by reconstructions and segmentation. First and second columns show cross-sections of the chromatin at the 500^th^ slice and the 400^th^ slice, respectively. Segmented images of ^31^P in the Nalm6 cell exhibits minimal low-density chromatin signal that might be due to other background signal from proteins and RNA, whereas segmentation in ^127^I provided all three partitions of the chromatin in the HeLa cell. (e) Bar plot of areas occupied by chromatin forms across about 350 sections (each data point represents a single section) in these the Nalm6 cell imaged in the ^31^P channel and the HeLa cell imaged using the ^127^I channel. Cell-type differences and two different channels contribute to the heterogeneity in the compact, decondensed, and low-density regions.

Cancer cells and aberrant immune cells (B cell lymphoblasts) typically have large and deformed nuclei with protein compositions and chromatin conformations distinct from those of normal cells;^22,23^ very little is known about the subcellular chromatic super-structures in these abnormal cells. To resolve spatial details in chromatin states, HeLa cells and Nalm6 cultures were incubated with ^127^I-dU and ^81^Br-dU to label replicating DNA, and signals from ^127^I, ^81^Br, and ^31^P channels were analyzed (**Fig. 3a**). Spatial subregions were defined by the spatial distribution of these signals. A direct calculation of signal intensity may be used to segment the chromatin; however, a fuzzy logic image segmentation (see **Materials and Methods**) was used for this analysis to consistently classify voxel distributions into subgroups of chromatin density states regardless of the dynamic range differences of the ion beam image values.

Chromatin states were defined as described previously as compacted (a largely inactive but dense state of chromatin), decondensed (regions of medium density chromatin), and very low density (regions with little active transcription).^24^ A spatially resolved color visualization of an individual HeLa cell for 225^th^ ion beam image slice showed chromatin in all three states (**Fig. 3b**). To evaluate the effect of image reconstruction on chromatin segmentation, raw and reconstructed images were segmented (**Fig. 3c**). The raw images exhibited smaller amounts of compact chromatin and higher amounts of very low-density chromatin compared to the segmented results from reconstructed data. These results demonstrate how mathematical improvements can spatially resolve subcellular structures in ion beam images. Analysis of a Nalm6 cell (**Fig. 3d**) revealed cell-type specific spatial nuclear maps of chromatin states (**Fig. 3d-e and Supplementary Figs. 20 and 21**).

#### Study of spatial relationships of replication and transcription

High-resolution imaging of replication is crucial to understanding the metabolic effects of therapies in cancer, particularly for drug resistance studies.^25^ To visualize replication forks, Nalm6 cells were incubated with ^127^I-dU for 30-60 minutes, then for 30 minutes or 2 hours (the chase) without label, and then with ^81^Br-dU for 30 minutes. Cells were fixed, and the subcellular ion beam images were collected (**Fig. 4a**). These ion beam image series were then converted with a 3D renderer (**Fig. 4b**). Images of the cells chased for 30 minute showed significant overlaps in the ^127^I-dU and ^81^Br-dU channels. These data suggest that with the short chase, ^81^Br-dU labels replication sites that are also labeled by ^127^I-dU, whereas the longer chase provides higher spatial separation of replication sites (**Fig. 4c**). Pulse-chase experiments performed in S-phase cells (synchronized with aphidicolin) revealed similar spatial overlap patterns (**Supplementary Figs. 22-25**). The correlation coefficient of ^127^I-dU and ^81^Br-dU channels across 600 depths was calculated to quantify the overlap in signal. The correlation coefficient was 0.38 for the 30-minute chase and 0.13 for the 2-hour chase (**Fig. 4d**). This agrees well with super-resolution fluorescence microscopy analyses.^26^

**Figure 4.**
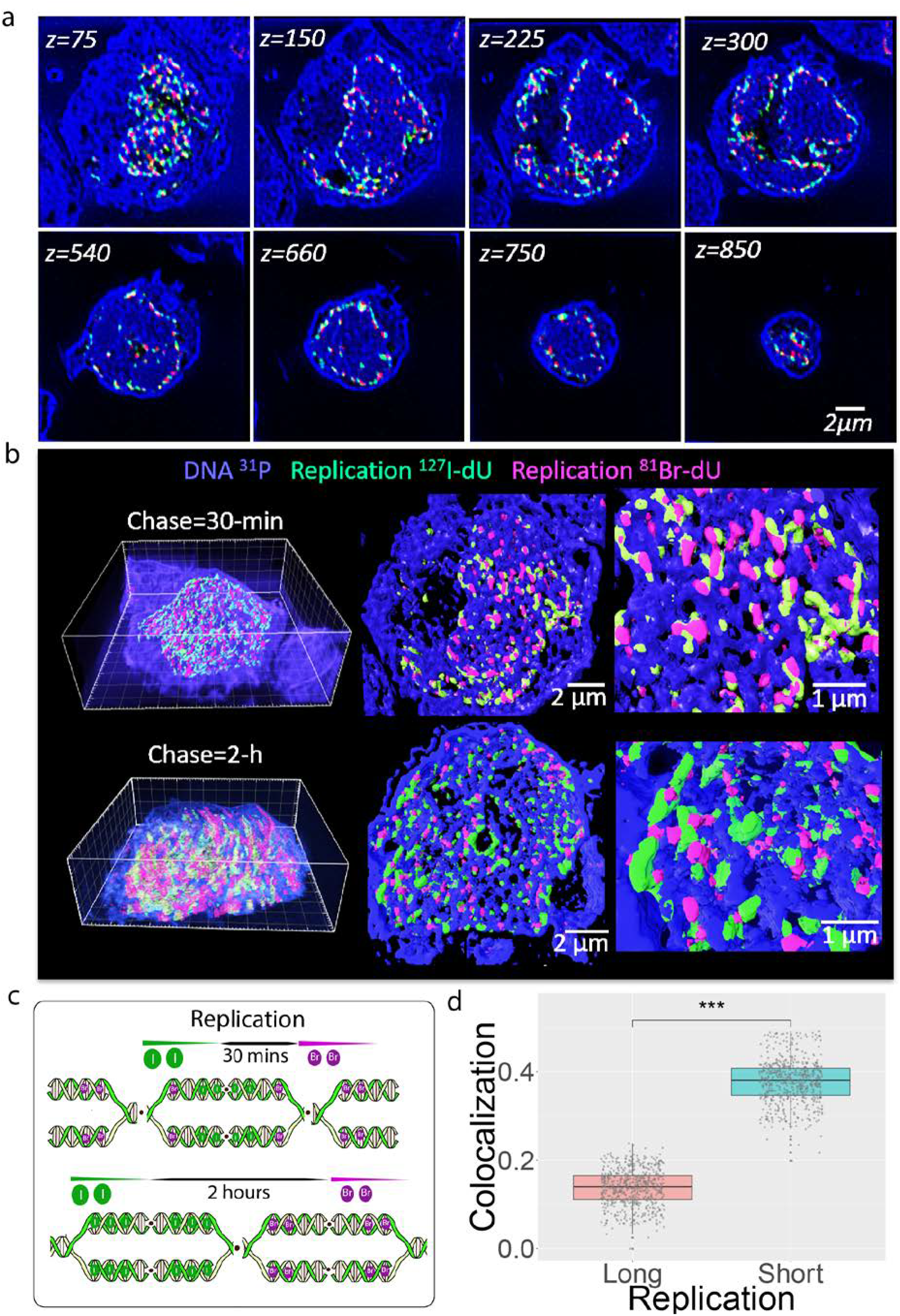
Ion beam tomograms reveal overlaps in the 3D distribution of DNA replication sites. (a) Nalm6 cells were incubated with ^127^I-dU for 30 minutes and then without label for 30 minutes or 2 hours. This chase was followed by a 30-minute incubation with ^81^Br-dU. Endogenous DNA backbone was visualized in the ^31^P channel (blue) and replication sites were detected in the ^127^I-dU (green) and ^81^Br-dU (red) channels. Reconstructed 2D ion beam images are shown from the 75^th^ slices to 850^th^ slice, clearly demonstrating the cell sections from the top to the bottom of the cell. (b) Upper images: 3D render of ^31^P (blue), replication sites labeled with ^127^I-dU (green), and replication sites labeled with ^81^Br-dU (magenta) across 1,000 slices with 30-minute chase (upper images) and 2-hour chase (lower images). The 100^th^ slice and 2D spatial distribution of replication sites are illustrated in the middle and right images, respectively. Lower images: 3D render of ^31^P (blue) across 800 slices. In middle and right images, the 100^th^ slice and the 2D spatial distributions are presented for chromatin with^31^P (blue), replication sites labeled with ^127^I-dU (green) and ^81^Br-dU (magenta). Spatial overlaps are much lower than in the cell subjected to the 30-minute chase. (c) Schematic of how the shorter chase time (30 minutes) incorporates the second color into the same replication site, leading to higher spatial overlap than the longer chase. (d) Bar plot of co-localization of ^127^I-dU and ^81^Br-dU channels for the 2-hour (long) and 30-minute (short) chase experiments. Each data point corresponds to co-localization values per ion beam depth slice. The difference of these two conditions exhibited significant P-value (***) by Wilcox test.

To analyze chromatin states and DNA and RNA synthesis, ^127^I-dU and ^81^Br-ribouridine (^81^Br-rU) were used to monitor synthesis of DNA and RNA, respectively, in three dimensions. The mutually exclusive mechanisms of transcription and replication have been widely studied by sequencing but not by multiplex imaging.^27–29^ Nalm6 cells synchronized in S phase or not were incubated for
 30 minutes or 2 hours with both ^127^I-dU and ^81^Br-rU, followed by fixation and ion beam tomographic imaging (**Fig. 5a**). To quantify relative spatial distribution, the correlation coefficient was determined by comparison of pixels in replication channel (^127^I) and the chromatin (^31^P) channel across 550 slices. For non-synchronized cells, the correlation coefficient was 0.32 for the replicated DNA and total nucleic acid but was negligible (correlation coefficient 0.03) for replicated DNA domains as compared to regions that reflect the presence of RNA (**Fig. 5c**). Newly synthesized transcripts were highly enriched in decondensed chromatin across 3D nuclear regions (**Supplementary Fig. 26**). The simultaneous RNA-DNA measurements in S-phase Nalm6 cells also exhibited the spatial disconnect (**Supplementary Figs. 27 and 28**). The longer incubation (2 h) increased the transcription signal but preserved the anticorrelated spatial patterns when compared to the shorter incubation (30 min) (**Fig. 5a and Supplementary Fig. 29**).

**Figure 5.**
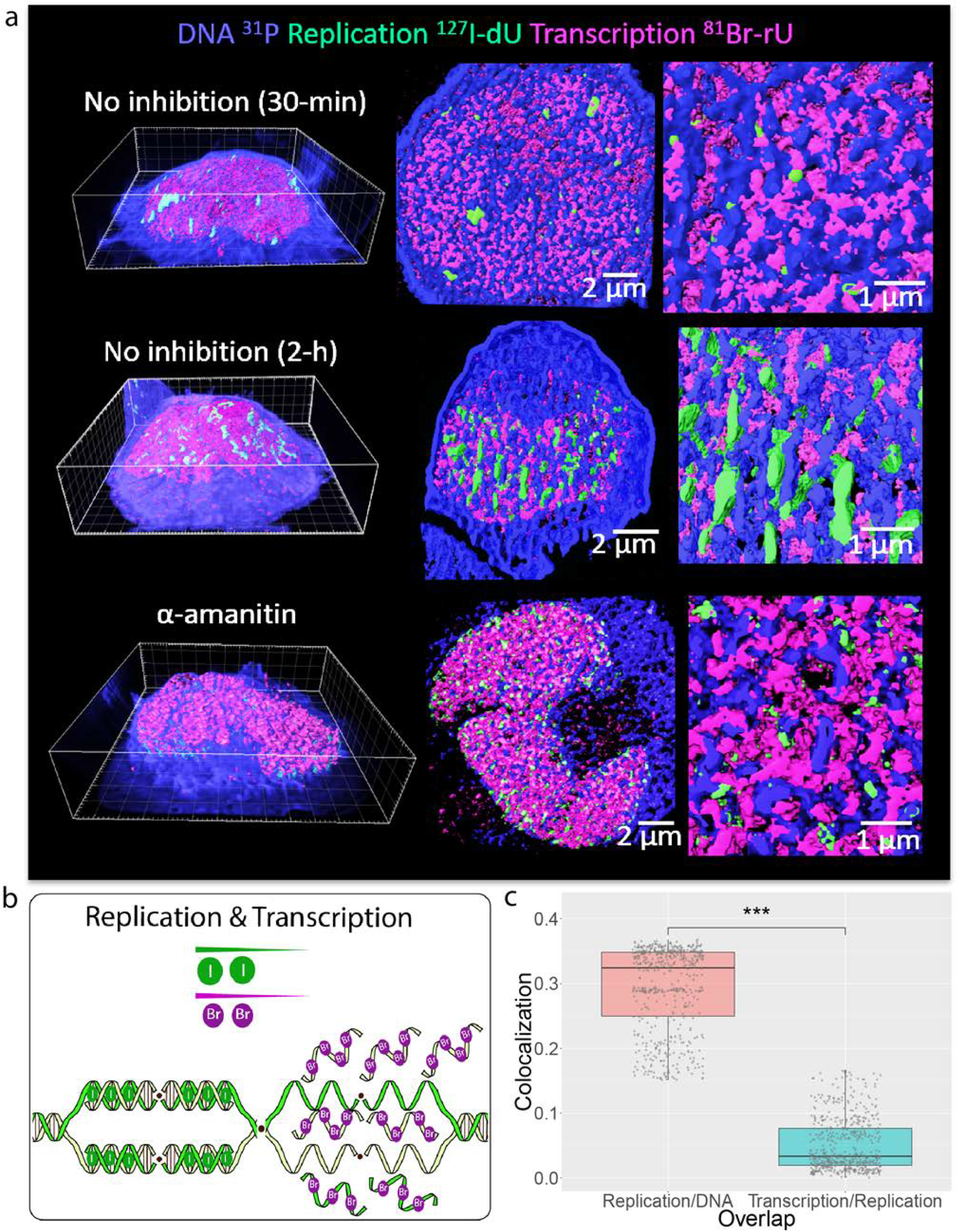
Spatial segregation of replication and transcription visualized by ion beam tomography. (a) Transcript synthesis was detected with ^81^Br-rU (magenta), and DNA replication was detected with ^127^I-dU (green) by simultaneous 30-minute and 2-hour incubation of Nalm6 cells with labels. The ^31^P channel (blue) reflects primarily nucleic acids. Top two row images are of cells not treated with drug and the lower images are of a cell treated with α-amanitin. The first image shows the 3D render. Middle and left images illustrate the spatial distribution of RNA transcripts (magenta) and replicating DNA (green) near the chromatin (blue). In the cell not treated with α-amanitin, RNAs were mostly localized around decondensed (medium chromatin compaction) regions, while the replication sites were closer to higher density chromatin regions. In the cell treated with α-amanitin, which inhibits transcription, the RNA was mostly observed in low density chromatin regions. (b) As shown schematically, our data indicate that replication and transcription machineries are spatially segregated. (c) Bar plot of co-localization of transcription and replication pixels. Each data point represents an ion beam depth section. The difference in the bar plots had significant P-value *(\*\*\**) by Wilcox test.

To validate our observations Nalm6 cells were pulsed with ^81^Br-rU for 30 minutes and then treated for 30 minutes with a transcription inhibitor α-amanitin^30^, and followed by fixation and ion beam imaging. During the incubation with α-amanitin, transcript synthesis was repressed primarily in the decondensed chromatin regions (**Supplementary Fig. 25**). The 3D renders and zoomed images from signals in the ^31^P and ^127^I-dU channels, both mostly reflective of DNA, validated the spatial segregation of replicated DNAs and newly synthesized RNAs near chromatin. To explore the efficacy of drug on active transcription, another cell that had been simultaneously treated with α-amanitin and ^81^Br-rU for 2 hours was fixed and imaged. Drug treatment significantly reduced transcription by 0.67-fold after cell volume normalization relative to a cell not treated with α-amanitin (**Supplementary Figs. 30 and 31**). To further validate the IBT approach, ^127^I-labeled rU was used to label transcripts and ^81^Br-dU was used to label replication forks in Nalm6 cells. Even at the level of individual pixels in raw ion images, spatial segregation of replicated DNAs and newly synthesized RNA was apparent in different depth sections (**Supplementary Fig. 32**).

#### Visualization of specific transcripts

The IBT method was used to visualize specific transcripts using the single-molecule FISH (smFISH) strategy.^31^ Instead of using fluorescent tags the oligonucleotide probes were conjugated to biotin and streptavidin-conjugated 5-nm ^197^Au nanoparticles were used to detect biotin. The nanoparticles allowed detection of single RNA molecules without the need for amplification. As a proof of concept, 24 biotinylated oligonucleotides of 20 nucleotides in length were designed to hybridize to the highly expressed *ActB* mRNA (**Supplementary Fig. 33a**). Use of multiple oligonucleotides that target the single mRNA result in significant signal enhancement relative to a single probe.

First, spin disk confocal fluorescence microscopy using streptavidin-conjugated Alexa 488 and an anti-Alexa 488 antibody was used to validate probe hybridization. *ActB* mRNA molecules were highly expressed in the cytoplasm of a HeLa cell and a few actively transcribing RNAs were also detected in the nucleus (**Supplementary Fig. 33b**). For IBT experiments, cells were incubated with ^127^I-dU for 23 hours, then RNA was labeled by incubation with ^81^Br-rU for 1 hour, followed by fixation, permeabilization, and smFISH. Newly synthesized RNAs were measured by analysis of incorporation of ^81^Br-rU, and *ActB* transcripts were detected by smFISH in the ^197^Au channel. In a HeLa cell, the 25-µm central portion was imaged across 450 depth sections. *ActB* mRNA was observed at depth levels ranging from 50 to 350 slices (**Supplementary Fig. 33c**). Ion beam tomographic RNA images were then visualized by 3D rendering (**Supplementary Fig. 33d**). In a Nalm6 cell (**Supplementary Fig. 34a**) and in a Jurkat cell (**Supplementary Fig. 34b**), ion beam images from the 30^th^ through the 100^th^ depths showed that newly transcribed RNAs were concentrated around decondensed chromatin and that *ActB* transcripts were mostly localized in cytoplasm around the periphery of the cells. Newly synthesized RNAs and *ActB* mRNAs minimally overlapped in the sections analyzed due to the very low concentration of *ActB* mRNA within the nuclear sub-volumes.

### Computational dissection of subcellular organization

Ion beam 3D subcellular organization was modeled by hierarchical clusters, correlation and association maps. Tomographic information cubes were represented as data frames with spatial coordinates (X, Y, Z) and ion pixel values. A systematic progression of spatial positions and association maps of replicated DNAs, newly synthesized RNA, phosphorous (DNAs, phosphorylated proteins and nucleic acids) and total proteins (^34^S as the base of amino acids) were studied in the ion beam tomograms from previously presented data.

First, Euclidian distances between pairs of ion beam voxels from each channel were calculated to determine the relative spatial positioning in two interacting ion sub-volumes for distinct ion channels. Clustering of these Euclidian distances for each pair-wise interaction in the ion images provided spatial proximity (**Fig. 6a**) of simultaneous incorporation of ^127^I-dU for replicated DNA (**Fig. 3a**) and ^81^Br-dU for another replicated DNA. Two large significant clusters were observed with high similarity (dark red) across the diagonal hierarchical maps. Since Thymidine-analogues were incorporated into the nearby locations within the chromatin, replicated DNA in both IdU and BrdU channels were organized at a high spatial proximity across subcellular volumetric partitions. Next, to study global organization matrices for voxel values of IdU, BrdU, and phosphorus (^31^P) of were converted to Pearson correlation coefficients and represented as 3 x 3 correlation maps (**Fig. 6b**). A high degree of correlation (dark blue) was obtained for the corresponding replicated DNA channels in IdU and BrdU, along with slightly reduced correlation of replicated DNA channels with phosphorous. An association map between the replicated DNAs showed wider circular coverage (corresponding to the higher association) in the Chord diagram owing to the similar spatial distributions of DNA pixels, while replicated DNAs showed reduced association with phosphorus due to the contaminants from additional nucleic acids and phosphorylated proteins in the that channel (**Fig. 6c**).

**Figure 6.**
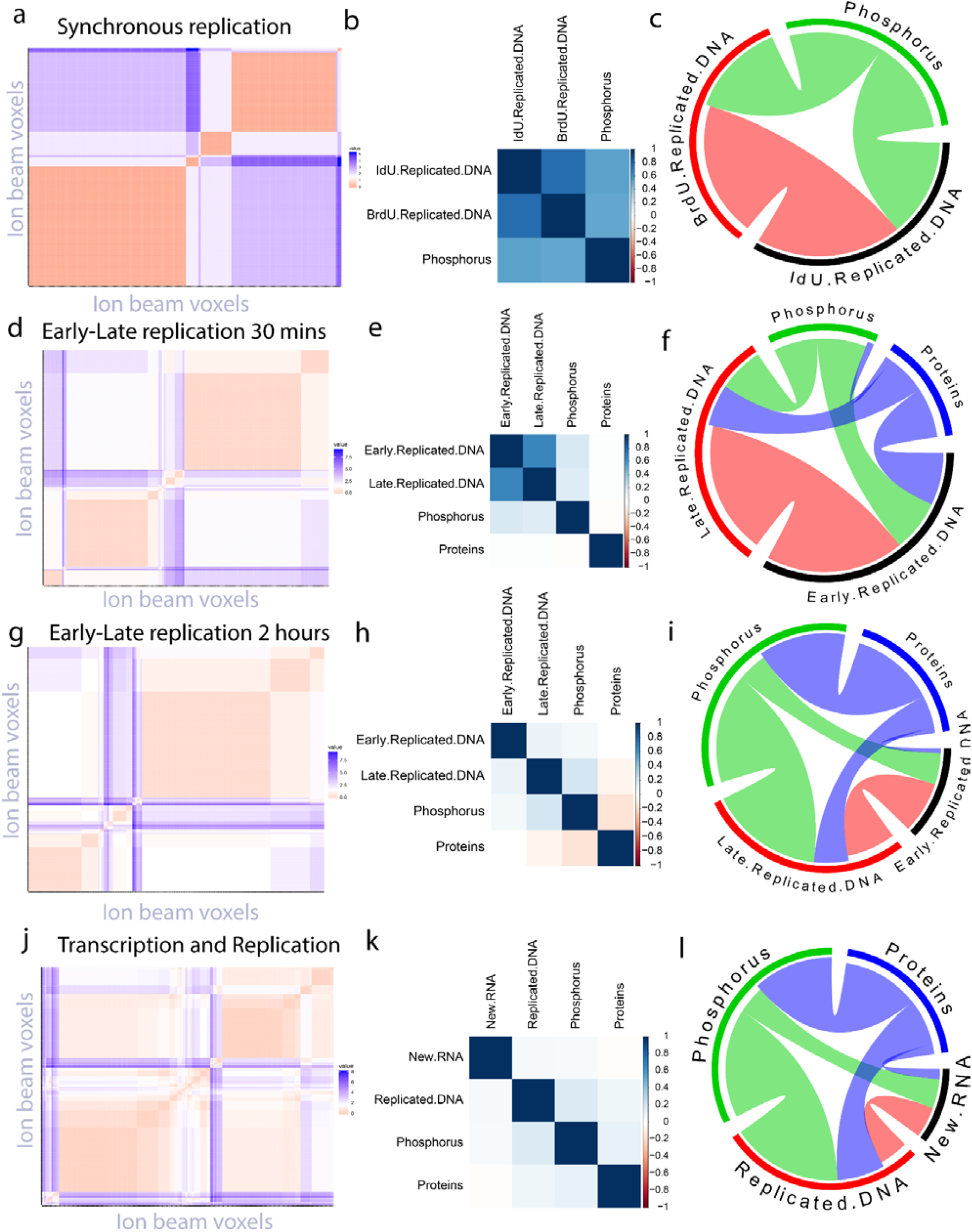
Hierarchical clusters, association and correlation maps revealed subcellular structural organization of replicated DNA, synthesized RNA, natural phosphorus and total proteins from ion beam tomographic information cubes. Correlation analysis in a HeLa cell with simultaneous incorporation of ^127^IdU and ^81^BrdU, presented in Figure 3a. (a) Euclidean distance based clustered data suggests high degree similarity (strong red color, two significant clusters) between IdU and BrdU images. (b) Correlation plot of replicated DNA in IdU and BrdU with phosphorus (^31^P) channel. IdU and BrdU voxels across 3D cell volume exhibited high correlation (dark blue). Phosphorous exhibited slightly lower correlation with IdU and BrdU. (c) Chord diagrams showed higher association of IdU with BrdU compared to the Phosphorus. Structural organization of early and late replication forks separated by 30 minutes chase in a Nalm6, shown in Supplementary Fig. 23. (d) Clustered data between 30-min separated IdU and BrdU showed slightly less correlation patterns (two significant clusters but smaller size) compared to the synchronous replication-based map, as expected. (e) Early and late replication sites remain correlated (lighter blue) with each other, while much less with phosphorus. Total proteins (^34^S) in the cell partially correlated with replication. (f) Direct association between IdU and BrdU (red) exhibited similar circular areas compared to the synchronous replication map in (a). Proteins (blue) interacted with replicated DNAs and partially with phosphorus. Similar hierarchical mapping of IdU and BrdU separated by 2 hours chase, demonstrated in Figure 4a. (g) Strong patterns (red) in clustered map reduced compared to the map from 30-min chase in (d) and blue values suggested higher dissimilarity. (h) Now the early-late replication forks are slightly correlated (very light blue) and decent correlation with phosphorous. Proteins were weakly correlated with replication and phosphorus signal. (i) Chord diagram showed consistent circular patterns with phosphorous (green), protein (blue) and early replicating DNA (black) and late replicating DNA (red). The association between early and late replicated DNA significantly reduced. Correlation maps of transcription and replication analysis (pulsed 2 hours) in a Nalm6, detailed in Figure 5a. (j) Multiple square patterns appeared in the clustered data from IdU (replication) and Br-rU (transcription), suggesting isolation of the two. (k) Correlation of newly synthesized RNA and replicated DNA is now almost negligible (faint blue). New RNAs are weakly correlated with phosphorus, whereas replicated DNA correlated with phosphorous signals. Protein exhibited considerable correlation with others. (l) A similar Chord diagram was also obtained as in (i). New RNAs have association with replicated DNAs (red) but less the previous association of early-late replicating DNAs in (i).

Second, the relative spatial distribution of early replicated DNA (IdU) and late replicated DNA (BrdU) that are separated by 30-minutes of cell division (**Supplementary Fig. 23**) was determined from the analysis of Euclidian distances between ion beam voxels from each ion channel. Hierarchical clustering of early replicated DNA and late replicated DNA revealed again two significant clusters with a smaller size (**Fig. 6d**) and lower Pearson coefficient values compared to the synchronous replication data (**Fig. 6e**). Global correlations (4×4 map) of early and late replicated DNA channels with phosphorus (^31^P) were lower due to the spatial enrichment of replicated DNAs in deconcentrated spatial regions of the original chromatin. Since decondensed chromatin exhibits lower phosphorus signal due to the lower original DNA concentrations in a flat cell, the correlations with replicated DNA would be expected to vary in this manner within cells. The correlation between the replicated DNAs and total proteins (^34^S) showed much lower values due to the spatial enrichment of dense proteins in the cytosol regions of the cell. In the association maps, early and late replicated DNA was associated the most with largest circular area, suggesting that the majority of the correlations are between these regions (**Fig. 6f**). Phosphorus and proteins were associated with considerable circular distributions due to the interplay with DNA-protein interactions in the nucleus.

Third, relative spatial modeling of early (IdU) and late (BrdU) replicated DNAs with 2-hours (**Fig. 4a**) delay were studied. Hierarchical clusters of Euclidian distances exhibited even smaller clusters (**Fig. 6g**) compared to both synchronous and 30-min delay replication data. A correlation map showed significantly lower values for early and late replicated DNAs and smaller values for phosphorus and total proteins (**Fig. 6h**). An association map exhibited a reduced circular area in the diagram (in other words, diminished correlation) between early and late replicated DNAs, together with similar associations of phosphorus and proteins (**Fig. 6i**).

Finally, the relative spatial positioning of replicated DNA (IdU) and newly synthesized RNA (Br-rU) after 2-hours incorporation (**Fig. 5a**) was explored. Clustering of Euclidian distances between the voxels of replicated DNAs and synthesized RNAs showed four distinct clusters (**Fig. 6j**). Two of these clusters were diagonal distributions for overlaps and the other two were in the top-left and bottom-right corners of the map. These support a spatial isolation model of replication and transcription machineries.^28,29^ Correlation maps showed minimal correlation values of replicated DNAs and new RNAs, while considerable relevance of phosphorous and proteins with replication and transcription (**Fig. 6k**). Replicated DNAs and RNAs covered isolated local spatial subvolumes, and at the same time, larger global volumetric portions (compared to those local regions) of the chromatin landscape exhibited concurrent emergence of replicated DNAs and synthesized RNAs in nearby positions. Association maps showed minimal relevance of replicated DNAs and new RNAs (**Fig. 6l**), and consistent connotations of phosphorous and proteins across previous diagrams (**Fig. 6 f, i, and l**).

Together, repeatability and consistency of association maps for unique cells with systematic metabolic labeling showed a hierarchical and comparative structural organization of replicated DNA, synthesized RNA, total proteins, and natural phosphorus signal. Addition of protein factors, transcriptional regulators, lipids, and natural “point of source” cell signatures can benefit from the presented computational modeling to decode subcellular patterns both in health and disease.

### IBT-based histology

To demonstrate IBT’s utility in primary tissue, chromatin and its interactions with other molecular distributions were evaluated in formalin-fixed paraffin-embedded (FFPE) 10-µm thick lymph node tissue section from a T cell lymphoblastic lymphoma patient. We detected ^31^P, ^34^S, ^37^Cl, ^14^N, and ^12^C in individual CD3^+^ T cells (identified by standard immunohistochemistry) at sub-100-nm ion beam resolution across 600 depth sections (**Supplementary Fig. 35a**). The ^31^P signal was located primarily in the nucleus, as expected, and the ^34^S ion signal was enriched in the cytosol. Other channels showed high signals mostly in the cytosol and extracellular regions due to the presence in the tissue of numerous proteins in the extracellular space. A 3D rendering is presented from two different viewing angles in **Supplementary Fig. 35b**.

Chromatin topology is known to be tightly regulated by interactions with proteins. Super-resolution microscopy has been deployed previously to visualize interactions of chromatin and protein clusters in individual cells from culture^32^. For pair-wise detection of chromatin interactions with protein in tissues, a molecular neighborhood analysis was performed on ^31^P (primarily chromatin) and ^34^S (proteins) images. To spatially partition subcellular regions, a 2 x 2-pixel region was segmented by the center-of-mass of the smaller regions and Delaunay triangulation was performed. Distinct spatial subregions showed enrichment of strong DNA-protein interactions in nucleus; protein-protein contacts were detected in the cytosolic and extracellular regions. The 2D neighborhood maps for the 50^th^ and the 300^th^ depth sections of the IBT images are shown in **Supplementary Fig. 35c**. 3D Delaunay tetrahedrons of subcellular volumes are shown in a 3D plot in **Supplementary Fig. 35d** and as a heatmap in **Supplementary Fig. 35e**. Such label-free neighborhood analyses at high resolution within FFPE samples will enable analyses of intracellular changes that occur upon cellular recognition events.

Multiplex histology has been limited to 2D spatial analysis.^5^ Volumetric analysis of tissue biopsy samples using IBT will enhance our understanding of cell-to-cell interactions in the context of super-resolved details of receptor and marker distributions.^33^ As a proof of principle for use of IBT in histology, we analyzed the interactions of immune cells (CD45^+^, label: ^169^Tm) and cancer cancers (CK19^+^, label: ^141^Pr) within a tumor sample from a patient with cholangiocellular carcinoma under oxygen bombardment (**Supplementary Methods**). A non-uniform 3D distribution of surface markers was observed (**Supplementary Fig. 36**). Polarization of surface-markers helps determine the potential capacity of cell from the ion beam snapshots. These 3D histology details show the promise of IBT for quantifying cell-to-cell interactions.

## SUMMARY

IBT is a mathematical and technical framework for analysis of three-dimensional MIBI. The method enables analysis of ion beam images acquired at multiple depths. In a typical IBT experiment, cells were initially imaged at 0.2-4 pA ion current over 1,000 axial scans. After binning of images, digital deconvolution was applied to the transformed ion image series using a hybrid deblurring algorithm and an ion beam current-dependent point-spread function. Three-dimensional processing was implemented by segmentation, mesh, and molecular neighborhoods. In cultured cells and tissues, IBT revealed three-dimensional volumetric distributions of genomic regions, RNA transcripts, and protein factors with 65-nm lateral and 5-nm axial resolution.

The presented IBT platform will open new avenues of subcellular imaging research to study aberrant cells:

- Direct visualization of 3D chromatin at sub-100-nm spatial details impacts study the role of nucleus in determination of cell function for dissecting epigenetic mechanisms.
- Subcellular 3D metabolic profiling for replication dynamics will find significant use in cell cycle regulation as a critical regular of therapeutic response. While not presented here, a systematic study of replication stress under various drug treatments in cancer models (human/mice) will shed light on subcellular dynamics in cultures and tissues.
- Nascent transcriptome plays a key role in cell function in cancers. Thus, the spatial visualization of relative DNA-RNA synthesis process will provide a subcellular signature in therapeutic effects.

When additional time points are evaluated and the strategy is combined with pseudotime techniques^1,34^, it should be possible to describe trajectories of changes in biological features at the single-cell level. The ion beam tomographic analysis pipeline described here can be used in analysis of data obtained using mass spectroscopy-based imaging technologies.^35,36^ In the future, 3D machine learning algorithms^37^ could be used to train and classify MIBI images of subcellular regions for cell-specific analyses. Deep learning-based IBT technology will eventually be used for the creation of an epigenetic atlas of cells^38^ that will be useful in both basic research and in the clinic.

## ACKNOWLEDGMENTS

We thank Stanford Nano Shared Facilities (SNSF), supported by the National Science Foundation under award ECCS-1542152. A.F.C. holds a Career Award at the Scientific Interface from Burroughs Wellcome Fund and National Institute of Health K25 Career Development Award (K25AI140783). X.R.-C. is supported by a long-term EMBO fellowship (ALTF 300-2017). This work was supported by NIH5R01NS08953304, NIH5U54CA14914505, Juno Therapeutics, Bill & Melinda Gates Foundation, Array BioPharma, NIH5UH2AR06767603, NIH5R25CA18099304, NIH5R01GM10983604, Department of the Army W81XWH-12-1-0591, W81XWH-14-1-0180, NIH5R01CA18496804, NIH5R01GM10983604, and the Rachford and Carlota A. Harris Endowed Professorship to G.P.N.

## AUTHOR CONTRIBUTIONS

A.F.C. designed the project and wrote the manuscript. G.H., S.J., S.Y.C., X.R.C., C.M.S., Y.B., and C.H. contributed to the methods. G. P.N. supervised the project and wrote the manuscript.

## ONLINE METHODS

### Synthetic ion beam image construction

Ion beam images were modeled based on linear image formation theory that is contaminated by a multiplicative noise and modified by the ion extraction efficiency:

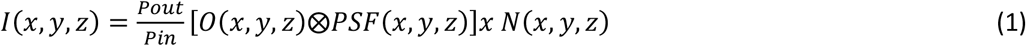

where *I(x,y,z)* is the detected image at the detector output, *O(x,y,z)* is the true object distribution that includes proteomic samples (individual proteins and cellular labels), *PSF(x,y,z)* is the point spread function of the ion beam imaging platform for each gun source and elemental labeling, *N(x,y,z)* is the multiplicative noise that is calculated from an ion image of interest, *pin* is the density of atoms impinging on the sample per unit time, and *Pout* is secondary ions extracted from the sample per unit time.

Each true object, *O(x, y, z)*, is an array of points distributed in space adjusted based on the required synthetic pattern. Initially, we assumed an array of proteomic signatures that were separated by a distance (*d*). The simulated true distribution was then convolved by *PSF(x,y,z)* of the ion beam imaging system. Typically, the x/y extent of the PSF is limited to ion beam width (50-500 nm) and the z extent is related to sample etch rate (1-30 nm). Using x/y:100, z:5 nm, we obtained a blurry image without noise. The convolved image was then contaminated by the additive and multiplicative noise factors from electronics and other factors. As background level is very low in ion beam imaging, we only considered multiplicative noise, *N(x, y, z)*, as a gamma distribution that was computed from a sample ion image. During the image acquisition, ions are lost during the secondary ion beam conversion and electronic read out. We included an additional multiplicative factor for ion extraction efficiency, *Pout/pin*, in the range of approximately 3%. Finally, the synthetic ion beam image, *I(x, y, z)*, was digitally sampled by a pixel size of x/y to be in the 50-100 nm range.

### Ion beam imaging experimental design

#### Ion beam source

Oxygen and cesium were used as the sources for NanoSIMS 50L (Cameca).

#### Acquisition

Ion beam current determines resolution and sensitivity. Lower current provided higher spatial resolution, whereas higher current in the cesium beam allowed deeper etching for faster 3D imaging (**Supplementary Fig. 14**). Lateral image resolution is governed by the width of the ion beam size for each source (Cs, O^-^/Au). The pixel size should be smaller than half of the highest spatial frequency of the ion beam image based on the Nyquist criterion. Thus, the pixel size should ideally be smaller than large portion of the beam width:

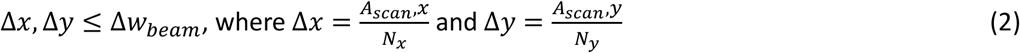

where Δ*x*, Δ*y* are pixel sizes, Δ*w*_*beam*_ is ion beam width, and *N*_*x*_, *N*_*y*_ are total number of pixels, and *A*_*scan*_, *x, A*_*scan*_, *y* are total ion image area. For instance, *N*_*x*_ *or N*_*y*_ = 256 x 256 for a cellular area of *A*_*scan x,y*_= 20 µm x 20 µm yields Δ*x*, Δ*y* =78 nm, a pixel size that performs well for Δw_*beam*_ = 100 nm. Resolution and pixel size comparisons were performed for the same cell (**Supplementary Figs. 9-10**). Much smaller pixel sizes are not optimal as the image acquisition time significantly increases. Dwell time per pixel must also be tuned. The typical value for the NanoSIMS is 1,000 ms. Better sensitivity was achieved around 2,000 ms at the cost of increasing acquisition time per pixel; however, ion beam images were distorted at dwell times longer than 2,000 ms.

#### Sample handling

Ion beam imaging requires conductive sample carriers, and thus, bare glass is not suitable. Custom-sized (7 mm x 7 mm) and (18 mm x 18 mm) silicon wafers (Silicon Valley Microelectronics) were used to host cells and tissues. Gold-coated substrates were not appropriate for long depth scans as the regions without cell materials were etched and caused charge buildup, which significantly distorted images.

#### Imaging settings

Samples were loaded into a NanoSIMS 50L (Cameca) device, and vacuum was set within the first 1-2 h. To adjust mass selection, 3 µL master mix buffer (^127^IdU and ^81^BrdU stock solutions) was dried on a large spot next to the cells and was imaged prior to cell or tissue imaging. After etching an elementally enriched region by high current ion beam sputtering, SIMS signal levels became detectable. Exact mass values were determined and set from the tuning window. Optics and detector settings were fine tuned to maximize the ion beam signal. Using optimized mass settings, cells were imaged at first high sputter rate (aperture setting D1_0 with >100 pA current) to remove outermost layers, followed by low current ion beam imaging (aperture setting D1_3 with >0.2-4 pA current) for high-resolution analysis. In this work, entrance slit and aperture slit values were kept at 0 for maximized ion signal. In the SIMS panel, elements need to be separated more than 3 atomic mass units. Based on the optimum parameters, a large ion beam scan was captured over a 50 µm x 50 µm area with 256 x 256 pixel window. A dwell time of 1,000 ms per pixel was used. Once a region of interest (ROI) was defined, a 20 µm x 20 µm subcellular area was scanned over 1,000 depths, typically taking 18 h of image acquisition per cell. Simultaneously data on seven ion channels (^12^C, ^12^C^14^N, ^19^F, ^34^S, ^31^P, ^81^Br, and ^127^I) and a secondary electron microscopy image were captured at each scan. The acquired images were visualized by an open source OpenMIMS plug in in Fiji and ImageJ. Batch analysis and 3D analysis are done by the presented analysis pipeline.

### Mathematical pipeline for ion beam tomography

#### Image conversion

Raw ion beam images were saved as .im format files, and a MATLAB function was used to convert .im files to TIFF files.

#### Registration

Image alignment is required for different depth scans as the ion beam drifts during the acquisition. To address this problem, a cellular subregion that stayed almost stationary across all the depths was selected, and depth images were spatially registered down to pixel levels. Additional fiducial markers were included to perform the registration. One approach was to coat microparticles such as iron (^55^Fe) microbeads (2-6 µm diameter) onto the cell sample and visualize the bead along with the subcellular ROI (e.g., entire nucleus) in the electron microscopy channel or specific elemental image (**Supplementary Fig. 12**).

#### Multi-slice processing

Raw ion beam images were summed over a few depth image sections (1 to *n*, where *n* varies between 3-20 slices) to improve the signal-to-noise ratio (SNR) of the combined ion beam images:

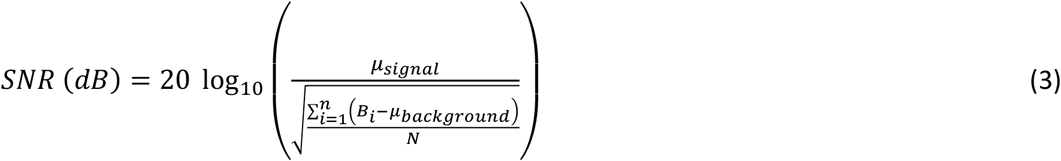

where *μ*_*signal*_ is the mean signal value and *μ*_*background*_ is the mean background value, 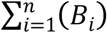 is the background pixel value, and *N* is the number of pixels in the background region of the ion beam cellular image.

Summing up to 10 subsequent slices increased SNR exponentially (up to SNR=7 dB), whereas SNR slightly increased after binning more than 10 slices (up to SNR=11 dB). Thus, subgrouping a large depth scan into *n*=10 slice groups provided optimum mathematical IBT performance. Based on the quality of raw image scans, the *n* was increased for dim ion signal and was decreased for bright ion images. To avoid axial resolution loss, we binned 10 slices and incremented by one slice at a time over the entire section. Thus, IBT preserves original depth resolving power down to 5-10 nm axial resolution.

#### Digital confocality

A 3D image enhancement algorithm was applied to the summed images for digital confocality on ion beam imaging. The Lucy-Richardson (LR) deconvolution algorithm was used to deblurr the image iteratively based on the modification of the PSF at each iteration:

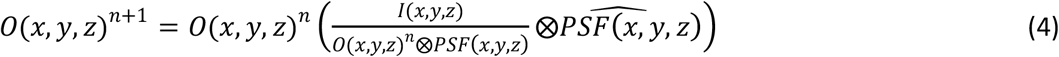

where *O(x, y, z)* ^*n*^ and *O(x, y, z)* ^*n*+1^ are the object distributions in *n*^*th*^ and *n+1*^*th*^ iterations, *I(x,y,z)* is the original ion beam image, ⨂ is the convolution operation, *PSF(x,y,z)* is the original PSF, and 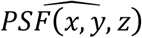 is the modified PSF. The algorithm outputs *O(x, y, z)* ^*n*+1^ ion image after *n+1* iterations. After setting the image standards for each channel and element, the iteration number was fixed between 3 and 10 to provide sharper ion beam images.

Here, we performed a hybrid deconvolution, a combination of iterative LR algorithm and a blind PSF. Based on the measured current levels of the ion beam imaging (0.2 to 4 pA), a PSF width was computed and included in the LR deconvolution algorithm. The deconvolved images were then lightly (2 x 2) filtered by a Gaussian function or median filter to reduce the image noise. The resultant images were passed to the 3D quantification and representation.

#### 3D visualization

Deconvolution results were processed to explore the 3D distribution of subcellular regions in individual cells. Here we used the *Imaris* module of Bitplane to handle IBT data sets. Using this 3D rendering pipeline, 3D movies were created of transcription, replication, and chromatin experiments in individual mammalian cells (**Supplementary movies 1-18**). Next, images were segmented in 3D and connected objects were labeled. For quantitative comparisons of specific subcellular distributions, centroids and volumes of each sub-object were listed as an output excel text file or MATLAB variable, followed by statistical analysis in R studio. Finally, to define cell-specific subcellular distribution, a mesh algorithm on image voxels was developed: Octrees subdivides a three-dimensional cellular space by iteratively partitioning it into eight octants. Instead of directly comparing voxel-to-voxels, a data structure was generated from Octrees that provides an identifier of a cell state from subcellular 3D labels (**Supplementary Fig. 16**).

## Analysis

### Image co-localization

Two separate ion channels exhibited overlaps or spatial segregation for transcription and replication studies. Co-localization of ion channels at each depth was quantified based on the 2D correlation coefficient:

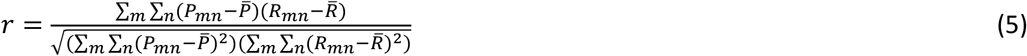

where *p*_*mn*_ and *R*_*mn*_ are two overlapping images, 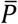 and 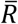 are the mean values of total pixels in each image, and *r* is the correlation coefficient. Higher *r* values suggested more overlap across all the pixels. Linear fit analysis for the entire 600-1,000 ion image slices generated a *r* distribution over 3D cell, which was then represented as box plots with scatter points.

#### Resolution

Spatial resolution was quantified based on the line scans. The signal drop from 84% to 16% of the maximum determined image resolution. The 84%-16% criterion provided 65-nm resolution for images acquired with the cesium source, whereas 395-nm resolution was achieved for experiments with the oxygen source. The axial resolution was high as each layer was etched only a few nanometers after interacting with the ion beam. Acquiring 1,000 depths across the more than half of the B cell showed less than 5-10 nm axial resolution as the cell size is around 10 µm. The 84%-16% criterion on line scan across depth profile also provided 5-nm axial resolution.

#### Sensitivity

Elements exhibit distinct sensitivities in ion beam tomographic analysis (**Supplementary Fig. 15**). High to low sensitivity of elements imaged by a cesium beam are S, Cl, Se, Br, I, F, Te, Au, Pt, O, C, Si, Ir, Ge, P, Ag, Sn, Os, and Sb. High to low sensitivity of elements imaged by an oxygen beam are Cs, Na, K, Li, Rb, Ca, Sc, Sr, Ga, In, Ba, Nd, Eu, Y, Pr, Al, Dy, Tb, Ho, Yb, Ce, Sm, Er, Mg, La, Gd, Tm, Lu, Ti, V, Zr, Tl, U, Cr, Nb, Mn, Mo, Th, Hf, Fe, Rh, Sn, Cu, Be, Ru, Ni, Si, Ag, Co, Ta, B, Pb, W, Pd, and Bi. The ratio of sensitivity levels was corrected for accurate mapping of true molecular concentrations in ion beam imaging experiments.

#### Molecular neighbourhood

The Delaunay triangles and tetrahedrons were extracted from IBT images after calculating center of masses in 2 x 2 pixels, 4 x 4 pixels, or 8 x 8 pixels subregions. The total ion image signal per area and centroids of triangles/tetrahedrons were extracted from MATLAB analysis. Heatmaps were then prepared in the R studio with *ggplot2* and *heatmap.2* packages. Ion signal levels of each corner and spatial positions were clustered together to spatially visualize subregions of pair-wise molecular interactions in chromatin.

#### Fuzzy logic segmentation

Chromatin states were determined based on grouping of pixel intensity values in the ion beam data. Fuzzy C-means (FCM) clustering was used to partition the volumetric ion beam data. The classified groups of clustered pixel values were than used to re-color the segmented cells in Fig. 3. Adjusting cluster dimension (C) allowed segmentation of fine chromatin sub-states. Tomographic ion images benefited from rapid identification of distinct density levels in the subcellular regions. K-means and other clustering methods can potentially provide similar segmentation results in subcellular tomographic images.

#### Similarity quantification

The structural similarity index (SSIM) is a quality metric (between 0 to 1) used to determine visual similarities of one image to another based on luminance (I: Intensity), contrast (C: Max/Min pixel intensity difference), and structure (S: features) of an image. SSIM was calculated as follows:

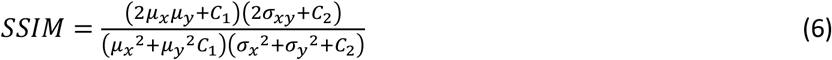

where *μ*_*x*_, *μ*_*y*_ are local means, *σ*_*x*_, *σ*_*y*_ are standard deviations, and *σ*_*xy*_ is cross-variance of the images.

In the MATLAB *ssim (A, ref)* function, the image of interest from the ^127^I-dU channel was compared to a reference from the ^81^Br-dU channel to determine similarity of spatial chromatin variations. To circumvent dynamic range issues in the raw and reconstructed DNA images, ion images were adjusted to [0, 1] range. These normalized images were then subjected to the SSIM quantification.

### Clustering, correlation and association maps

To explore relative spatial positioning across newly synthesized DNA and RNA, pair-wise distances were converted to Euclidian distances 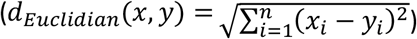 and the resultant distance matrix was clustered to determine the subcellular organization (**Fig. 6 a, d, g, and j**). To calculate global correlation patterns across the entire cell volume, 3D ion tomographic information cube was organized as a matrix array (X, Y, Z, ion voxel value), followed by calculation of Pearson correlation coefficients among replication and transcription data (**Fig. 6 b, e, h, and k**). To visualize association of each subcellular component, chord diagrams were created from the correlation matrix from the previous step (**Fig. 6 j, f, i, and l**).

### Cell culture

HeLa cells were cultured in Dulbecco’s Modified Eagle’s medium (DMEM 1×; Life Technologies) with 10% fetal bovine serum (FBS, Omega Scientific) and penicillin-streptomycin (Life Technologies) at 37 °C, 5% CO_2_ and passaged every few days with TrypLE Express (Thermo Fisher). Prior to imaging, cells were cultured on 18 mm x 18 mm or 7 mm x 7 mm silicon substrates overnight. For metabolic experiments, cells were incubated with the isotope of interest for the time indicated. After 24 h, cells on silicon substrates were washed with phosphate-buffered saline (PBS), followed by fixation with 1.6% paraformaldehyde (PFA; Electron Microscopy Sciences) for 10 min. Cells were then washed with PBS and deionized (DI) water a few times and then treated with ethanol. Samples were then dried in a vacuum desiccator for a few days before loading into the SIMS device for experiments. Jurkat cells were grown in RPMI 1640 (Life Technologies), 10% FBS (Omega Scientific), and 2 mM L-glutamine (Life Technologies). NALM6 clone G5 (ATCC, CRL-3273) cells were grown in RPMI 1640 (Life Technologies) and 10% FBS (Omega Scientific). Non-adherent Jurkat and Nalm6 cells were coated onto poly-L-lysine (Sigma-Aldrich) modified silicon substrates by spin coating at 30 RCF for 5 mins, followed by fixation with 1.6% PFA for 10 min. Cells were either permeabilized for antibody staining or directly washed with PBS and DI water for SIMS experiments.

### Pulse-chase metabolic experiments

HeLa cells on silicon substrates were cultured overnight. For chromatin tracing, cells were incubated in media with 10 μM ^127^I-dU (Sigma-Aldrich) and ^81^Br-dU (BioLegend) for 24 h. For replication studies, HeLa cells were pulsed for 30 min with 20 μM ^127^I-dU. After washing the cells twice with warm PBS, cells were incubated for either 30 min or 2 h in medium without label. Cells were then incubated for 30 min with 20 μM ^81^Br-dU. Jurkat and Nalm6 cells were pulsed and chased in tubes, followed by centrifugation at 125 RCF for 3 min to exchange buffers. Cells were then coated onto poly-L-lysine-modified silicon substrates and prepared for SIMS as above. For transcription studies, cells were pulsed with 2 mM ^81^Br-rU (Sigma-Aldrich) or 2 mM ^127^I-labeled rU (Sigma-Aldrich) for 30-60 min, followed by washes with PBS and DI water, and PFA fixation. To inhibit transcription, cells were treated with 10 µg/mL α-amanitin (Abcam) for 30 min on ice, followed by washes and PFA fixation. To synchronize the cells in S phase, cells were treated with 5 µg/mL aphidicolin (Sigma-Aldrich) for 16 h, followed by two washes with warm PBS.

### RNA detection

Twenty-four DNA probes (20-mers of unique sequence) were designed against different regions of *ActB* mRNA in Stellaris Probe Designer (LGC Biosearch Technologies). Each oligonucleotide was biotinylated at the 5’ end during synthesis (Integrated DNA Technologies), and all were pooled. Endogenous biotin and avidin binding sites in cells were blocked to avoid non-specific binding (Abcam, cat # ab64212). Cells were incubated with 2 nM oligonucleotide mixture in a buffer containing 2×SSC, 10% dextran sulfate, and 30% formamide at room temperature. After overnight incubation, the cell sample was washed with 30% formamide for 20 min and twice with 2×SSC for 5 min. Samples were then incubated with streptavidin-conjugated 5-nm gold nanoparticles (Cytodiagnostics, cat# AC-5-04-05) in a PBS1× buffer for 1 h at room temperature. After washing the cells with DI water, the sample was dehydrated and imaged by IBT.

### Confocal microscopy

Cells were prepared on coverslips (22 mm x 22 mm) for fluorescence experiments. Labeling and culturing conditions were the same as for IBT protocols. A spinning disk microscope (Andor DragonFly) was used to capture control fluorescence images of cells. A 60× objective lens with 1.6× tube magnification was used, providing total 96× magnification. Streptavidin-conjugated Alexa 488 (Thermo Fischer, cat # S32354) were used at 1:100-1:500 dilutions to target antigens for control experiments in cells. Power levels were adjusted to 50% of 488 laser. Micromanager interface was used to capture images. ImageJ was used to merge images for pseudocoloring.

### Code availability

Custom MATLAB and R source codes (**Supplementary Software**) with test images are available at https://github.com/nolanlab.

