## Supplemental Figures for "Ion beam subcellular tomography"

| Supplementary File | Title |
|---|---|
| **Supplementary Figure 1** | Ion beam tomography pipeline |
| **Supplementary Figure 2** | Synthetic ion beam imaging and microscopy data |
| **Supplementary Figure 3** | Lateral resolution analysis of ion images captured by Cesium gun source for Au coated Si samples |
| **Supplementary Figure 4** | Lateral resolution analysis of ion images captured by Cesium gun source for Jurkat cell samples |
| **Supplementary Figure 5** | Resolution analysis of ion beam images captured by Oxygen gun source for Al coated Si samples |
| **Supplementary Figure 6** | Axial resolution analysis by a circular replication site |
| **Supplementary Figure 7** | Axial changes of sixty slices for subcellular region |
| **Supplementary Figure 8** | Axial resolution by cell height divided by total scans |
| **Supplementary Figure 9** | Modeling of ion beam imaging parameters |
| **Supplementary Figure 10** | Pixel size effect of ion beam acquisition on analysis |
| **Supplementary Figure 11** | Signal to noise ratio analysis on multi-slice sums |
| **Supplementary Figure 12** | Registration patterns for IBT |
| **Supplementary Figure 13** | Rapid depth etching by high current levels |
| **Supplementary Figure 14** | Schematic for depth imaging by IBT |
| **Supplementary Figure 15** | Elemental sensitivity analysis for Cesium and Oxygen |
| **Supplementary Figure 16** | Voxel and octree analysis of replication forks |
| **Supplementary Figure 17** | IBT results for chromatin images |
| **Supplementary Figure 18** | Structural similarity index values for DNA images |
| **Supplementary Figure 19** | Similarity of DNA images in another Nalm6 cell |
| **Supplementary Figure 20** | Fuzzy logic segmentation in a B cell lymphoblast |
| **Supplementary Figure 21** | Fuzzy logic segmentation in a HeLa cell |
| **Supplementary Figure 22** | Four channel replication experiment (2-hours) |
| **Supplementary Figure 23** | Repeat of replication dynamics experiment (30-min) |
| **Supplementary Figure 24** | Repeat of replication dynamics experiment (2-hours) |
| **Supplementary Figure 25** | Statistics of overlaps in replication experiments |
| **Supplementary Figure 26** | Newly synthesized transcript localization |
| **Supplementary Figure 27** | Replicate for transcription and replication in Nalm6 |
| **Supplementary Figure 28** | Transcription and replication in a Nalm6 cell |
| **Supplementary Figure 29** | Long pulse (2-h) for transcription and replication |
| **Supplementary Figure 30** | Transcription inhibitors during 2-h metabolic labeling |
| **Supplementary Figure 31** | Transcription signal reduction by α-amanitin |
| **Supplementary Figure 32** | Transcription and replication (color swap) in Nalm6 |
| **Supplementary Figure 33** | 3D mature transcription in HeLa by RNA FISH labeling |
| **Supplementary Figure 34** | Mature and nascent transcription in Jurkat and Nalm6 |

| | |
|---|---|
| **Supplementary Figure 35** | Label-free mass imaging in FFPE tissues |
| **Supplementary Figure 36** | Volumetric histology by IBT |
| **Supplementary Methods** | |

**Computational pipeline for Ion beam subcellular tomography**

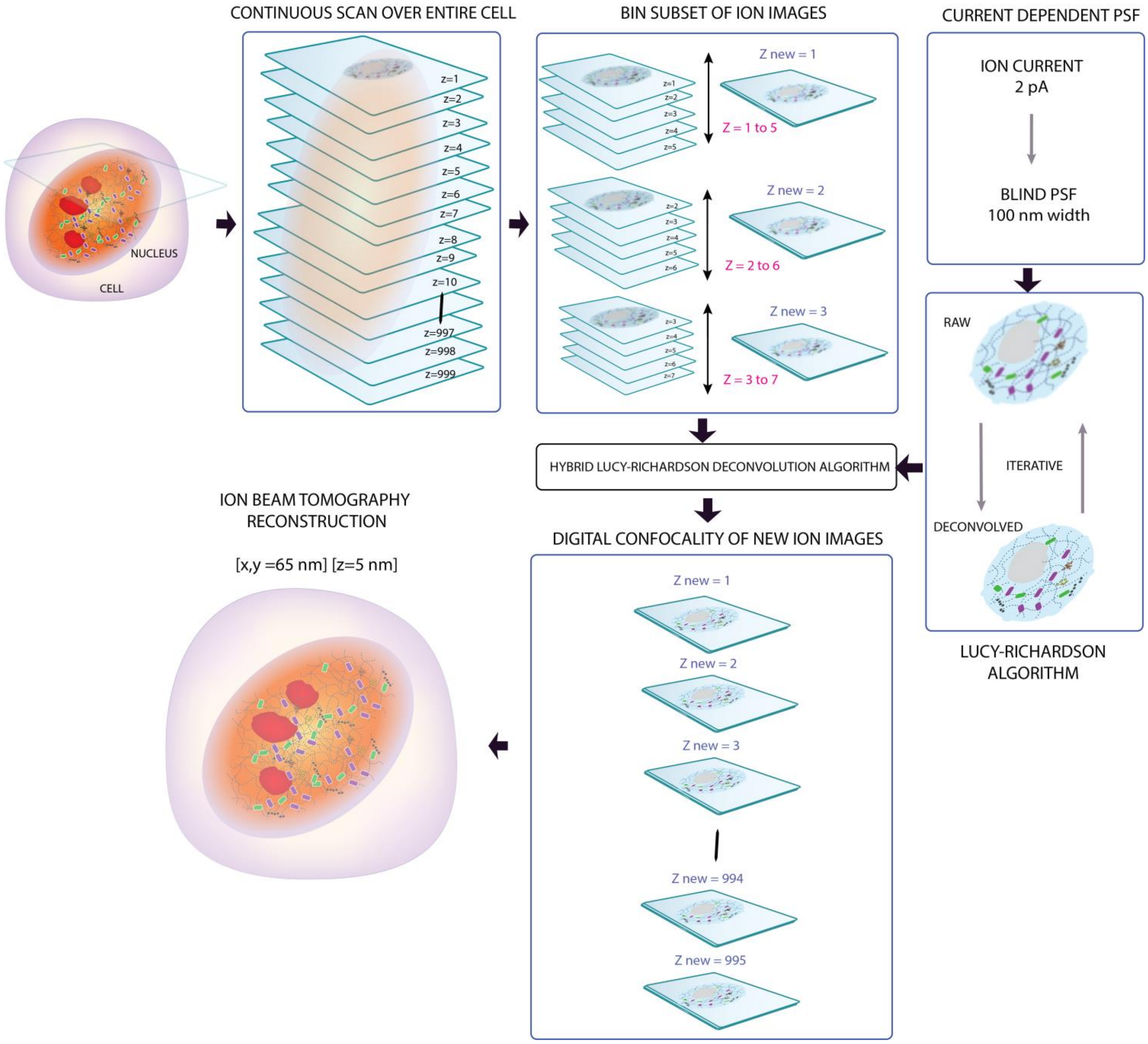

**Supplementary Figure 1.** IBT mathematical pipeline.

Ion images are acquired across 1,000 depth sections.

A subset of raw ion scans is binned (in this example, over five slices) to generate a new ion image with high signal to noise ratio (SNR) and less noise. This process is incremented by one slice at a time, providing a new ion image series with a similar dimension. The last few slices are skipped due to windowing.

A current-dependent point spread function is calculated for the deconvolution process.

Each ion image is then deblurred by an iterative Lucy Richardson deconvolution algorithm.

Five iterations were optimum for typical ion images but must be optimized based on SNR of the images.

The resultant sharper ion images are then stitched over axial direction for error-free reconstructions.

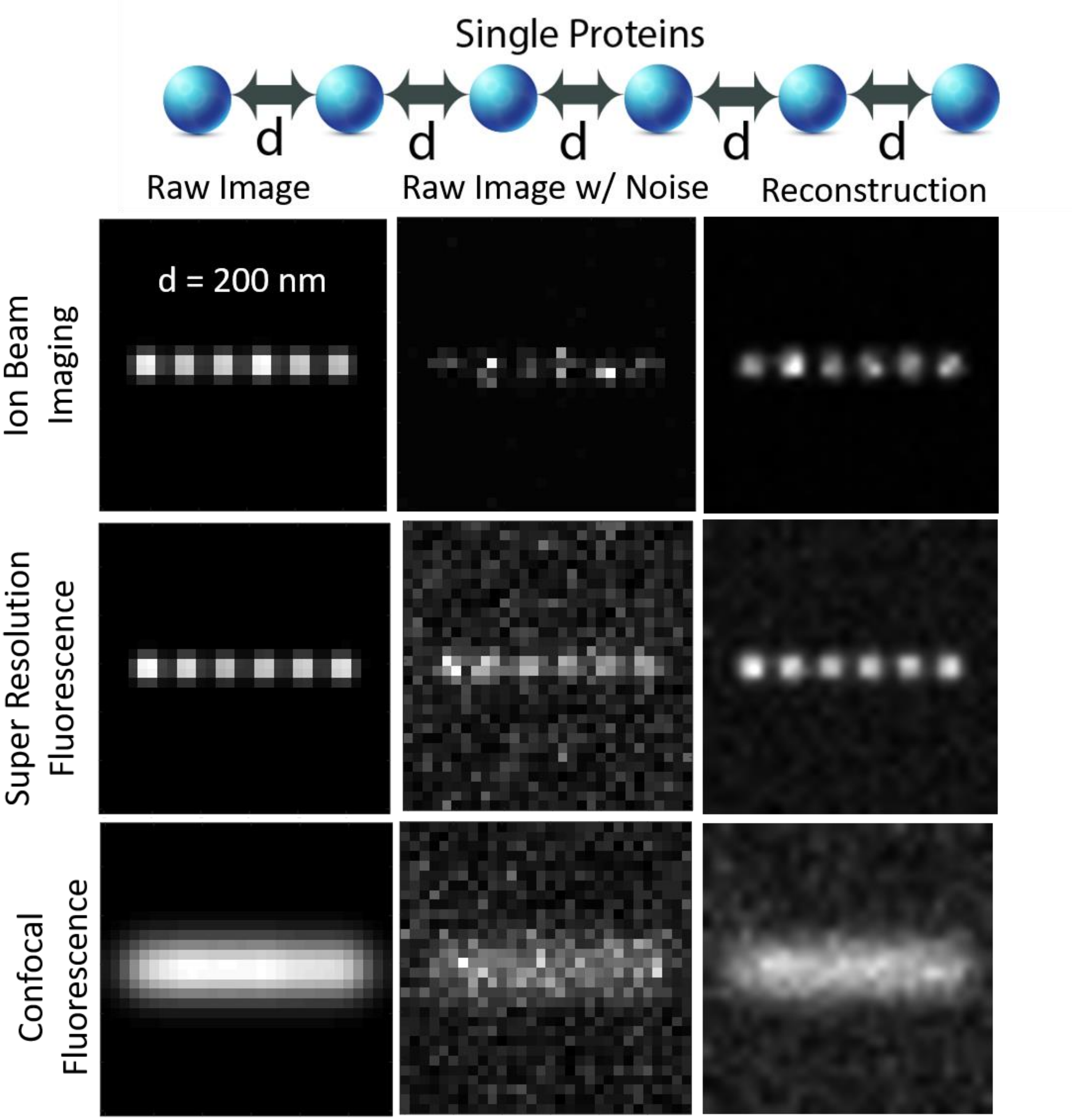
Single Proteins
d
d
d
d
d
Raw Image
Raw Image w/ Noise
Reconstruction
d = 200 nm
Ion Beam Imaging
Super Resolution Fluorescence
Confocal Fluorescence

**Supplementary Figure 2.** A simulated array used to evaluate the ion beam analysis pipeline.

Six protein signatures with a separation of 200 nm were generated for

Ion beam imaging (PSF: 120 nm; pixel size: 50 nm)
Super-resolution fluorescence imaging (PSF: 50 nm; pixel size: 50 nm)
Confocal fluorescence microscopy (PSF: 250 nm; pixel size: 100 nm).

Super resolution was assumed to be a STORM imaging platform for a computational PSF of 50 nm after reconstructions (real PSF size and localization process are not detailed here).

Noise was then measured from experimental images and used to contaminate each synthetic image.

Ion beam images have higher noise variation than fluorescence images.

The IBT mathematical pipeline was then validated after reconstruction of the noisy ion beam and fluorescence images.

A

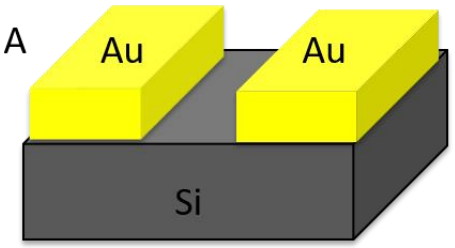


C

| | D1_1 | | D1_2 | | D1_3 | |
|---|---|---|---|---|---|---|
| | **Math** | **Raw** | **Math** | **Raw** | **Math** | **Raw** |
| Set | 689 | 731 | 96 | 244 | 68 nm | 215 |

Set

B

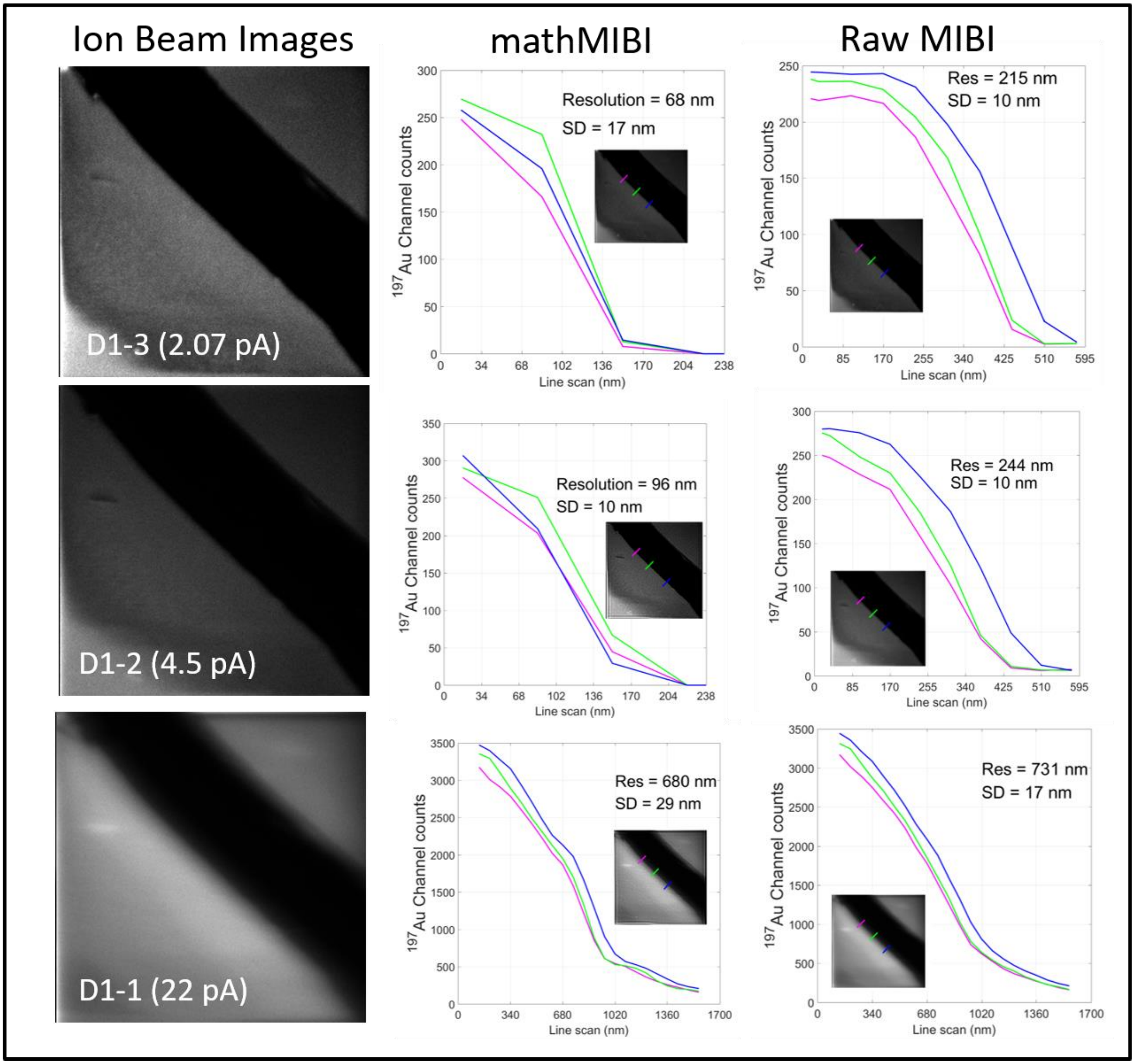

**Supplementary Figure 3.** IBT mathematical framework results in down to 68-nm lateral resolution.

(a) Au-coated silicon substrates were used to quantify spatial resolution in ion beam images generated by a cesium source.

(b) Ion beam images were acquired at 2.07, 4.5, and 22 pA using D1-3, D1-2, and D1-1 apertures, respectively. Three cross-sections at the edge of Au-Si were used to quantify spatial resolution. Insets show processed ion images with blue, green, and magenta line plots for 2.07, 4.5, and 22 pA, respectively.

(c) Summary of resolution limits of raw MIBI and reconstructed IBT images by mathematical analysis of MIBI (mathMIBI).

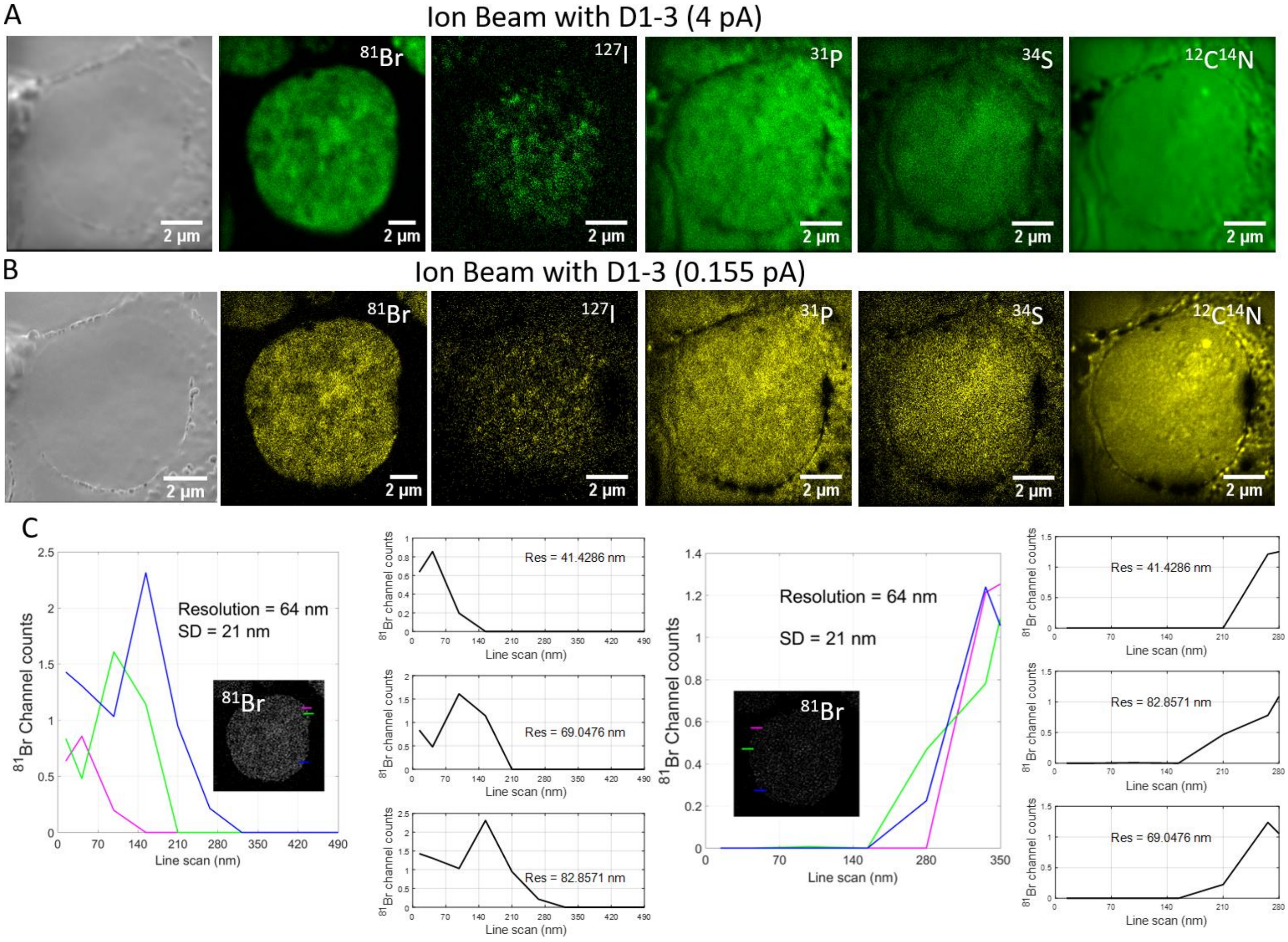
A
Ion Beam with D1-3 (4 pA)
81Br
127I
31P
34S
12C14N
2 µm
B
Ion Beam with D1-3 (0.155 pA)
81Br
127I
31P
34S
12C14N
2 µm
C
81Br Channel counts
Line scan (nm)
Resolution = 64 nm
SD = 21 nm
81Br
81Br channel counts
Res = 41.4286 nm
Res = 69.0476 nm
Res = 82.8571 nm

**Supplementary Figure 4.** Six-color ion beam imaging experiment for endogenous elements ($^{12}C$, $^{14}N$, $^{34}S$: total proteins; $^{31}P$: DNA backbone) and metabolite labels and replication loci (labeled with $^{127}I$-dU for 24 hours and $^{81}Br$-dU for 30 minutes) in Jurkat cells.

(a) Seven channel ion beam images of a Jurkat cell by D1_3 setting corresponding to 4 pA of ion current.

(b) Same cell was imaged at 0.155 pA current, highest resolution setting with smallest feasible ion beam width.

(c) Three distinct line scans were measured across right and left edges of the image detected at the $^{81}Br$ channel. Blue, red and magenta colors correspond to the line scan positions on the cell image and cross section plots. In both edges, the distance through which the %84 ion signal drops to 16% of the maximum yielded 65 nm±21 nm (STD) spatial resolution, agreeing well with the gold edge on silicon substrate experiment shown in Supplementary Fig. 3.

A

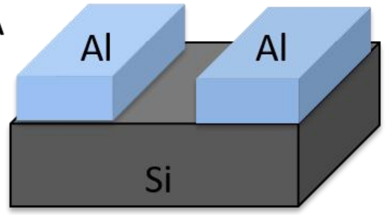


C

| | D1_2-159pA | | D1_3-71pA | | D1_3-8pA | |
|---|---|---|---|---|---|---|
| | Math | Raw | Math | Raw | Math | Raw |
| Set | 907 | 913 | 613 | 621 | 395 nm | 411 |

B

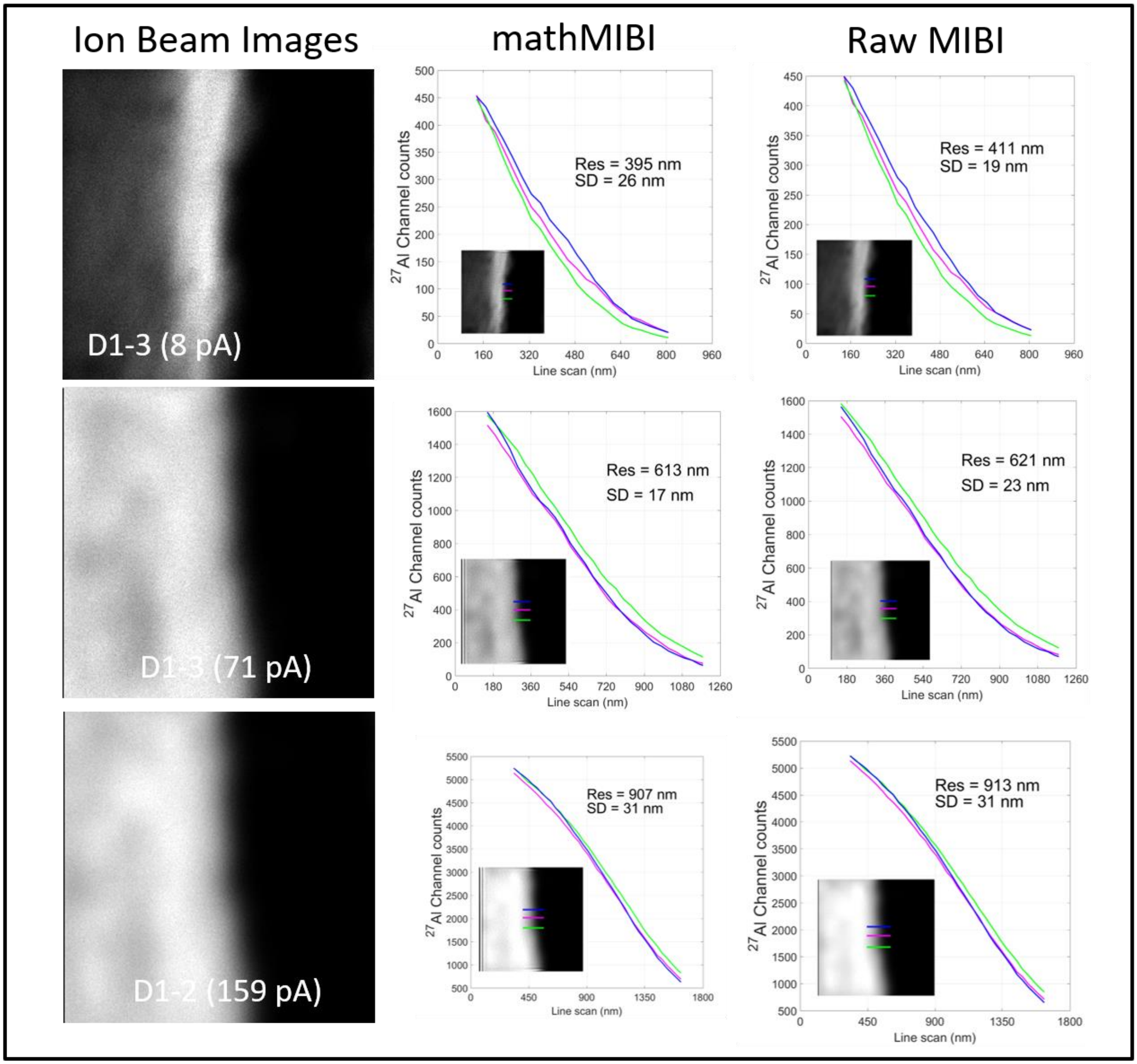

**Supplementary Figure 5.** Resolution characterization for ion images acquired using an oxygen source using deconvolution and analysis parameters similar to those described in Supplementary Fig. 2.

(a) An Al-coated silicon substrate was used to measure the edge profile.

(b) Ion beam images were acquired at 8, 71, and 159 pA using D1-3, D1-2, and D1-1 apertures, respectively. Three cross-sections at the edge of Al-Si were used to quantify spatial resolution. Insets show processed ion images with blue, green, and magenta line plots for 8, 71, and 159 pA, respectively.

(c) Summary of resolution limits of raw MIBI and reconstructed IBT images by mathematical analysis of MIBI data (mathMIBI).

Spatial resolution levels down to 395 nm were achieved. Larger iteration numbers will reduce these resolution levels while image artifacts may increase as a result of the processing. Due to the large beam width, computation faces challenges to decouple overlapping ion signals in the digital processing. Had the edge of this test substrate been sharper more stringent resolution criteria could have been obtained. Oxygen beam experiments are not the focus of this paper, but the IBT algorithm works on these images with decent resolution enhancements. Sub-500 nm resolution levels are sufficient for study of tumor samples obtained from biopsies.

**Axial resolution determination by a single replication fork**

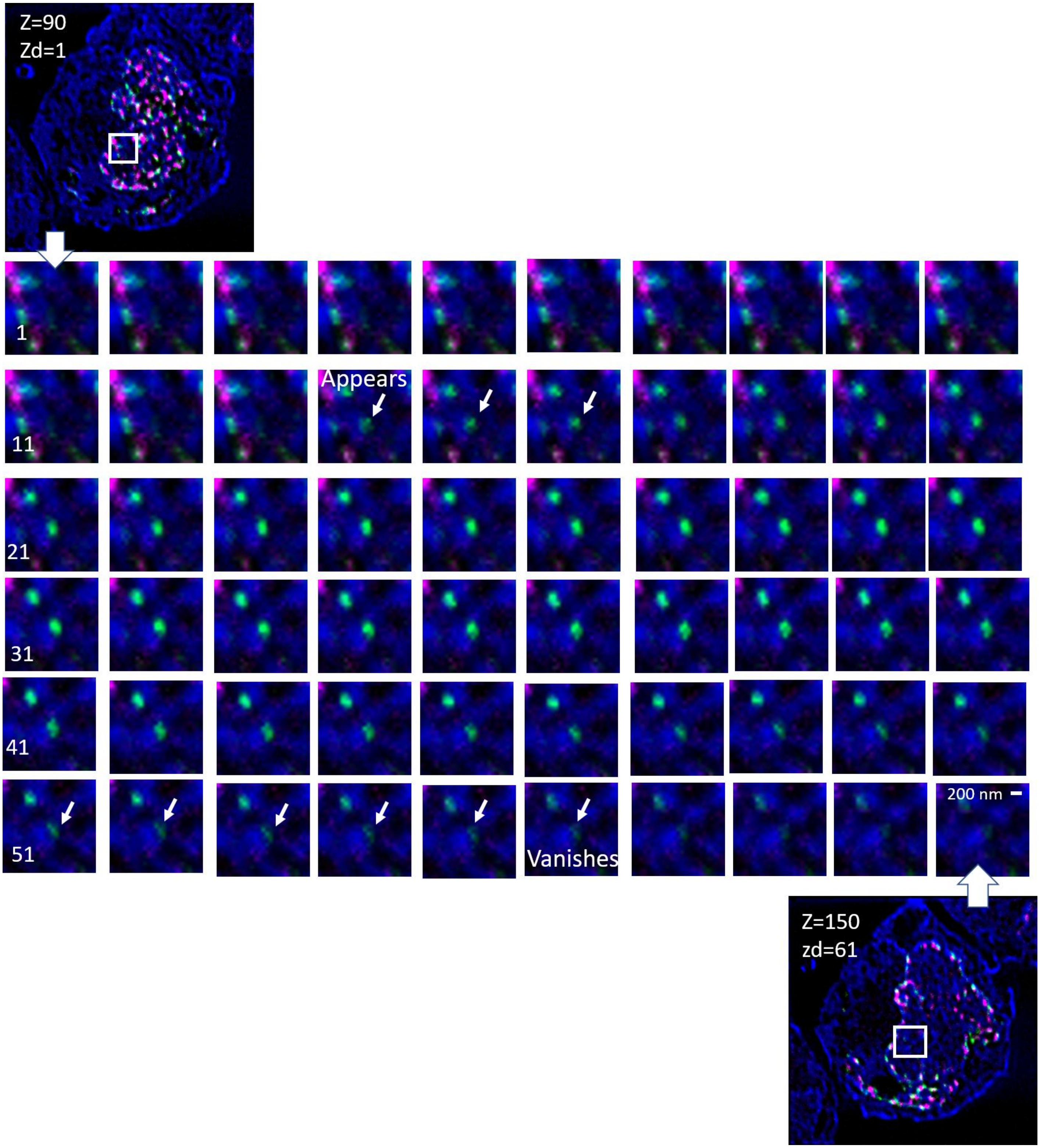

**Supplementary Figure 6.** Estimation of axial resolution by replication fork imaging.

$^{31}P$ channel (largely chromatin, referred as DNA) in blue, $^{127}I$-dU was a replication site in green, and $^{81}Br$-dU was another replication site in magenta. Each replication was a result of 30 minutes incorporation.

Sixty slices of two replication forks in $^{127}I$-dU channel reveal a 3-pixel replication site (size=175 nm in one direction) appearing around the 15$^{th}$ slice and vanishing around 55$^{th}$ slice.

From this image, the axial resolution was estimated to be approximately 5 nm as the 175-nm replication site extends over 40 slices.

**High resolution depth imaging of replication forks by ion beam tomography**

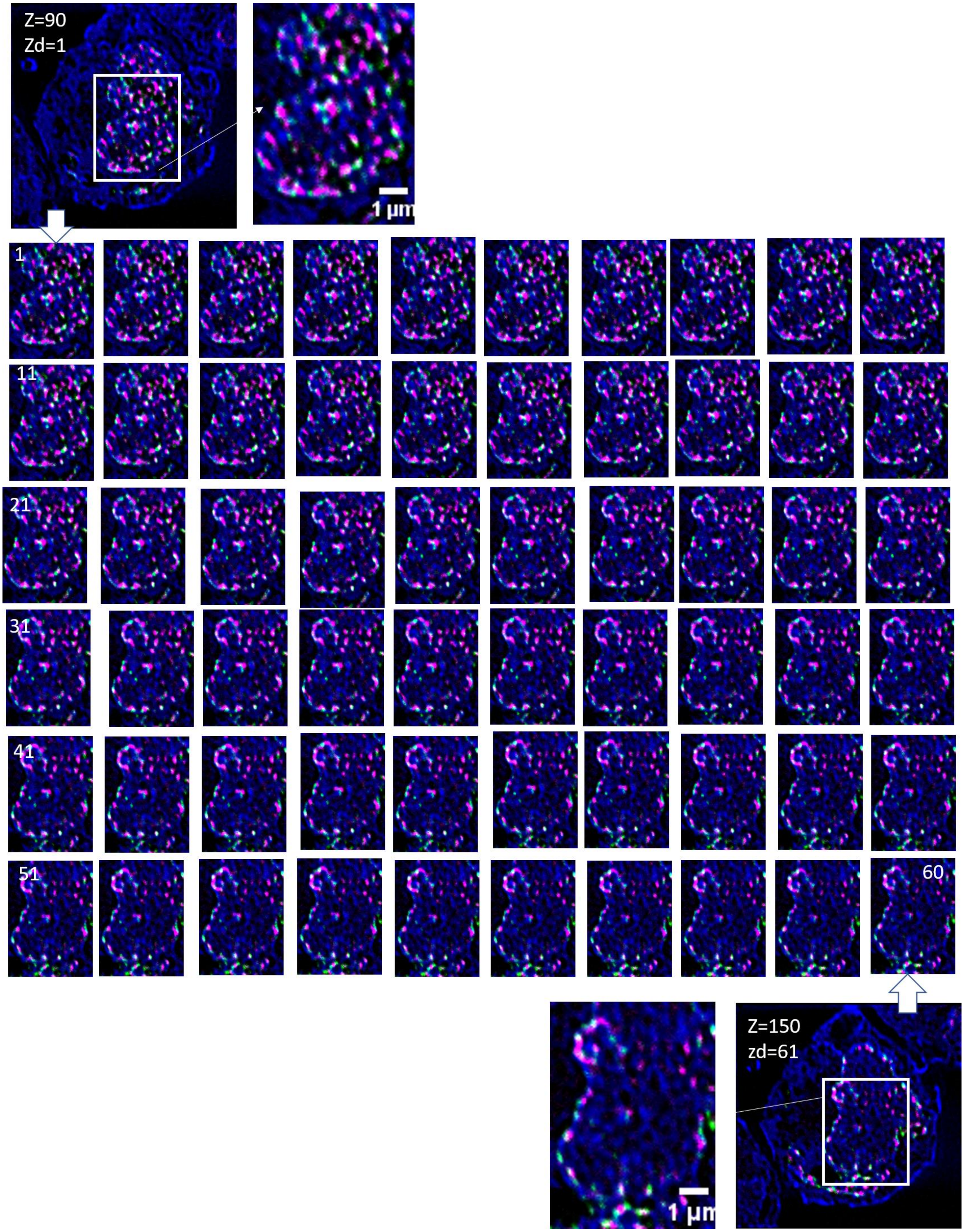

**Supplementary Figure 7.** Estimation of axial resolution by imaging of a single cell.

Similar to the previous figure: $^{31}P$ channel (blue), $^{127}I$-dU (green), and $^{81}Br$-dU (magenta).

Sixty consecutive depth images of a Nalm6 cell shows slight differences at every scan with 5-nm slice thickness indicative of approximately 5-nm axial resolution for IBT.

**Tomographic thickness of a cell for axial resolution analysis**

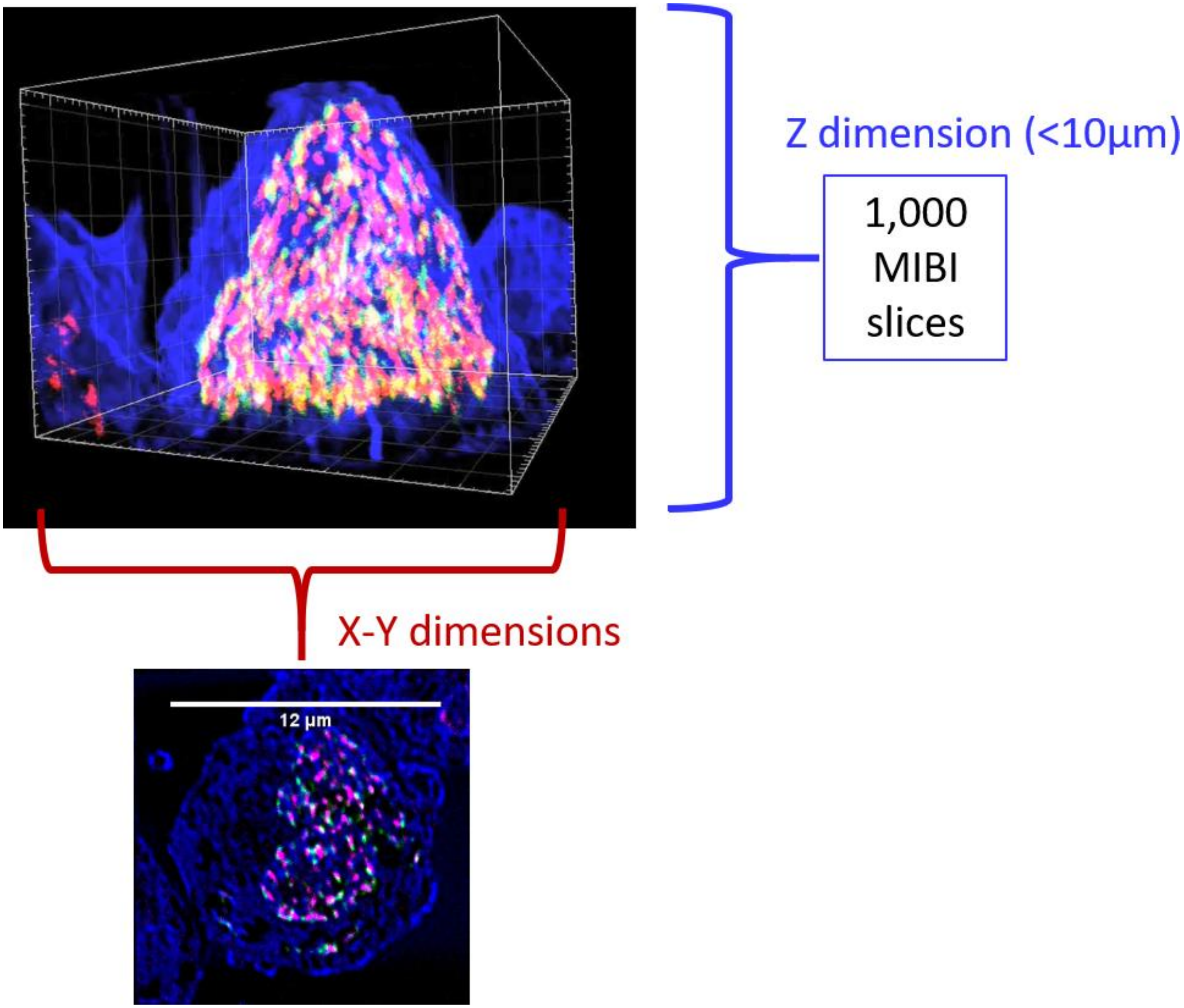

**Supplementary Figure 8.** Estimation of axial resolution by imaging of a single cell.

One thousand scans were etched through a Nalm6 cell. The lateral extent of the cell is between 10 and 12 µm.

Due to the dehydration process and spin coating of cells during preparation, the axial extent is between 5 and 10 µm, yielding an estimate of 5-10 nm axial resolution.

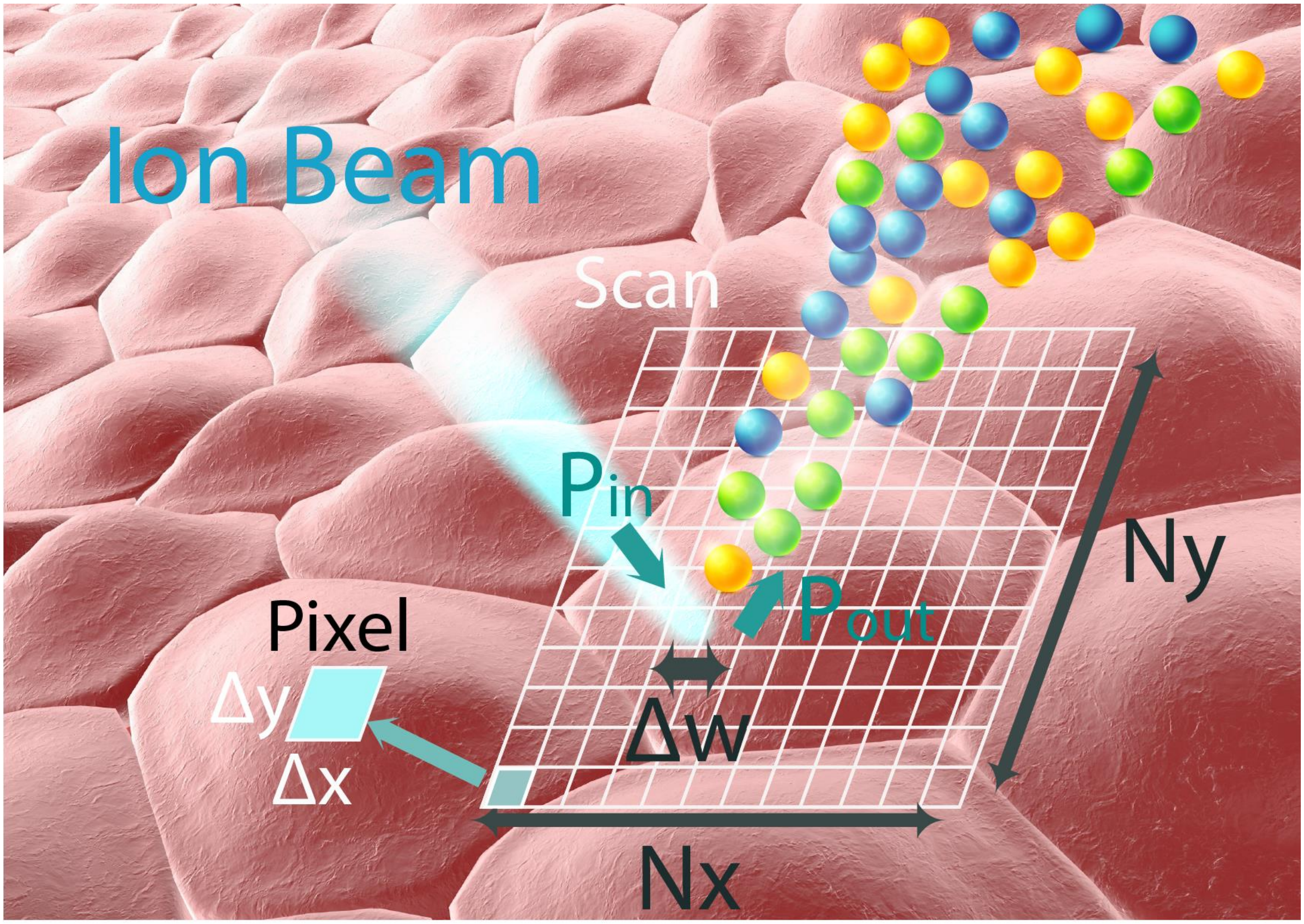
Ion Beam
Scan
$P_{in}$
$P_{out}$
Pixel
Δy
Δx
Δw
Ny
Nx

**Supplementary Figure 9.** Modeling of ion beam imaging parameters.

Ion beam source (blue) corresponds to either Cesium or Oxygen source, impinging on the sample of interest.

After focusing down to sub-100 nm, incoming ion signal ($P_{in}$) to the sample within a small subcellular voxel.

Total secondary ions extracted from the sample ($P_{out}$) and was calculated to be 3%. Distinct color (round objects) correspond to different mass labels.

Pixel size dimensions Δx and Δy of these ion beam images: In the range of 50-150 nm in both dimensions.

Total number of pixels per axis, $N_x$ x $N_y$ pixels, typically 256x 256 pixels, 512x 512 pixels.

Ion beam width of $\Delta w$ that is determined by the ion beam optics and aperture settings: 50-500 nm wide.

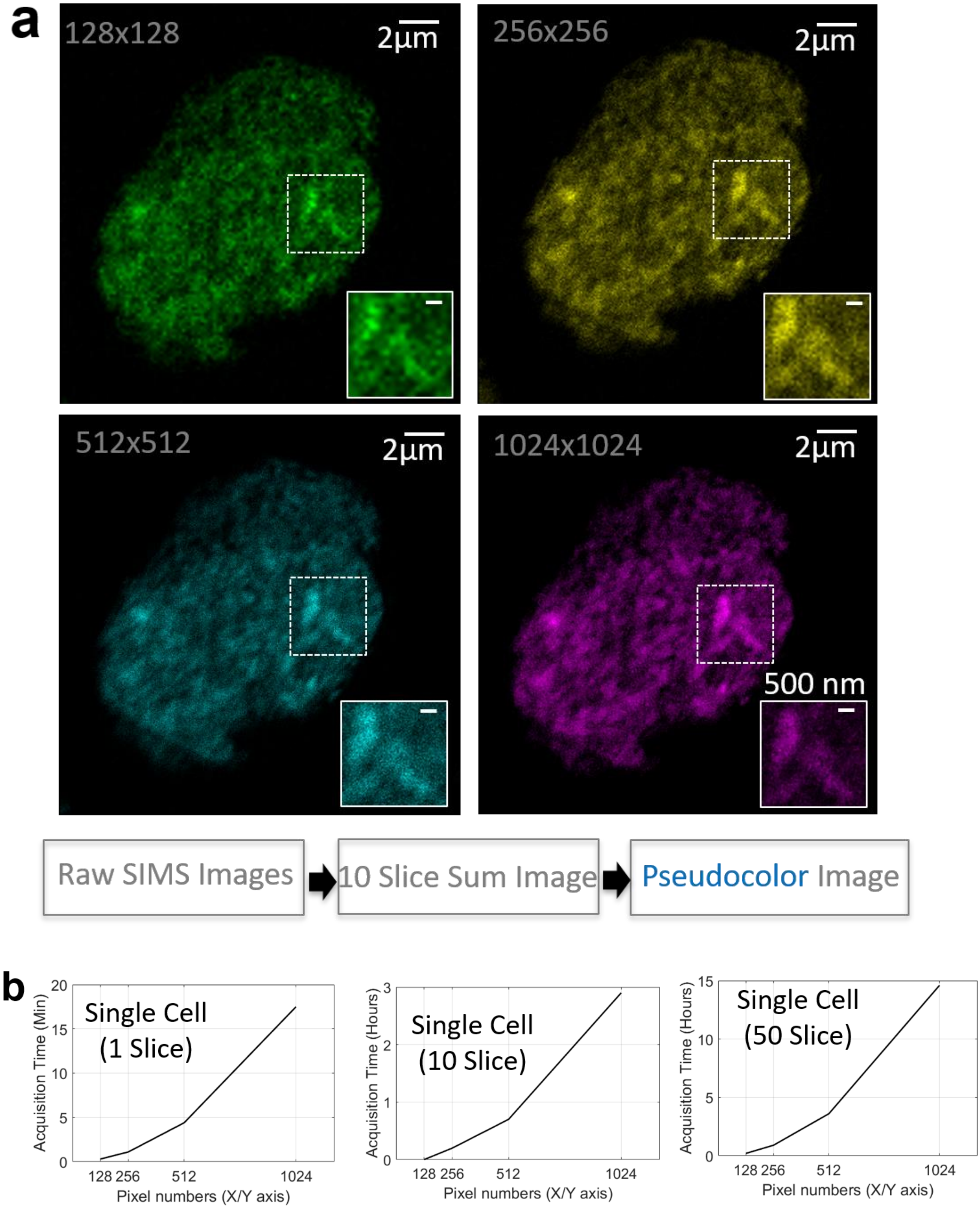

a
128x128
2µm
256x256
2µm
512x512
2µm
1024x1024
2µm
500 nm
Raw SIMS Images
10 Slice Sum Image
Pseudocolor Image
b
Single Cell (1 Slice)
Acquisition Time (Min)
Single Cell (10 Slice)
Acquisition Time (Hours)
Single Cell (50 Slice)
Acquisition Time (Hours)
Pixel numbers (X/Y axis)
128 256 512 1024

**Supplementary Figure 10.** Pixel size effect for ion beam imaging.

(a) HeLa cell DNA imaged by $^{127}I$ channel after 24-hour incubation with $^{127}I$-dU to label DNA.

Four different scans with unique pixel size values were captured for the same cell.

A 25-µm window was imaged and divided into 128 x 128 pixels, 256 x 256 pixels, 512 x 512 pixels, or 124 x 1024 pixels.

Dividing the imaging window by the number of pixels yielded 156 nm, 78 nm, 39 nm, and 19 nm pixel sizes, respectively.

Ten consecutive frames were then summed and pseudocolored in MATLAB. These images were captured with 100-nm PSF width. Imaging pixel size must be smaller than 100 nm to record details of interest.

Thus, the 156-nm pixel size image was not able to resolve details whereas 78-, 39-, and 19-nm pixel sizes provided similar quality images. Sub-50-nm pixel size exposes the same area to ion beams multiple times, creating an oversampling effect in the images.

(b) Image acquisition time scales with numbers of pixels and slices:

Single slice takes more than 15 minutes at 1024x1024 sampling.
Ten slices require up to 3 hours for the same pixel size.
Fifty slices consume about 15 hours with the same pixel dimensions.

A window size of 256 x 256 pixels is the optimal for acquisition time, sampling accuracy, and spatial resolving capability.

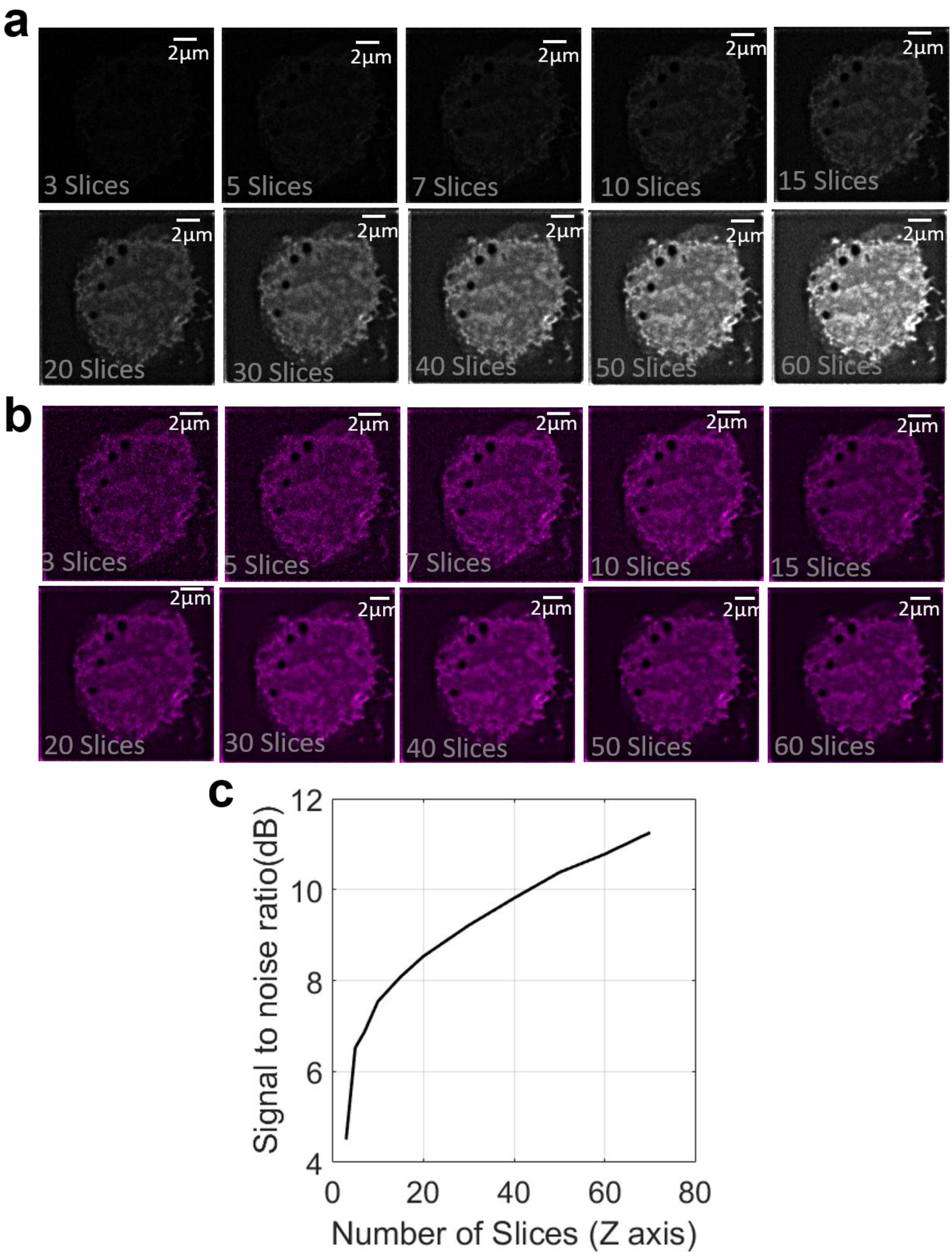
a
2μm
3 Slices
5 Slices
7 Slices
10 Slices
15 Slices
20 Slices
30 Slices
40 Slices
50 Slices
60 Slices
b
3 Slices
5 Slices
7 Slices
10 Slices
15 Slices
20 Slices
30 Slices
40 Slices
50 Slices
60 Slices
c
Signal to noise ratio(dB)
Number of Slices (Z axis)
4
6
8
10
12
0
20
40
60
80

**Supplementary Figure 11.** Multiple depth binning enhances signal-to-noise ratio.

(a) Seventy sections from the same K562 cell were analyzed. The signal-to-noise ratio (SNR) was enhanced as slices were added.

(b) Separately adjusted contrast levels of ion beam images demonstrated optimal spatial details with 10 to 20 slices summed.

(c) The plot of SNR versus number of slices summed indicates that there was exponential enhancement up to 15 slice sums, after which the SNR enhancement provided less improvement.

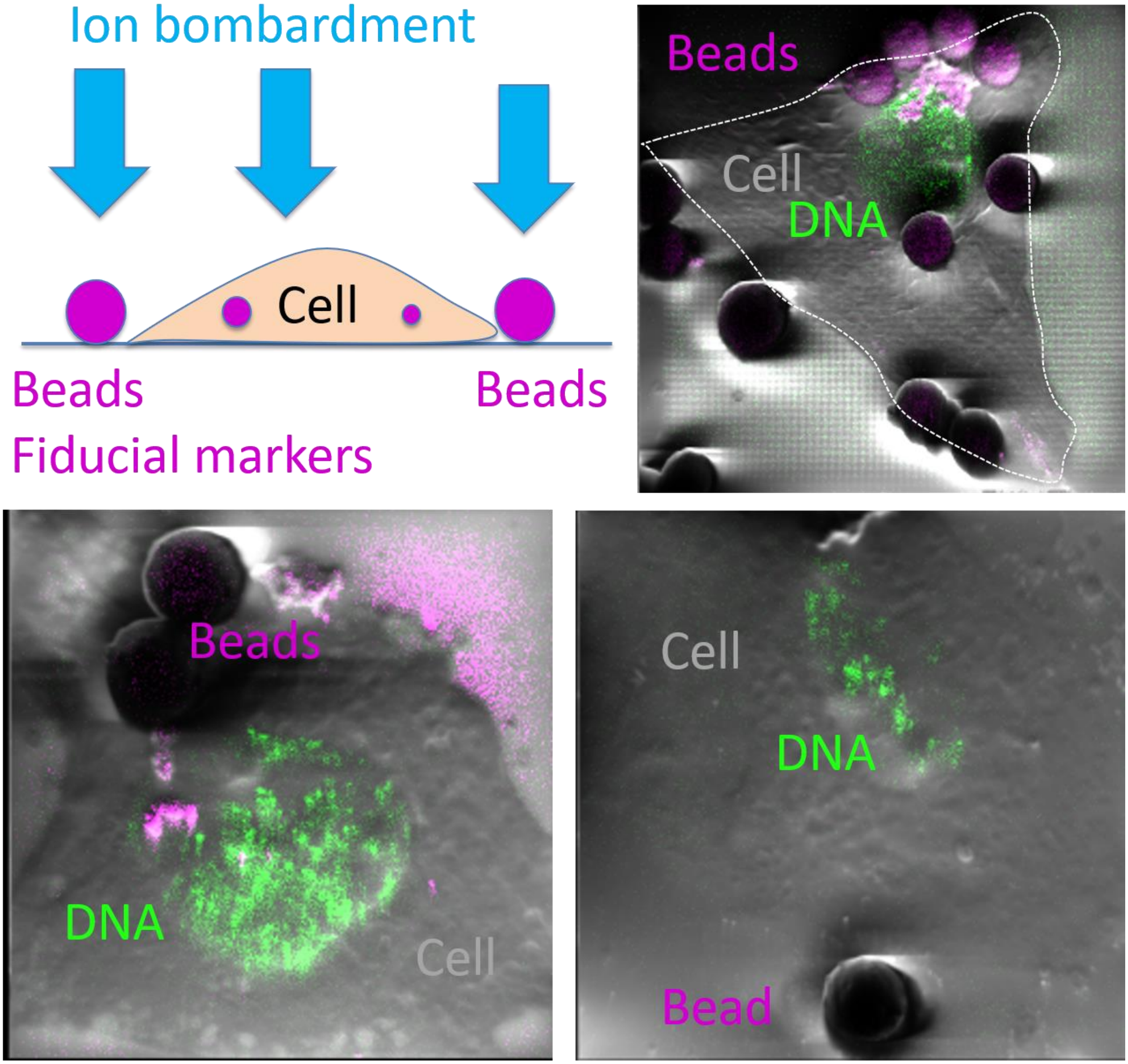
Ion bombardment
Cell
Beads
Beads
Fiducial markers
Beads
Cell
DNA
Beads
DNA
Cell
Cell
DNA
Bead

**Supplementary Figure 12.** Microbead addition enables correction for spatial offsets for each depth image. Beads can be used either outside the cell or inside the cell.

As the bead is made up of harder material than cellular contents, the etching of the bead requires a longer time than the etching of the cellular material. As a result of the differences in etch rates, the bead pattern looks similar across different scans.

The edge or the center of the bead is then used to correct the lateral shifts.

To reduce coagulation, beads are washed with water and ethanol prior to the experiments. The bead solution is then loaded onto the silicon substrate and spin coated at 125 RCF for 5 minutes.

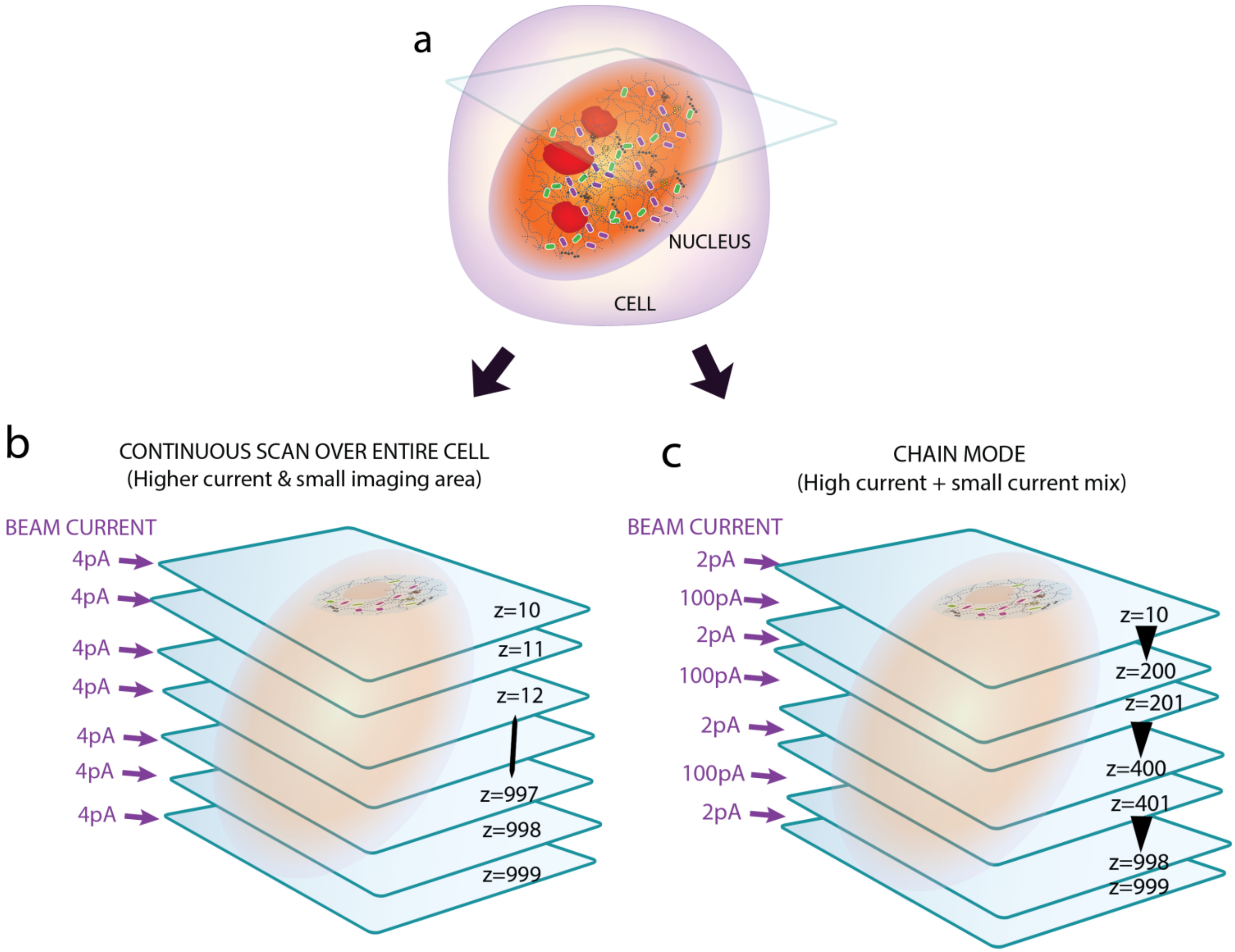

a
NUCLEUS
CELL
b
CONTINUOUS SCAN OVER ENTIRE CELL
(Higher current & small imaging area)
BEAM CURRENT
4pA
4pA
4pA
4pA
4pA
4pA
4pA
z=10
z=11
z=12
z=997
z=998
z=999
c
CHAIN MODE
(High current + small current mix)
BEAM CURRENT
2pA
100pA
2pA
100pA
2pA
100pA
2pA
z=10
z=200
z=201
z=400
z=401
z=998
z=999

**Supplementary Figure 13.** Alternative imaging modes are available.

(a) A single cell is imaged from top to bottom by ion beam tomography.

(b) 3D image acquisition by continuous scans using low current (in the range of 0.2 to 4 pA) only for high resolution mapping of the entire cellular volume.

(c) Another ion imaging modality is chained mode.

Depth scans were acquired by a mix of high current (100pA) and low current (e.g., 2 pA), enhancing overall IBT throughput.

HeLa Cells (IdU Labeled DNA at $^{127}$I Channel)

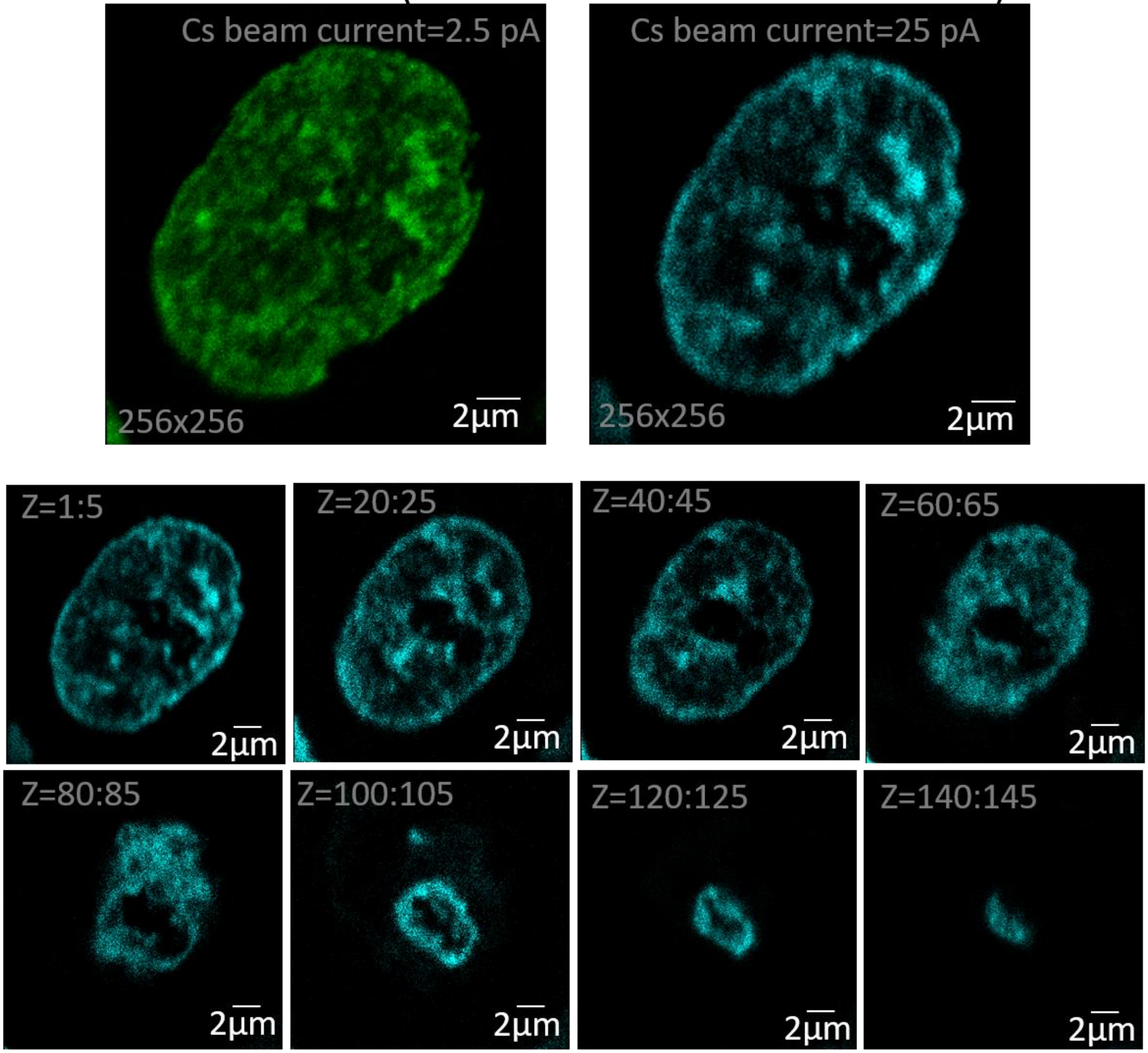

**Supplementary Figure 14.** Rapid depth imaging at decent spatial resolution.

HeLa cell was labeled with $^{127}$I-dU for tagging DNA to be imaged by ion beams.

A higher current level (25 pA) was used to acquire the cell across 150 slices, rather than capturing up to 1,000 at 2.5 pA.

This approach reduced imaging time per cell, allowing to scan more cell in shorter duration.

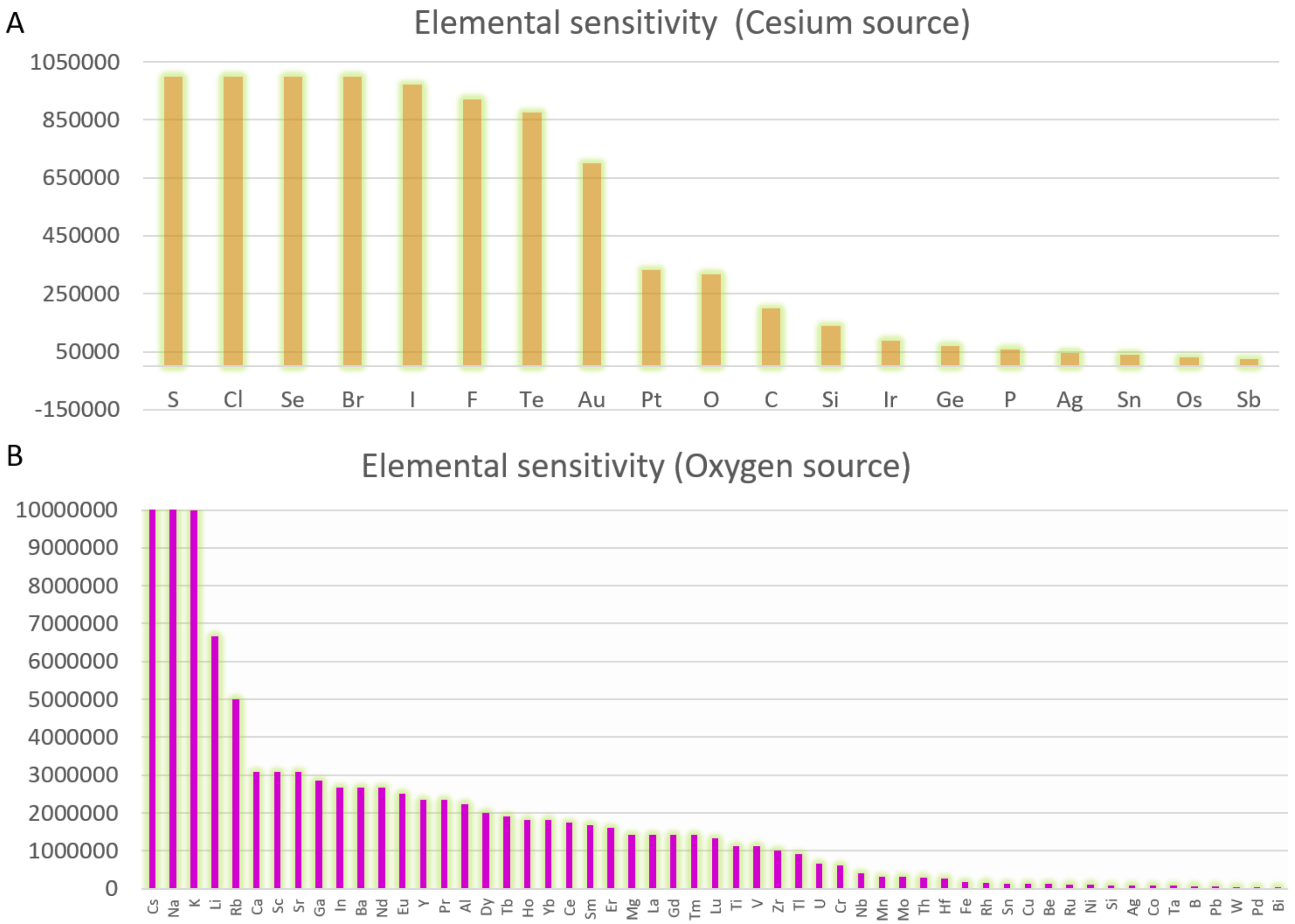
A
Elemental sensitivity (Cesium source)
1050000
850000
650000
450000
250000
50000
-150000
S Cl Se Br I F Te Au Pt O C Si Ir Ge P Ag Sn Os Sb
B
Elemental sensitivity (Oxygen source)
10000000
9000000
8000000
7000000
6000000
5000000
4000000
3000000
2000000
1000000
0
Cs Na K Li Rb Ca Sc Sr Ga In Ba Nd Eu Y Pr Al Dy Tb Ho Yb Ce Sm Er Mg La Gd Tm Lu Ti V Zr Tl U Cr Nb Mn Mo Th Hf Fe Rh Sn Cu Be Ru Ni Si Ag Co Ta B Pb W Pd Bi

**Supplementary Figure 15.** Elemental sensitivity analysis of ion images acquired using cesium and oxygen sources.

The raw values were extracted from a previous publication (*Wilson. SIMS quantification in Si, GaAs, and diamond – an update. International Journal of Mass spectroscopy and Ion Processes, 143, 43-49, 1995; Table 1*).

For the cesium source-based ion beam imaging, the relative sensitivity factors (RSF) of $E^-$ RSF[b] (Negative secondary ions measured by 14.5 keV Cs+ source) were divided by the max value of the RSF (lowest sensitivity number), providing a direct measure of sensitivity.

For the oxygen source, $E^+$ RSF[a] (Positive secondary ions measured by 4 keV O+ source) values were also normalized by the largest number to enable sensitivity comparisons.

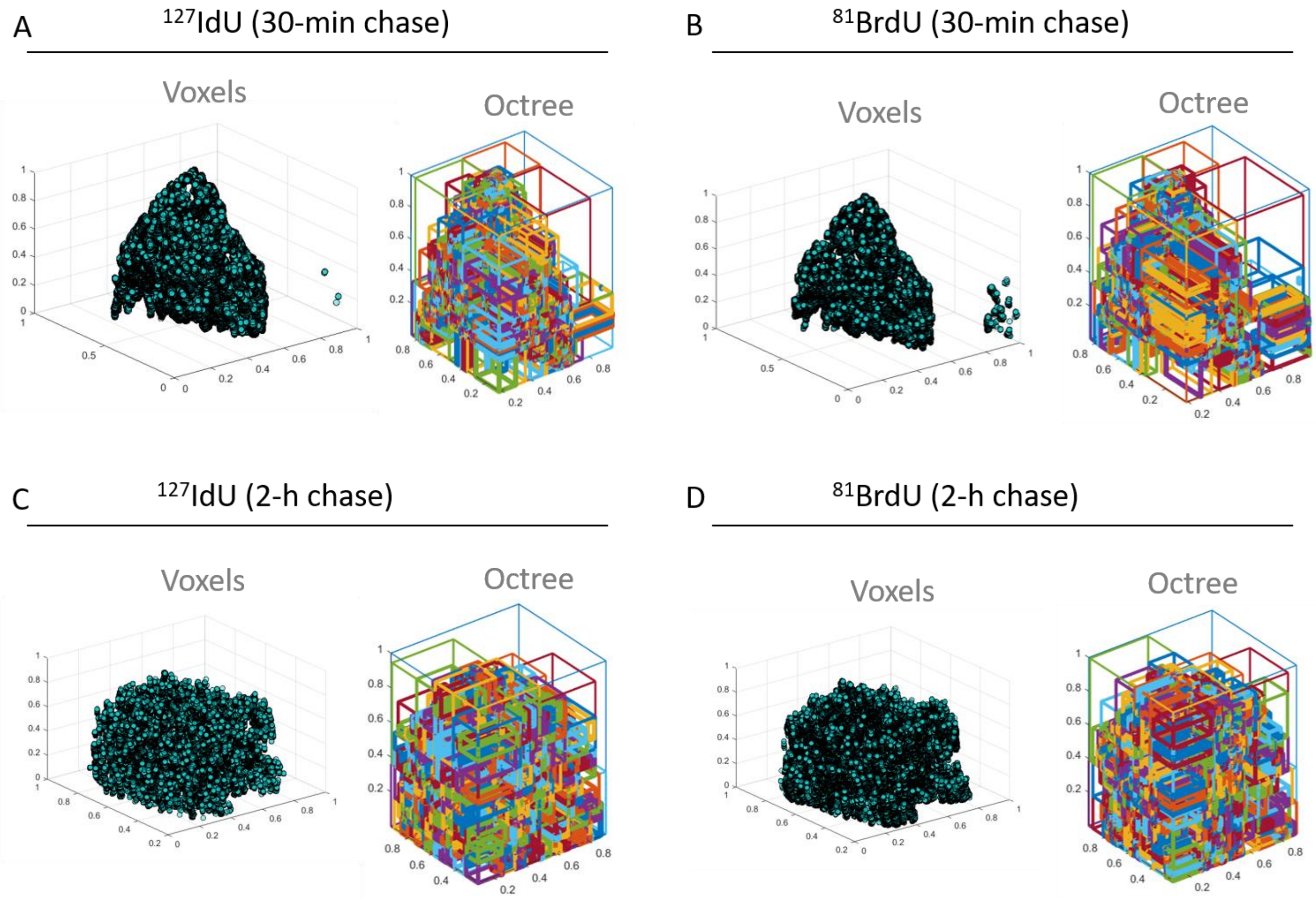
A
127IdU (30-min chase)
Voxels
Octree
B
81BrdU (30-min chase)
Voxels
Octree
C
127IdU (2-h chase)
Voxels
Octree
D
81BrdU (2-h chase)
Voxels
Octree

**Supplementary Figure 16.** Voxel distributions and octree representations of replication forks in Nalm6 cells. Ion beam signal distribution in a single cell volume was partitioned into the eight octant subvolumes as part of the octree algorithm. The boxes show the density of ion signal within that particular volumetric space.

(a) Left corresponds to the cell that was pulsed with $^{127}$I-dU incorporation for 30 minutes, and

(b) the right image shows another 30 minutes pulse with $^{81}$Br-dU, that were chased 30 minutes in between the pulses in the culture. Octree distributions in these upper panels showed similar box patterns.

Similarly,

(c) lower left demonstrates I-dU and (d) right shows Br-dU incorporation separated with 2 hours chase time. Octree method yielded repeatable box patterns in the single cell volume.

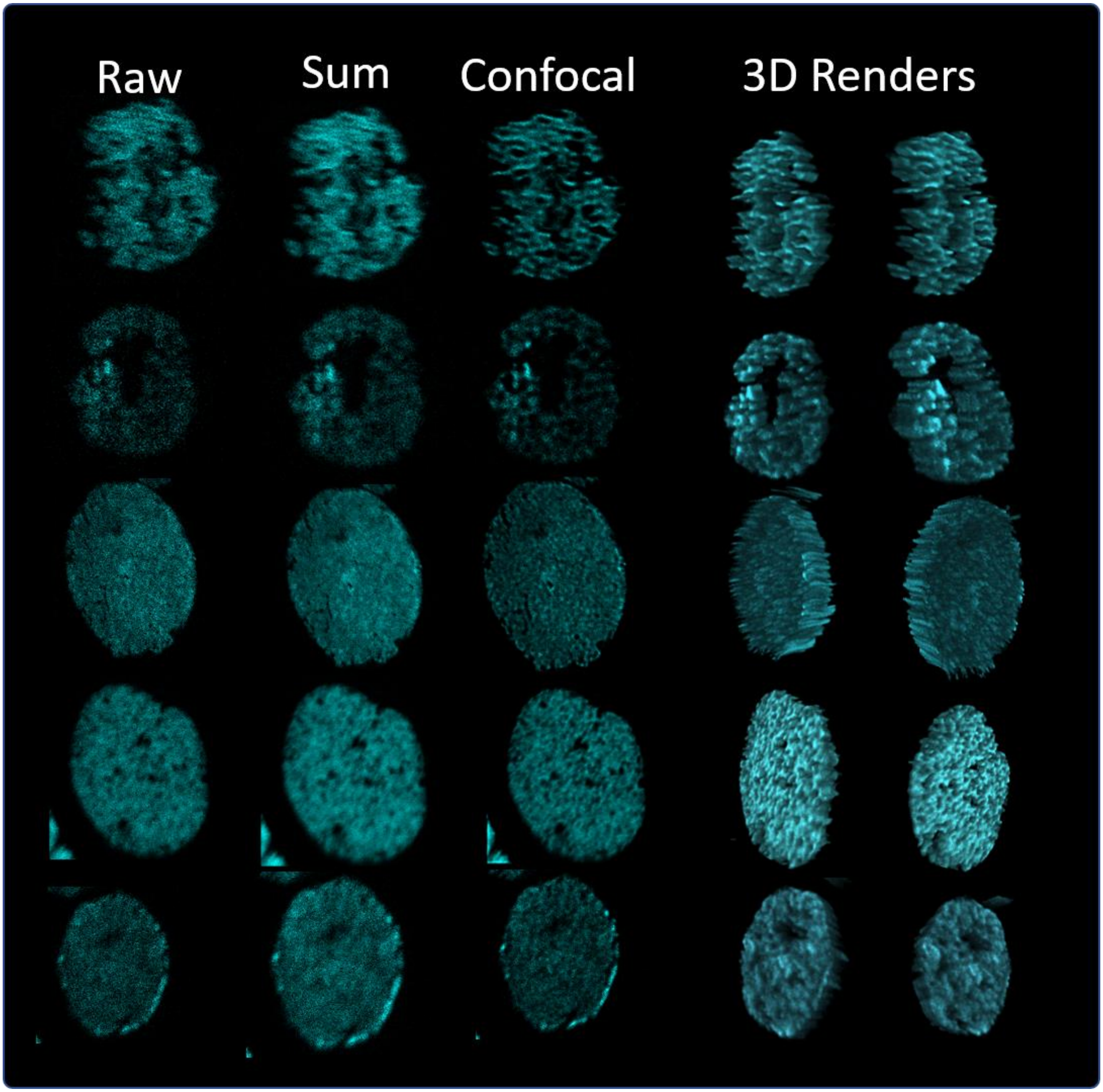
Raw
Sum
Confocal
3D Renders

**Supplementary Figure 17.** IBT approach was validated in genomic images by $^{127}$I channel using 24 hours of $^{127}$I-dU labeling in Jurkat cells (top 2 rows) and HeLa cells (bottom 3 rows).

Raw: Acquired ion beam at each depth scan.

Sum: Binning of 5 subsequent depth slices.

Confocal: Digitally deconvolved images on binned sections.

3D Renders from the reconstructed ion beam tomographic slices from two different viewing angles.

The high quality of final images compared to the initial ion images demonstrated the benefit of the presented mathematical analysis pipeline.

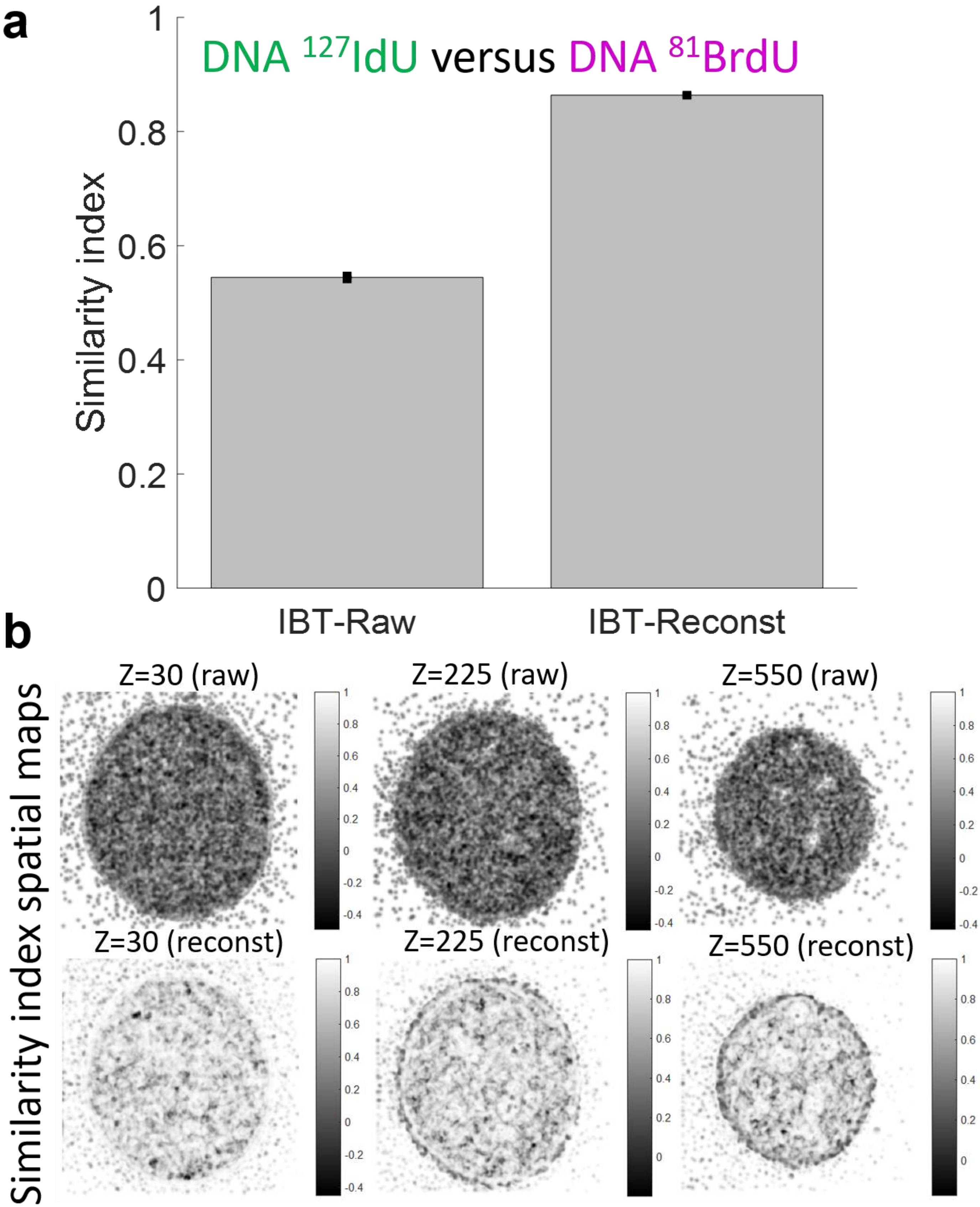
a
DNA 127IdU versus DNA 81BrdU
Similarity index
1
0.8
0.6
0.4
0.2
0
IBT-Raw
IBT-Reconst
b
Similarity index spatial maps
Z=30 (raw)
Z=225 (raw)
Z=550 (raw)
Z=30 (reconst)
Z=225 (reconst)
Z=550 (reconst)

**Supplementary Figure 18.** Structural similarity index (SSIM) values were used to quantify the "similarity" measure of DNA images at the $^{127}$I-dU and $^{81}$Br-dU channels for the raw and reconstructed ion images that were presented in Fig. 3.

(a) Reconstructed DNA images exhibit significantly higher similarity (SSIM: 0.86 ±SE 0.001) compared to the raw DNA images (0.54 ±SE 0.0037).

IBT-raw: SSIM values for measuring the similarity of the raw DNA images from $^{127}$I-dU and $^{81}$Br-dU.

IBT-reconst: SSIM values for calculating the similarity of the reconstructed images from $^{127}$I-dU and $^{81}$Br-dU.

*n* = 737 sections were used in the bar graph with mean and standard error of SSIM values.

(b) Spatial map of SSIM quantification in DNA images for 30$^{th}$, 225$^{th}$, and 550$^{th}$ ion beam tomographic slices.

(Top row) SSIM index values per pixels were plotted as a spatial map for raw DNA images. Most pixels average around dark grey SSIM values, suggesting high dissimilarity.

(Bottom row) SSIM spatial values for reconstructed DNA images. Most pixels span around light grey SSIM values, providing high degree similarity between images.

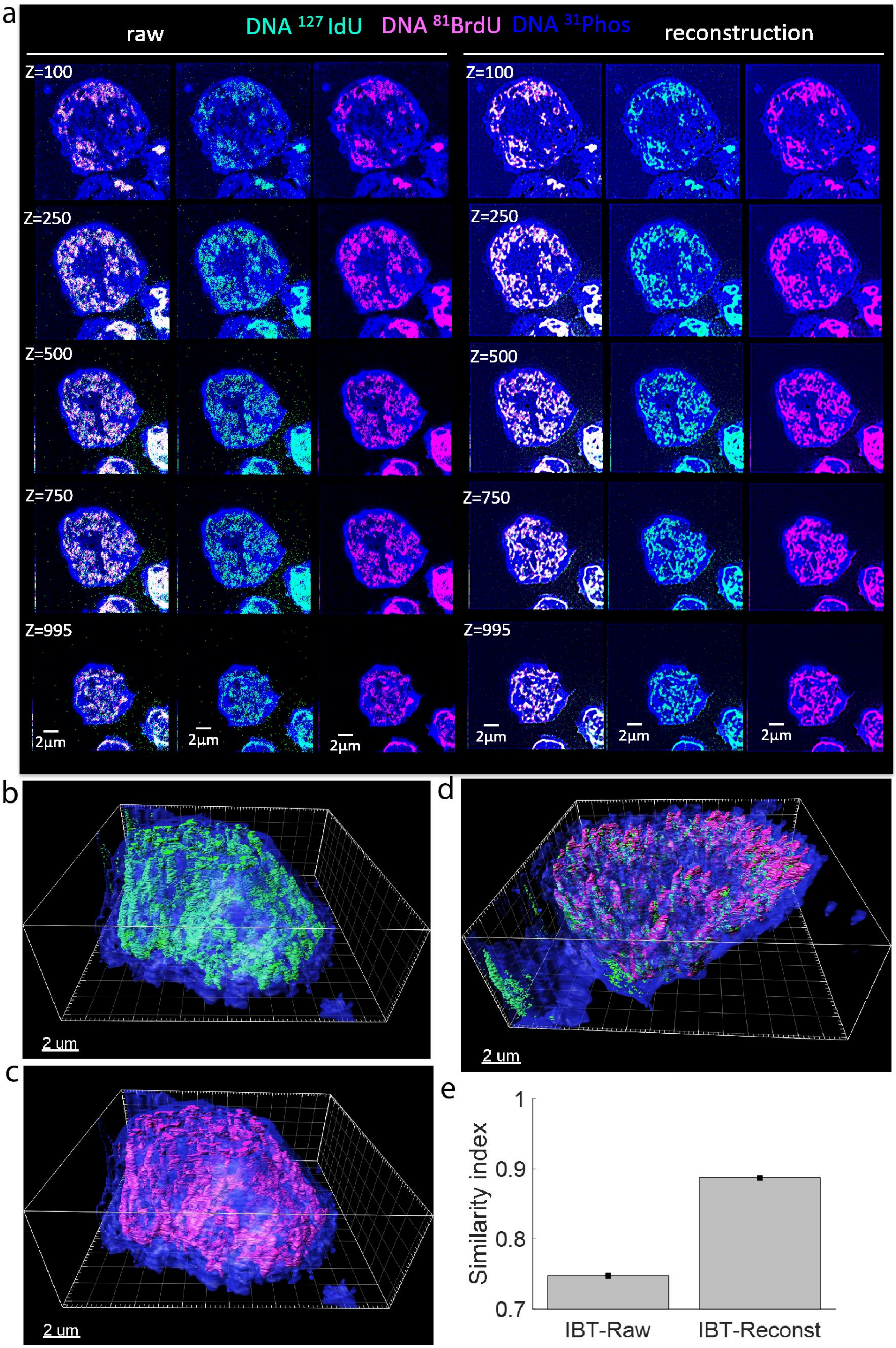
a
raw
DNA 127 IdU
DNA 81BrdU
DNA 31Phos
reconstruction
Z=100
Z=250
Z=500
Z=750
Z=995
2µm
b
2 um
c
2 um
d
2 um
e
Similarity index
1
0.9
0.8
0.7
IBT-Raw
IBT-Reconst

**Supplementary Figure 19.** SSIM values of DNA images at the $^{127}$I-dU and $^{81}$Br-dU channels for the raw and reconstructed ion images in a Nalm6 cell.

(a) Similar to the Figure 3, but for Nalm6 cells. Raw images of replicated DNA (green and magenta) and largely chromatin (blue) are noisy and reconstructed images show spatially resolved patterns in the chromatin.
(b) 3D renders of Nalm6 ion beam tomogram. Replicated DNA by $^{127}$I-dU (green) and largely chromatin by $^{31}$P (blue).
(c) 3D visualization of Nalm6 with Replicated DNA by $^{81}$Br-dU (magenta) and largely chromatin by $^{31}$P (blue).

(d) Inverted 3D position of the same Nalm6 cell with $^{127}$I-dU (green), $^{81}$Br-dU (magenta), and chromatin by $^{31}$P (blue).

(e) Reconstructed chromatin images exhibit higher similarity (SSIM: 0.88 ±SE 0.001) compared to the raw DNA images (0.74 ±SE 0.001).

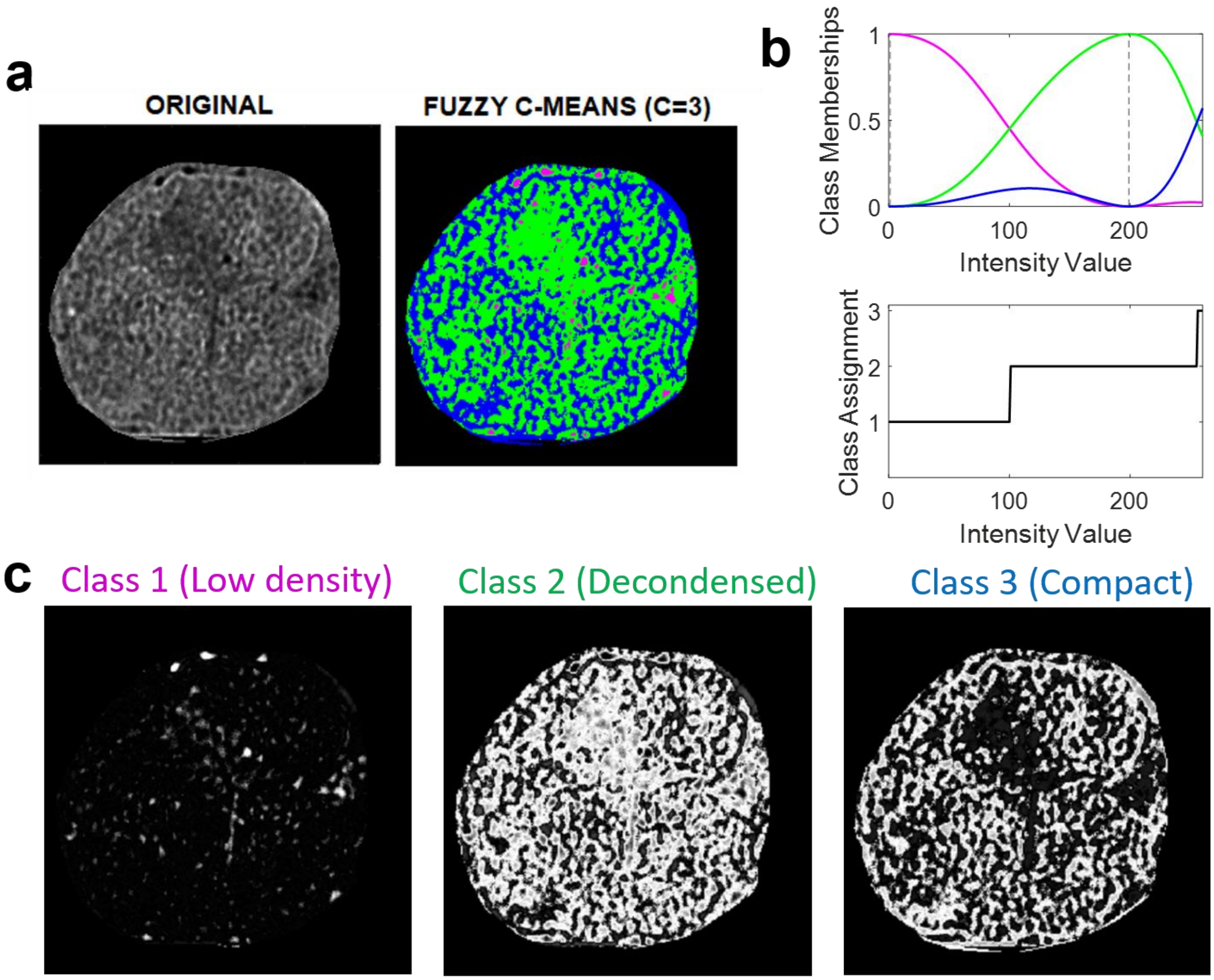
a
ORIGINAL
FUZZY C-MEANS (C=3)
b
Class Memberships
Intensity Value
Class Assignment
Intensity Value
c
Class 1 (Low density)
Class 2 (Decondensed)
Class 3 (Compact)

**Supplementary Figure 20.** Fuzzy logic segmentation in a B cell lymphoblast for 50th slice.

(a) Original image (improved by mathematical pipeline) was used for segmentation process. A fuzzy C-means clustering (C=3) was calculated for the $^{31}P$ image based on intensity values.

(b) Pixel values were sorted into class memberships based on density of chromatin distribution.

(c) Each ion image corresponding to the assignments of class 1 (low chromatin density), class 2 (decondensed chromatin), and class 3 (compact chromatin).

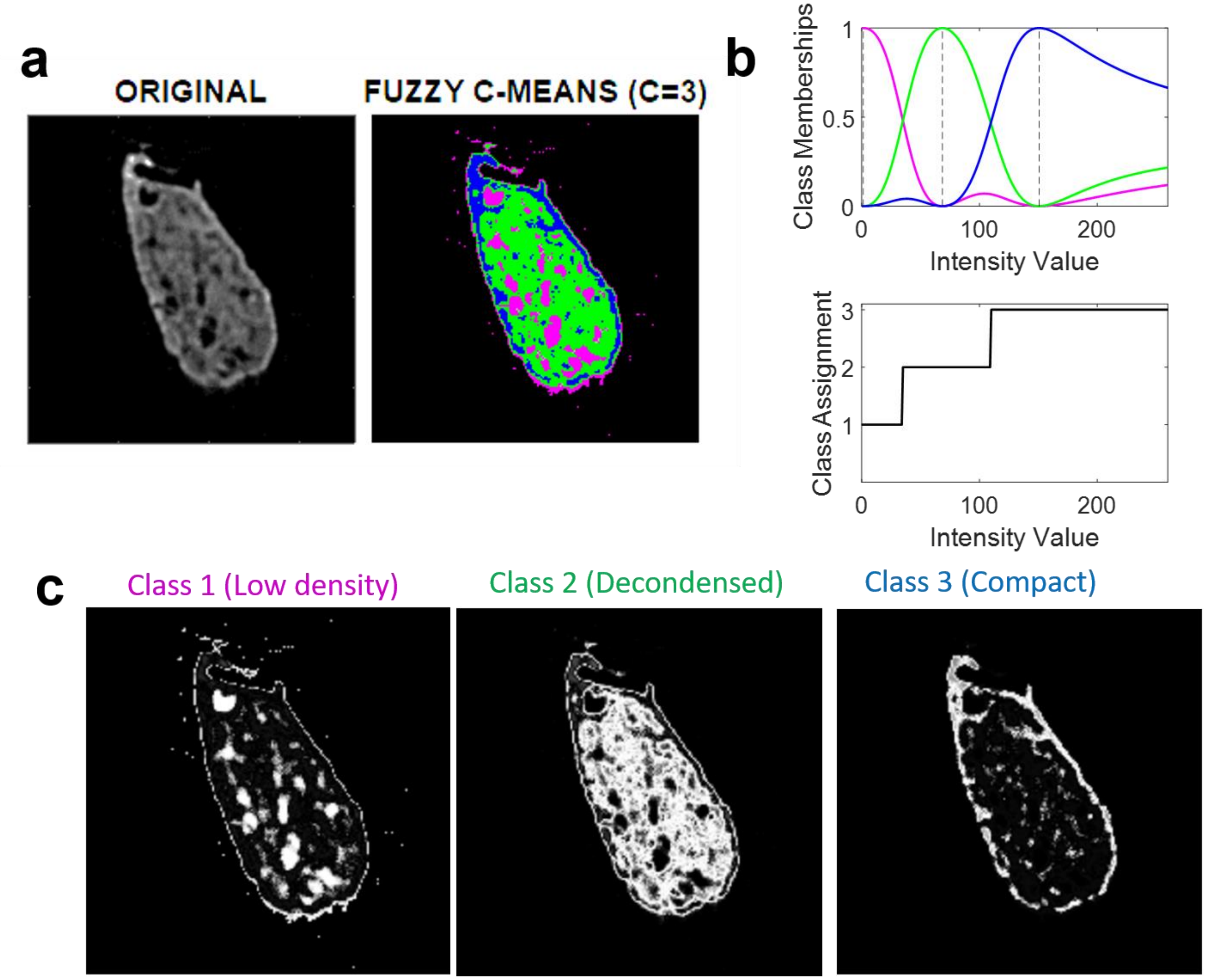
a
ORIGINAL
FUZZY C-MEANS (C=3)
b
Class Memberships
1
0.5
0
0
100
200
Intensity Value
Class Assignment
3
2
1
0
100
200
Intensity Value
c
Class 1 (Low density)
Class 2 (Decondensed)
Class 3 (Compact)

**Supplementary Figure 21.** Fuzzy logic segmentation in a HeLa cell for 500$^{th}$ slice.

Similar to the previous Supplementary Figure 15,

(a) C-means clustering (C=3) was used to sort the pixel values,

(b) class assignments for three density levels of chromatin, and

(c) segmented images of chromatin for classes 1-3.

Instead of using direct calculation of intensity values, Fuzzy C-means based clustering of ion images provided determination of chromatin states without manually changing intensity threshold levels for segmentation.

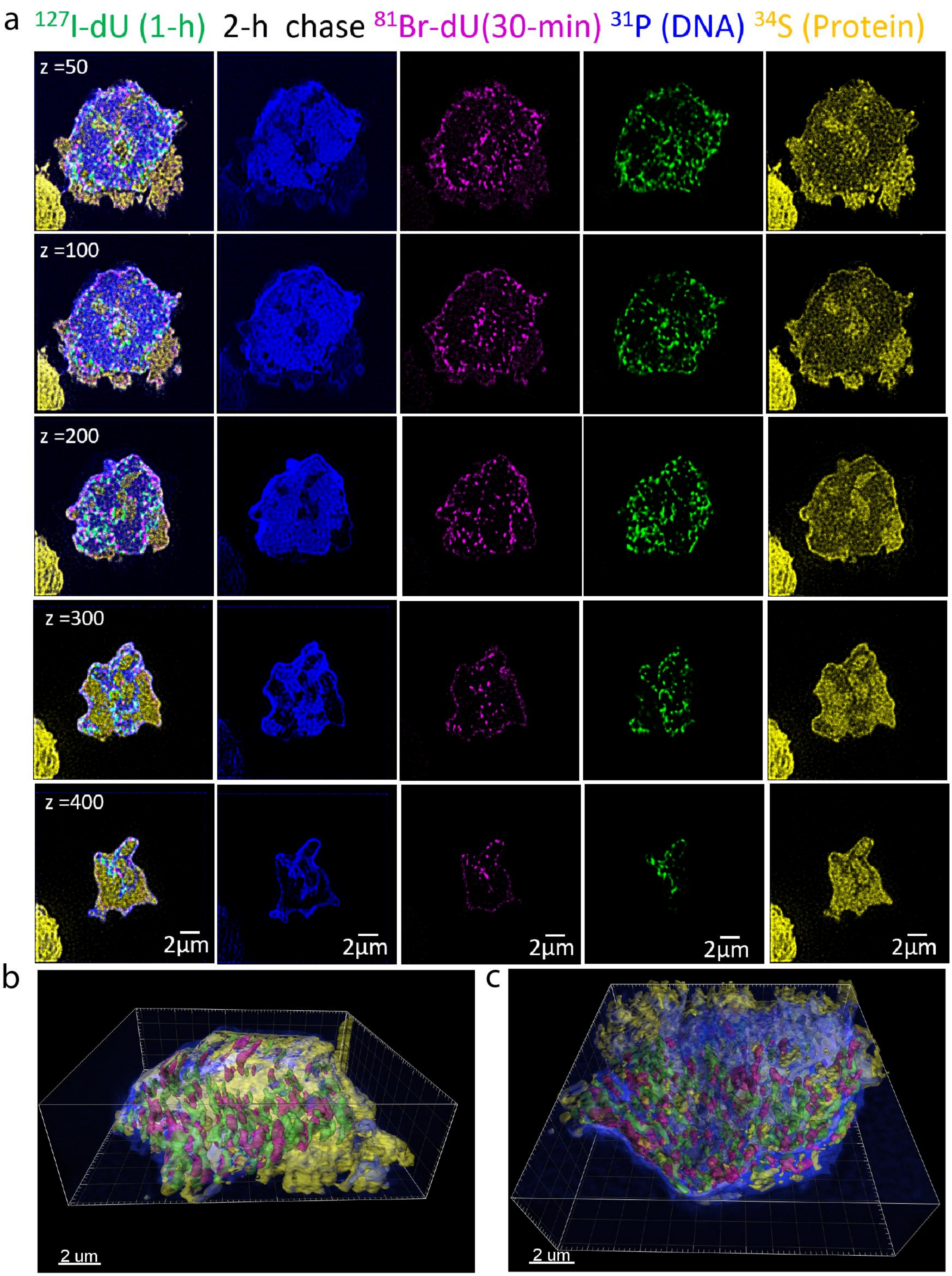
a
$^{127}$I-dU (1-h) 2-h chase $^{81}$Br-dU(30-min) $^{31}$P (DNA) $^{34}$S (Protein)
z =50
z =100
z =200
z =300
z =400
2μm
2μm
2μm
2μm
2μm
b
c
2 um
2 um

**Supplementary Figure 22.** Four channel representation of replication data in Figure 4 (2-h chase).

(a) Nalm6 cell was pulsed with $^{127}$I-dU (green) for 1 hour, and then chased in culture media for 2 hours, followed by a second pulse with $^{81}$Br-dU (magenta) for 30 minutes. IBT image sections for 50$^{th}$, 100$^{th}$, 200$^{th}$, 300$^{th}$, and 400$^{th}$ slices were shown. $^{31}$P (blue) was used for chromatin analysis. $^{34}$S (yellow) was utilized as an indicator for protein concentration.

(b) 3D visualization of 600 slices in a Nalm6 cell. Replication sites (green and magenta), chromatin (blue), and proteins (yellow). Higher concentration of replication forks was around the half bottom of the cell.

(c) An inverted 3D renders of the same Nalm6 cell with four colors as denoted.

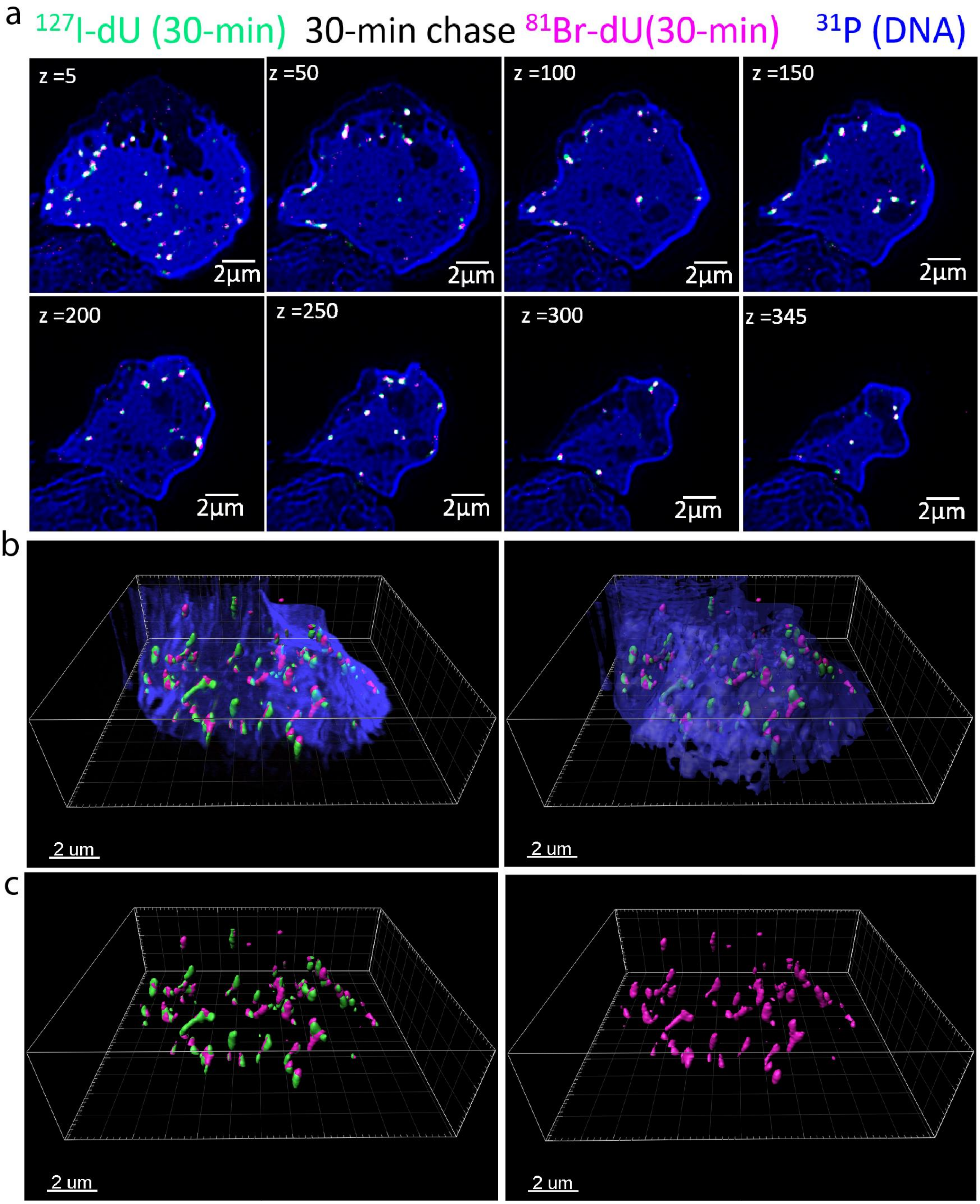
a
127I-dU (30-min) 30-min chase 81Br-dU(30-min) 31P (DNA)
z =5
z =50
z =100
z =150
z =200
z =250
z =300
z =345
2μm
b
2 um
c
2 um

**Supplementary Figure 23.** Biological replicate experiment for replication dynamics in Figure 4 (30-min chase).

(a) Nalm6 cell was pulsed with $^{127}$I-dU (green) for 30 minutes, and then chased in culture media for 30 minutes, followed by a second pulse with $^{81}$Br-dU (magenta) for 30 minutes. Digitally processed IBT image sections for 5$^{th}$, 50$^{th}$, 100$^{th}$, 150$^{th}$, 200$^{th}$, 250$^{th}$, 300$^{th}$, and 345$^{th}$ slices were shown. $^{31}$P (blue) was used to identify largely chromatin.

(b) 3D renders of 350 sections in the Nalm6 cell. Surface function was used to colorize replication site images. The 31P channel was original data (left) and surface renders with 50%transperancy setting in the 3D render.

(c) Only replication forks (green and red) were highlighted in 3D, showing significant overlaps and variability in spatial distribution in cellular volumes.

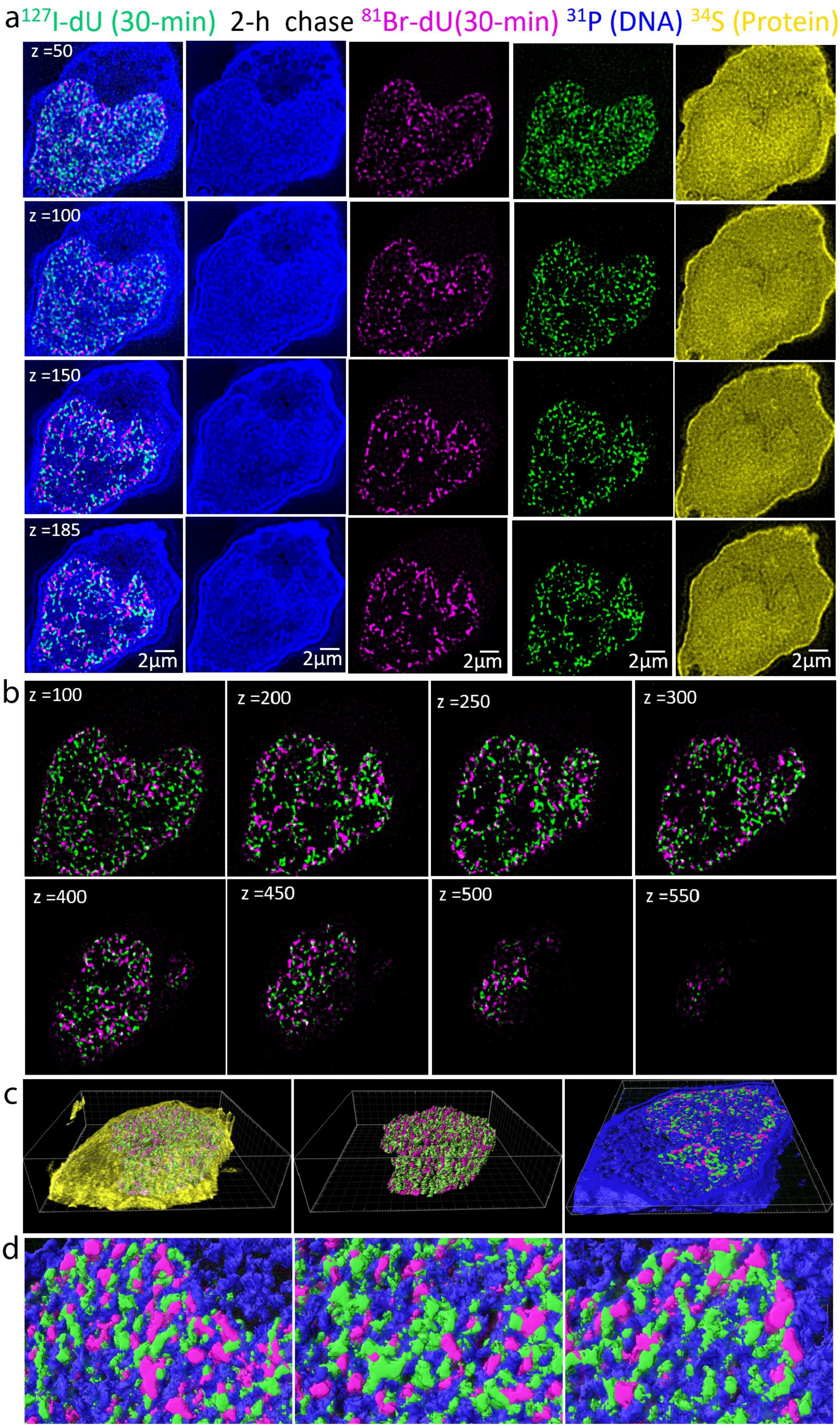
a
127I-dU (30-min) 2-h chase 81Br-dU(30-min) 31P (DNA) 34S (Protein)
z =50
z =100
z =150
z =185
2μm
2μm
2μm
2μm
2μm
b
z =100
z =200
z =250
z =300
z =400
z =450
z =500
z =550
c
d

**Supplementary Figure 24.** Biological repeat for replication dynamics in Figure 4 (2-h chase).

(a) Nalm6 cell was pulsed with $^{127}$I-dU (green) for 30-min, and then chased in culture media for 2 hours, followed by a second pulse with $^{81}$Br-dU (magenta) for 30 minutes. IBT image sections for 50$^{th}$, 100$^{th}$, 180$^{th}$, and 185$^{th}$ slices were shown. $^{31}$P (blue) was used for chromatin and $^{34}$S (yellow) was imaged for proteins.

(b) Overlaps of replication forks are minimal. 2D sections from 100$^{th}$, 200$^{th}$, 250$^{th}$, 300$^{th}$, 400$^{th}$, 500$^{th}$, and 550$^{th}$ ion images.

(c) 3D renders of 600 slices in a Nalm6 cell. Replication sites (green and magenta), proteins (yellow), and chromatin (blue).

(d) Zoomed images from chromatin and replication forks, extracted from a 3D renders with 185 slices.

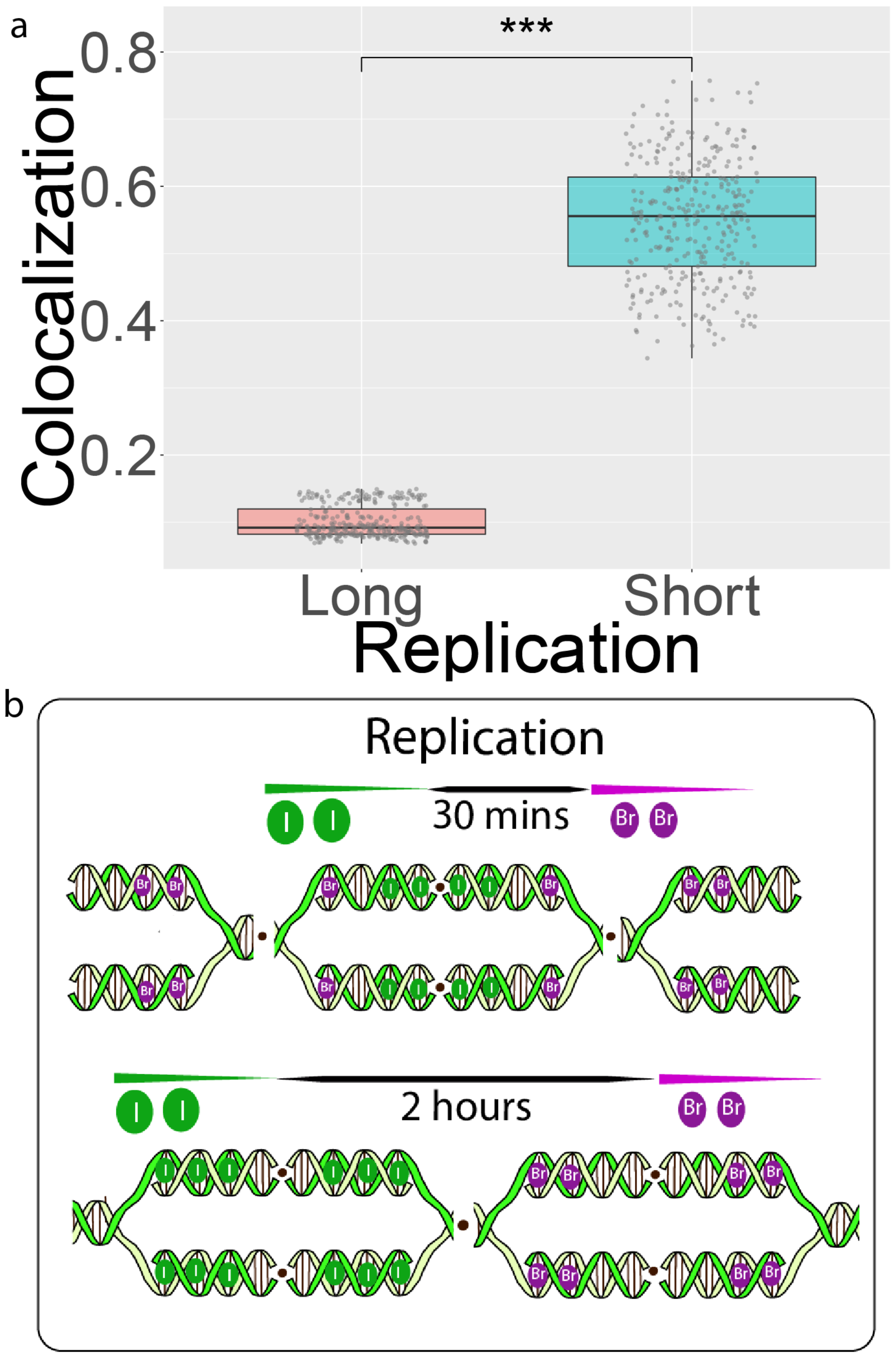
a
***
Colocalization
0.8
0.6
0.4
0.2
Long
Short
Replication
b
Replication
30 mins
I
Br
2 hours

**Supplementary Figure 25.** Spatial overlaps of replication forks were higher in short chase experiments.

(a) Correlation coefficient (CC) of 30-min chase experiments (from Supplementary Fig. 23) and 2-h chase experiment (from Supplementary Fig. 24) was computed. Long chase provided 0.55 CC and short chase exhibited 0.09 CC values, demonstrating the significantly more overlaps in 30-min compared to 2-h chase. Each dot corresponds to a single depth slice from the ion beam tomographic data.

(b) Schematic model of spatial overlaps in replication dynamics. The first pulse (Green, $^{127}$I-dU) and second pulse ($^{81}$Br-dU) occurs in similar replication loci when the time difference between the two is shorter.

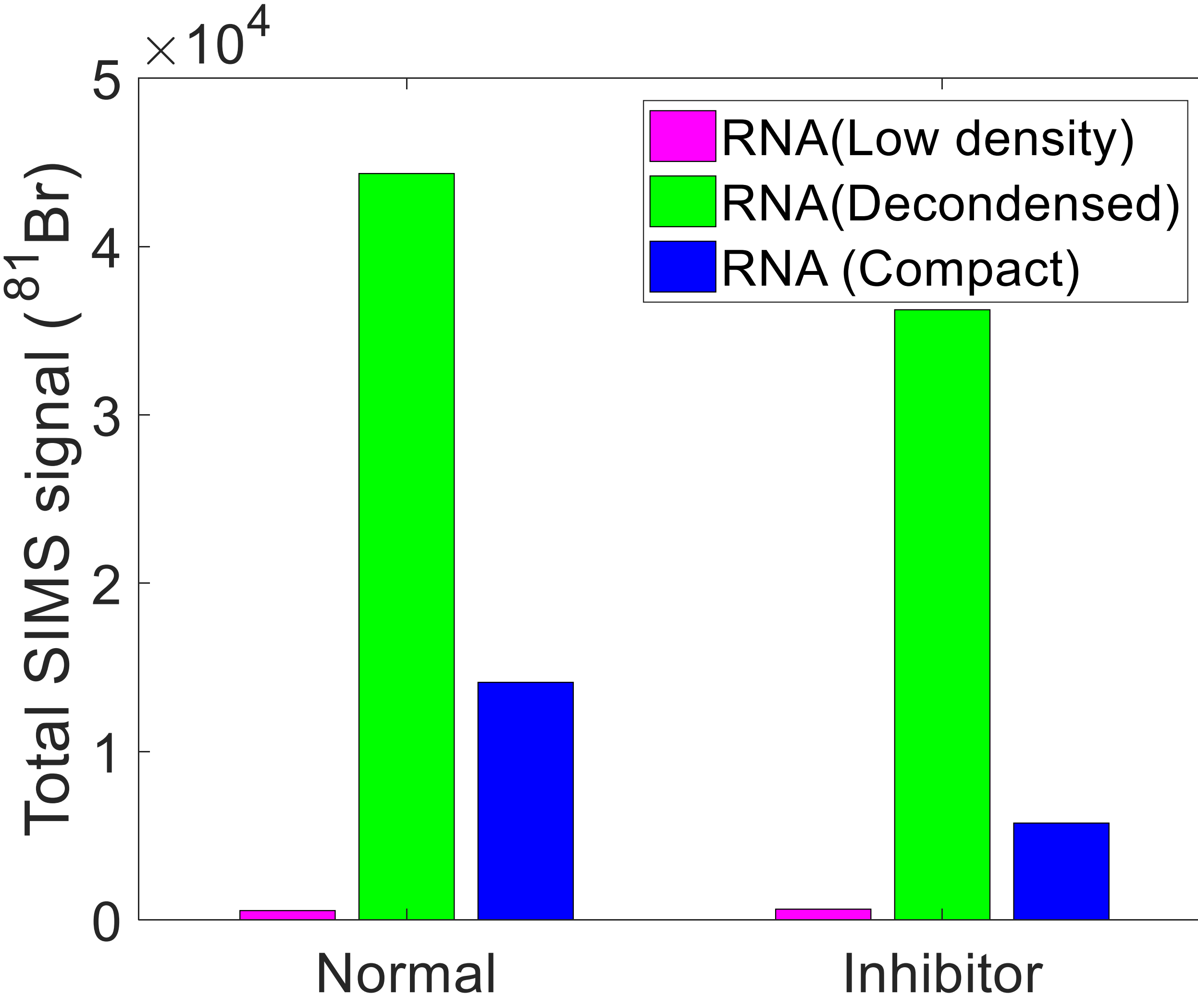
×10^4
5
4
3
2
1
0
Total SIMS signal (^81Br)
RNA(Low density)
RNA(Decondensed)
RNA (Compact)
Normal
Inhibitor

**Supplementary Figure 26.** Newly synthesized RNAs are enriched in the decondensed region of the chromatin. 30 minutes of 81 Br-rU incorporation into Nalm6 cells.

Nascent RNA distribution at the 15$^{th}$ section of the first Nalm6 cell (Normal) showed high concentration in the decondensed region. Lower RNA concentration was in the compact and low-density chromatin regions.

In the existence of α-amanitin drug for 30 minutes after the RNA labeling in the second Nalm6 cell (inhibitor), newly synthesized RNA distribution was reduced in the decondensed as well as low density chromatin regions. Nascent RNA transcripts were partially repressed due to the degradation.

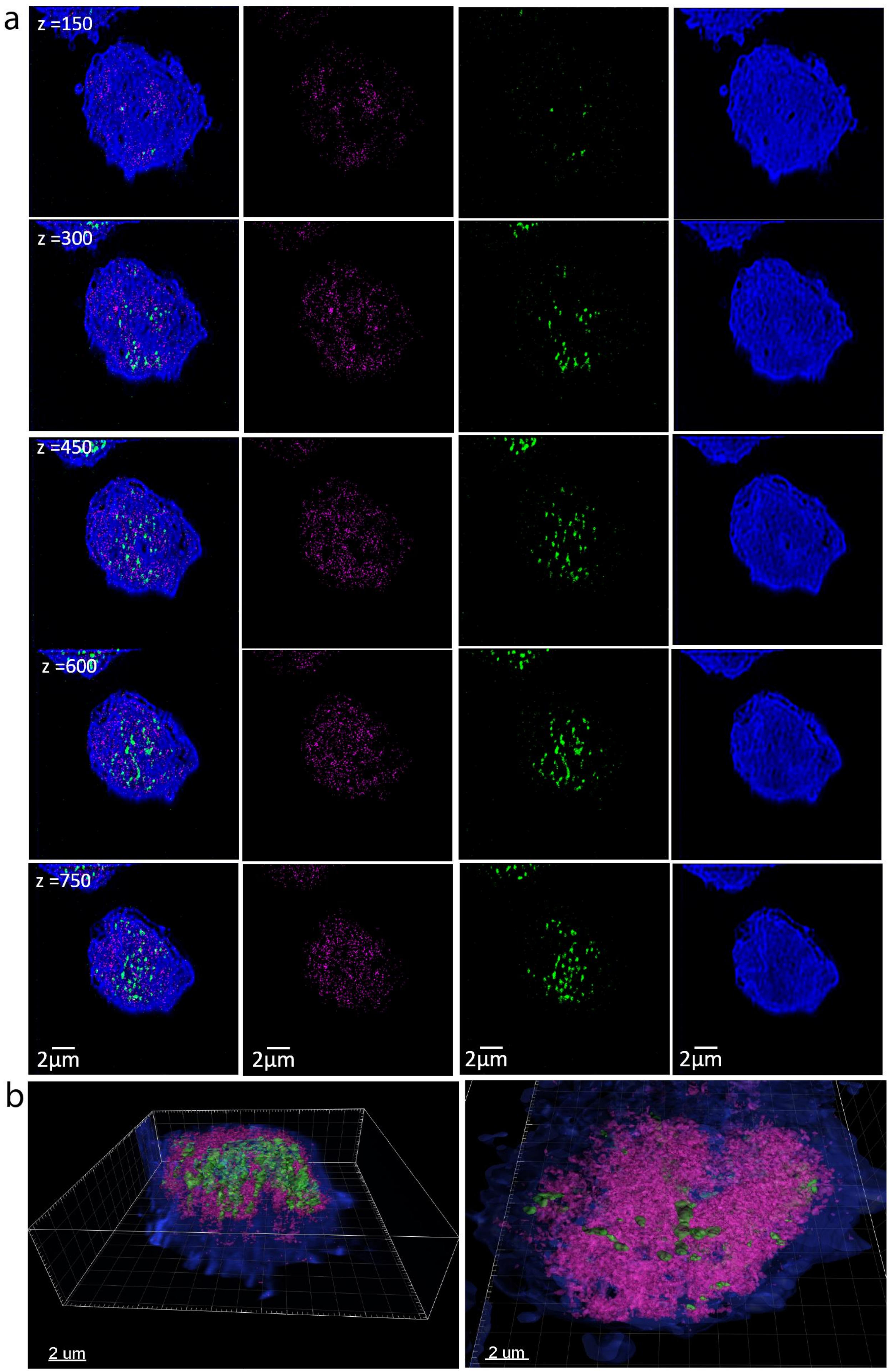

a
z =150
z =300
z =450
z =600
z =750
2μm
2μm
2μm
2μm
b
2 um
2 um

**Supplementary Figure 27.** Biological replicate data for nascent transcripts and replication (30-min pulse).

(a) Nalm6 cell was simultaneously incubated with $^{127}$I-dU (green) to visualize replication and $^{81}$Br-rU (magenta) to monitor transcription for 30 minutes. Tomographic ion slices for 150$^{th}$, 300$^{th}$, 450$^{th}$, 600$^{th}$, and 750$^{th}$ depths were presented. $^{31}$P (blue) showed folding chromatin patterns.

(b) 3D renders of 750 slices for the Nalm6 cell. Replication sites (green), nascent transcripts (magenta), and chromatin (blue). Spatial isolation of RNA transcripts and replication forks was detected.

Note that spatial regions that look slightly darker in the rightest column (chromatin) exhibited higher transcription, yielding a direct visualization of globally decondensed chromatin landscape.

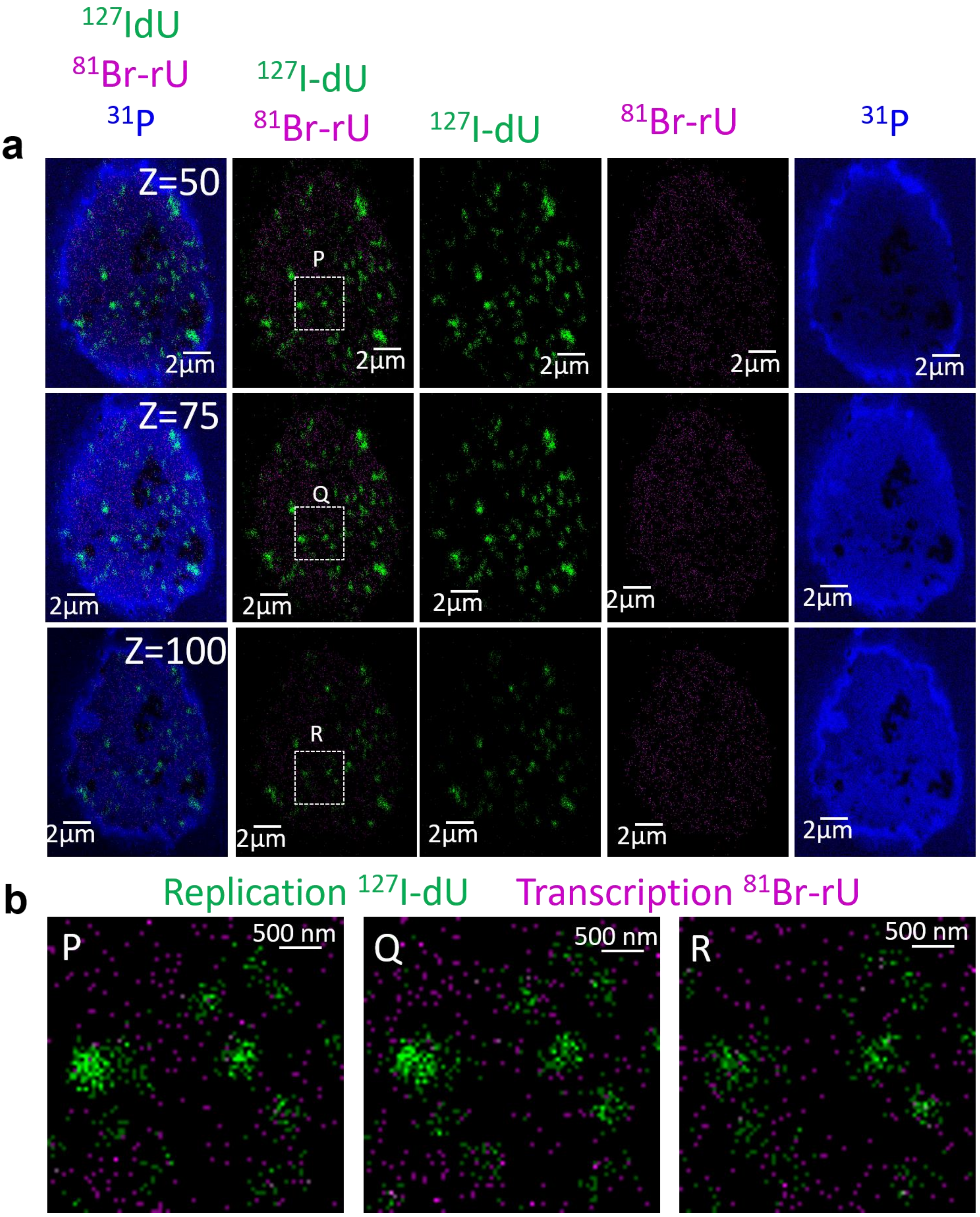

a
$^{127}$IdU
$^{81}$Br-rU
$^{31}$P
$^{127}$I-dU
$^{81}$Br-rU
$^{127}$I-dU
$^{81}$Br-rU
$^{31}$P
Z=50
Z=75
Z=100
P
Q
R
2μm
b
Replication $^{127}$I-dU
Transcription $^{81}$Br-rU
P
Q
R
500 nm

**Supplementary Figure 28.** Another replicate for simultaneous analysis of transcription and replication by IBT.

(a) Nalm6 cells were incubated with $^{127}$I-dU and $^{81}$Br-rU for 30 minutes in culture. Ion images for each channel was presented in each row for 50$^{th}$, 75$^{th}$, and 100$^{th}$ slices.

(b) Pixels from two distinct images for newly synthesized RNAs (magenta) and DNAs (green) do not overlap in the zoomed regions (P, Q, and R), demonstrating a spatial segregation of transcription and replication.

Note that these images were not processed by the mathematical pipeline, rather raw images are presented. 512x512 scans were used at 2 pA imaging current.

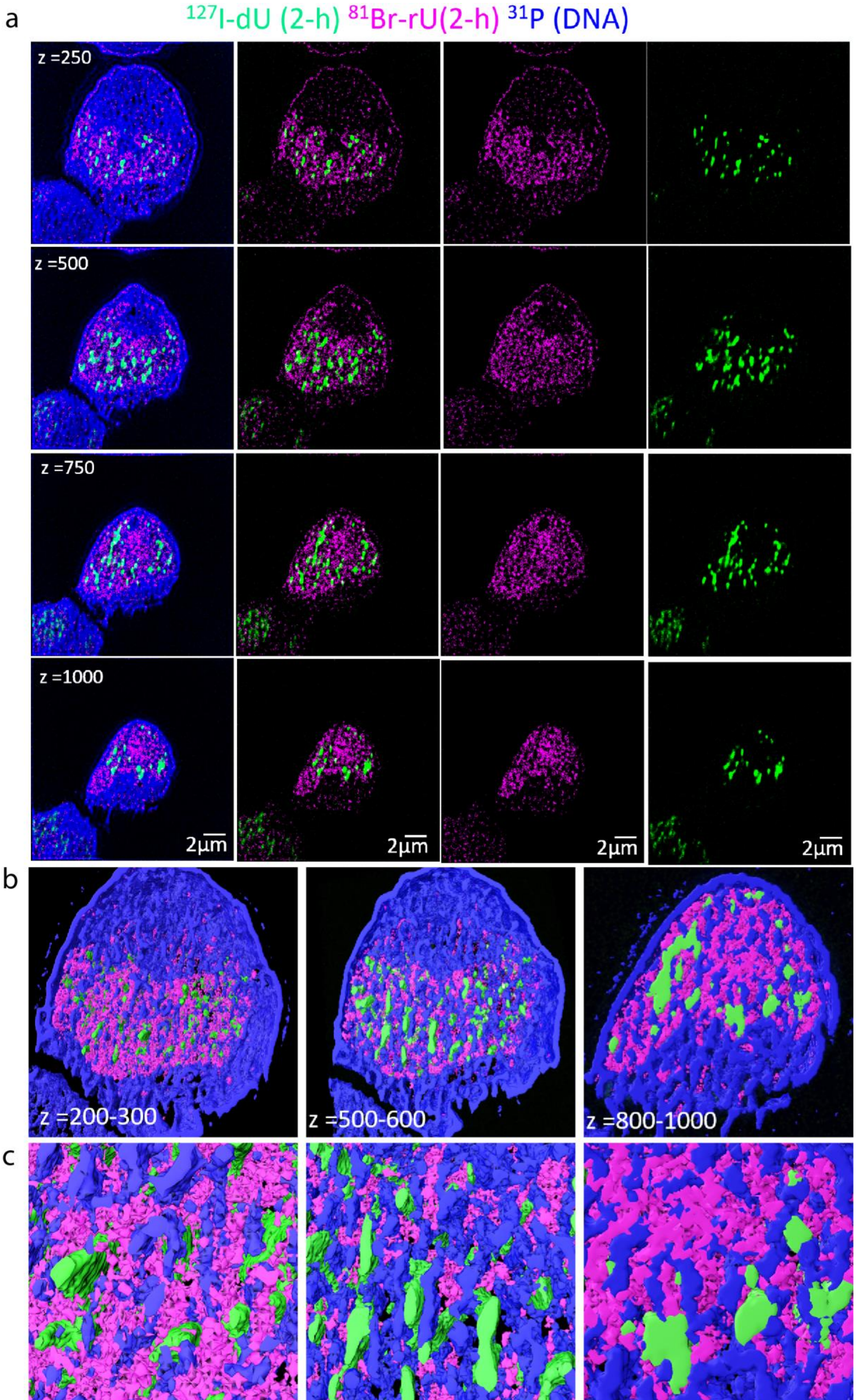
a
127I-dU (2-h) 81Br-rU(2-h) 31P (DNA)
z =250
z =500
z =750
z =1000
2μm
2μm
2μm
2μm
b
z =200-300
z =500-600
z =800-1000
c

**Supplementary Figure 29.** Visualization of transcription and replication after 2-h pulse with $^{127}$I-dU and $^{81}$Br-rU.

(a) Digitally processed images for IBT at 250$^{th}$, 500$^{th}$, 750$^{th}$, and 1000$^{th}$ slices. Chromatin was detected at $^{31}$P (blue), replication forks by $^{127}$I-dU (green) and nascent transcripts by $^{81}$Br-rU channel (magenta). Long incubation time led to longer replication sites and higher transcription signal. Spatial isolation of replication and transcription was preserved even at this long pulse experiment.

(b) Partial 3D renders from more than 100 slices: 200-300, 500-600, and 800-1000 depths. Chromatin patterns and the spatial distribution of newly synthesized RNA and DNA around original chromatin were visualized.

(c) Zoomed images from previous 3D renders. Fine features of replication and transcription were detected. Spatial segregation model is further validated in these images from 2-h incorporation of metabolic labeling.

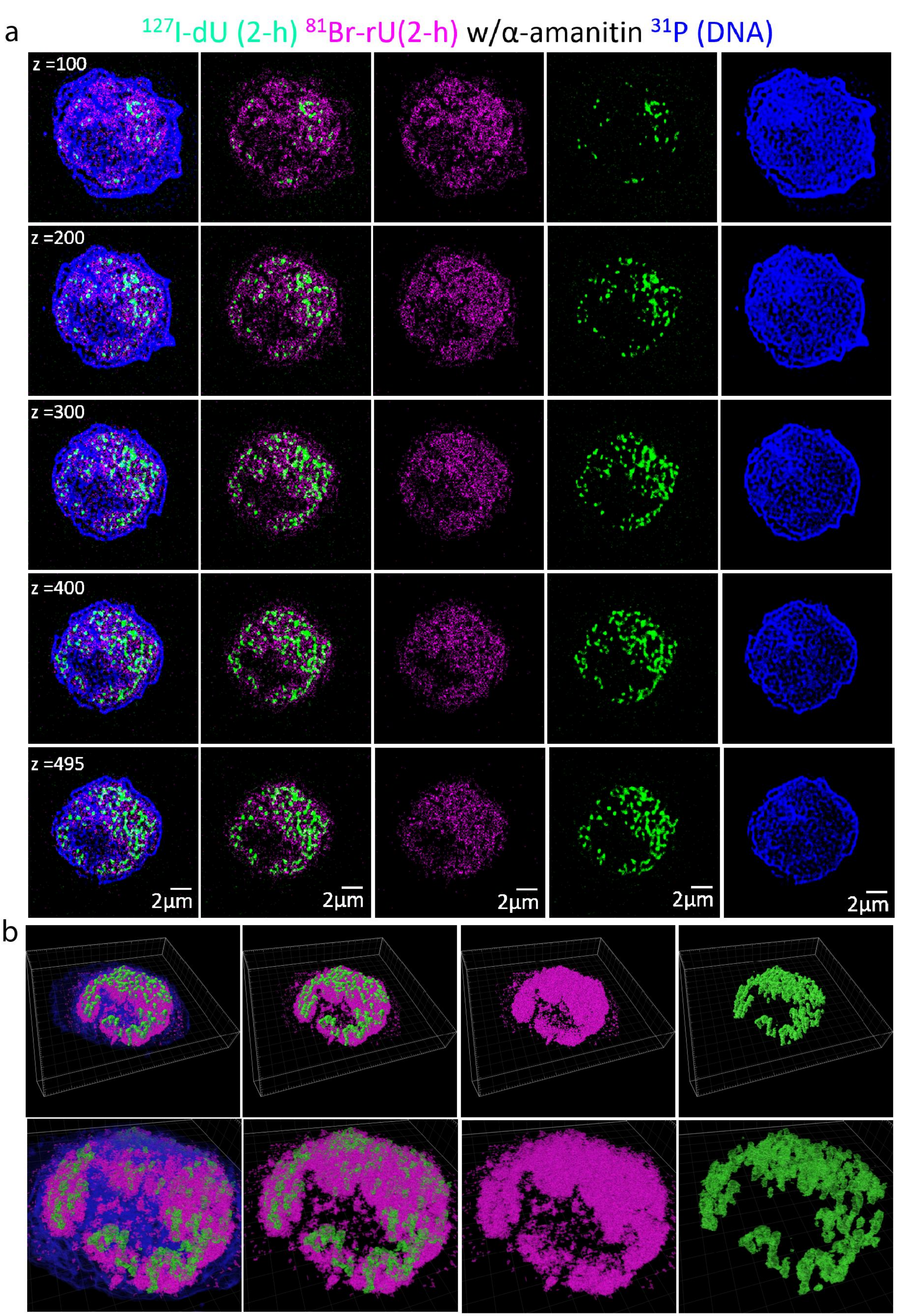
a
127I-dU (2-h) 81Br-rU(2-h) w/α-amanitin 31P (DNA)
z =100
z =200
z =300
z =400
z =495
2μm
2μm
2μm
2μm
2μm
b

**Supplementary Figure 30.** Transcription inhibition during the labeling by α-amanitin for 2 hours at 10μM.

(a) Visualization of transcription (magenta) by $^{81}$Br-rU, replication (green) by $^{127}$I-dU, and chromatin (blue) by $^{31}$P signal. Processed ion images from 100$^{th}$, 200$^{th}$, 300$^{th}$, 400$^{th}$, and 495$^{th}$ slices were presented.

(b) 3D Renders of transcription and replication around chromatin at a viewing angle. Top row corresponds to solid surfaces and bottom row shows partially transparent surfaces in the IBT images.

Spatial distribution of replication and distribution was consistent with previous results:

- Isolation of transcribing DNA and replicating DNA in 3D.
- Enrichment of transcription in decondensed chromatin regions.

The effect of α-amanitin in active transcription:

- Total transcription signal reduced compared to the Supplementary Fig. 29 (same conditions, 2-h).
- Partial repression but not total transcription block.
- Replication did not get affected from transcription inhibitors.

a

Unnormalized transcription signal after inhibition

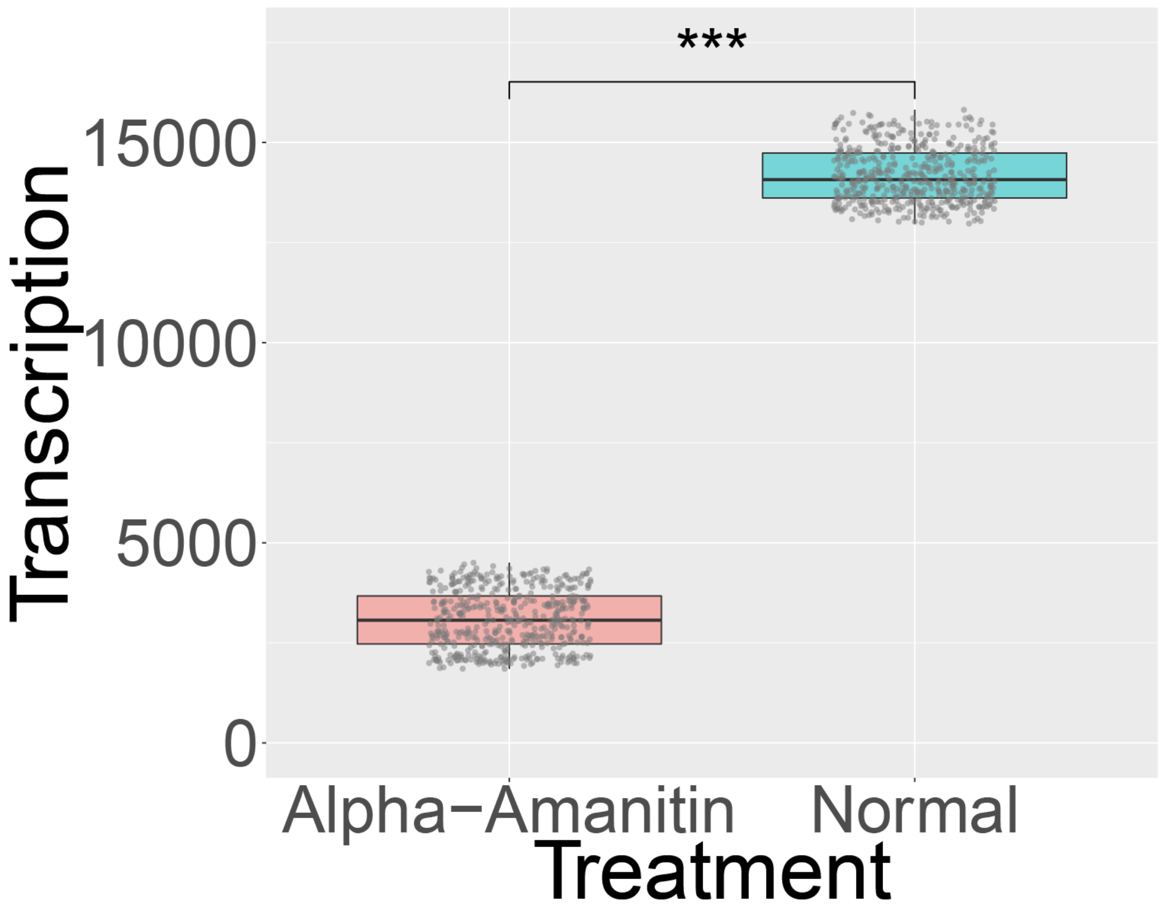


b

Normalized transcription signal after inhibition

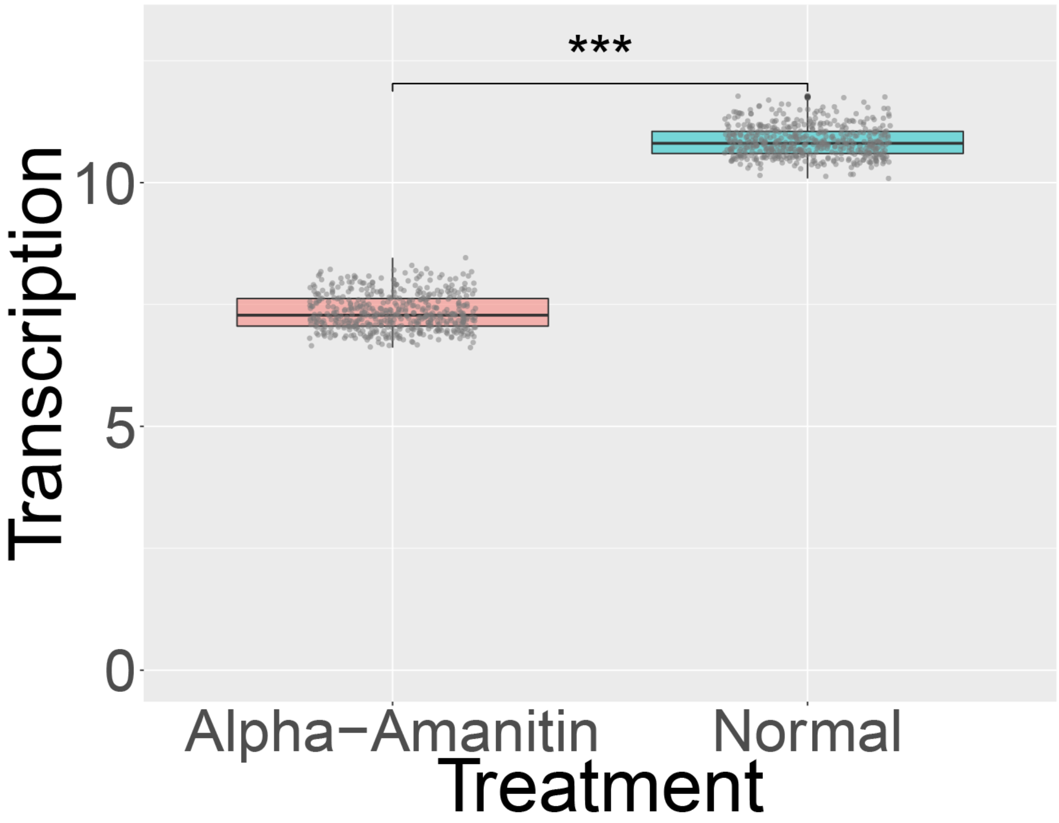

**Supplementary Figure 31.** Transcription inhibition reduced the nascent transcriptome signal.

(a) Statistics of total transcription signal at the $^{81}$Br-rU channel. Each data point corresponds to single depth from ion beam tomograms. It exhibited reduced signal down to the 0.21-fold of the transcription without inhibition. Each scatter point corresponds to single depth slice from ion beam tomographic dataset.

(b) Normalization of the transcription signal to the cell area per depth image. This step alleviates the issues of cell size on transcription signal analysis. It showed reduction down to 0.67-fold of the original transcription signal without inhibition. Each dot is for single slice in ion tomograms.

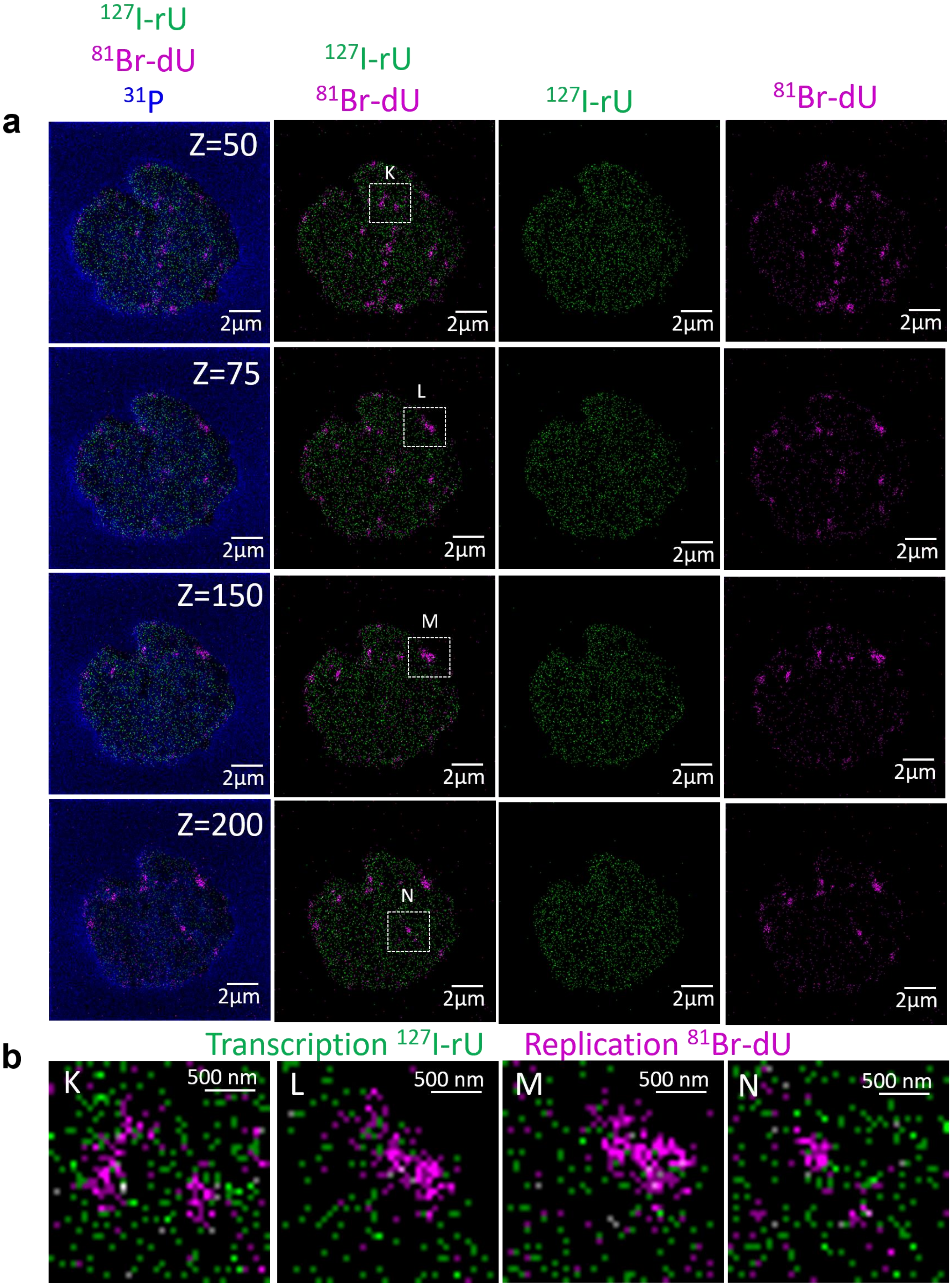

$^{127}$I-rU
$^{81}$Br-dU
$^{31}$P
a
Z=50
Z=75
Z=150
Z=200
K
L
M
N
2µm
b
Transcription $^{127}$I-rU
Replication $^{81}$Br-dU
500 nm

**Supplementary Figure 32.** IBT results do not depend on label. The experiment is the same as described in Supplementary Figure 18 except that transcripts were labeled with $^{127}$I-rU and DNA was labeled with $^{81}$Br-dU.

(a) Ion images for each row corresponds to a 50$^{th}$, 75$^{th}$, 150$^{th}$, and 200$^{th}$ slices.

Note that these images were not processed by the mathematical pipeline, rather raw images were presented to describe spatial differences of individual pixels. 256x256 scans were used at 2 pA imaging current.

(b) Zoomed images from distinct slices (K, L, M, and N) showed non-overlapping pixels for newly synthesized transcripts and DNAs.

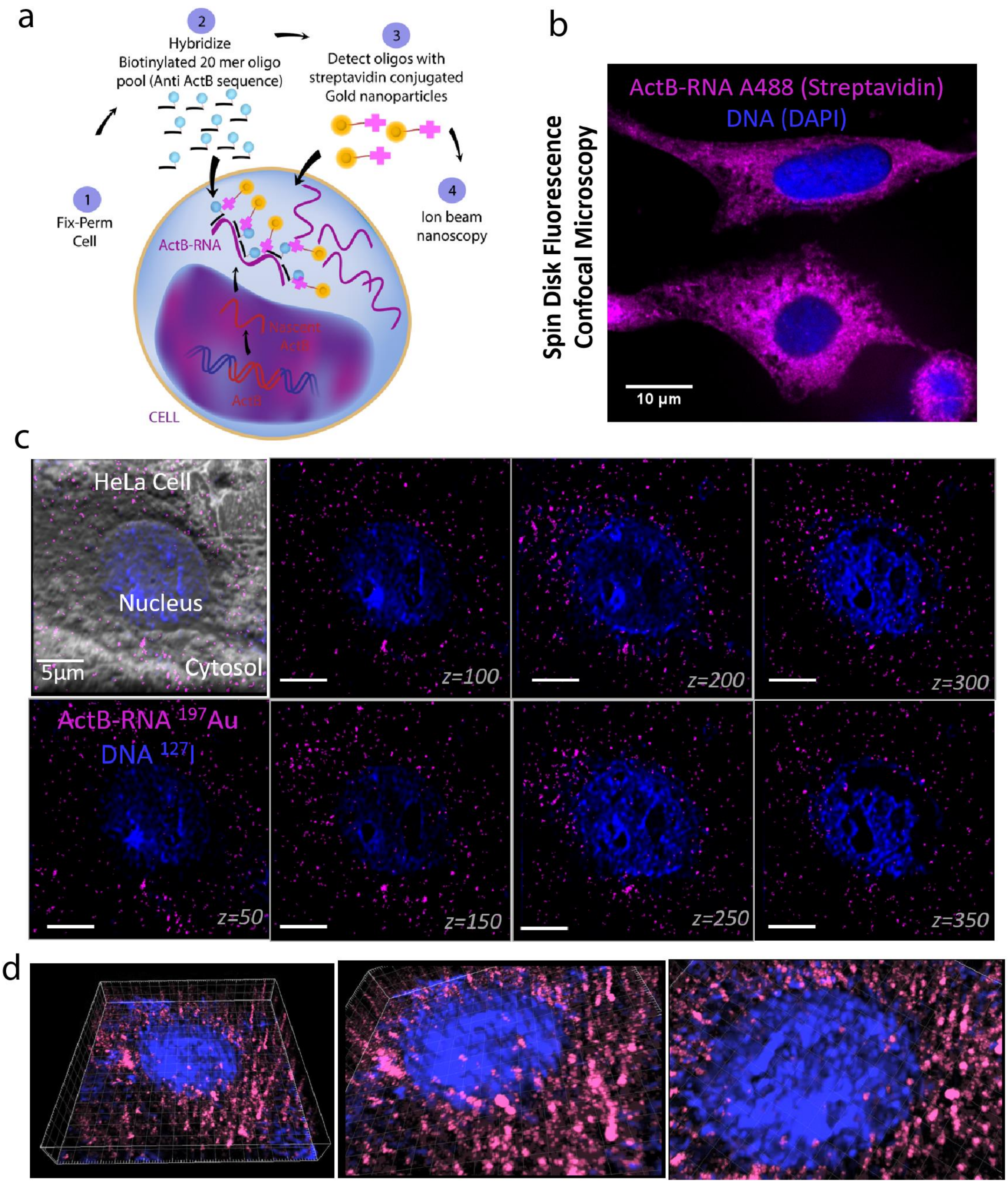
a
1
Fix-Perm
Cell
2
Hybridize
Biotinylated 20 mer oligo
pool (Anti ActB sequence)
3
Detect oligos with
streptavidin conjugated
Gold nanoparticles
4
Ion beam
nanoscopy
ActB-RNA
Nascent
ActB
ActB
CELL
b
Spin Disk Fluorescence
Confocal Microscopy
ActB-RNA A488 (Streptavidin)
DNA (DAPI)
10 μm
c
HeLa Cell
Nucleus
Cytosol
5μm
z=100
z=200
z=300
ActB-RNA 197Au
DNA 127I
z=50
z=150
z=250
z=350
d

**Supplementary Figure 33. Ion beam tomography maps transcriptional activity in individual HeLa cells**.

(a) RNA FISH was used to visualize *ActB* mRNA in a HeLa cell. After fixing and permeabilization of the cells, twenty-four biotinylated DNA probes designed to hybridize to the *ActB* mRNA were added to the cells. Streptavidin-conjugated gold nanoparticles were then incubated with the cells. Ion beam imaging was used to detect the probes in the $^{197}$Au channel.

(b) Using streptavidin-A488 secondary conjugates against oligonucleotides, we validated detection of *ActB* (magenta) in HeLa cells by spin disk fluorescence confocal microscopy. DAPI was used to visualize DNA (blue). Scale bar 10 µm. Most of these transcripts were mature, but we captured a few transcripts in nuclear regions.

(c) IBT images of *ActB* mRNA in the 25-µm wide central part of a HeLa cell acquired across 450 slices. $^{127}$I-dU (blue) was used to label DNA. Scale bars, 5 µm.

(d) 3D renders of ion beam tomographic RNA images for *ActB* in a single HeLa cell from different viewing angles.

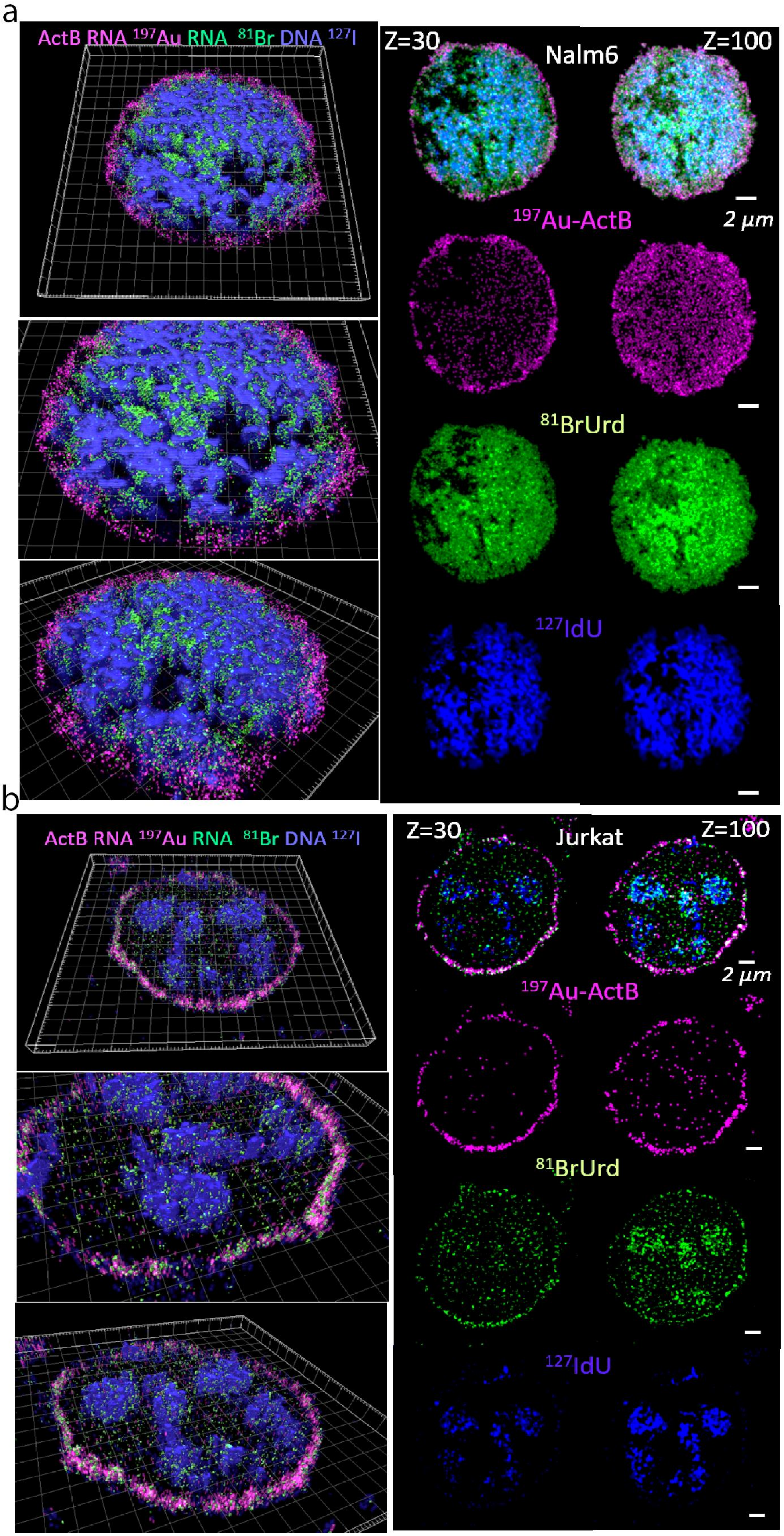

a
ActB RNA 197Au RNA 81Br DNA 127I
Z=30
Nalm6
Z=100
2 μm
197Au-ActB
81BrUrd
127IdU
b
ActB RNA 197Au RNA 81Br DNA 127I
Z=30
Jurkat
Z=100
2 μm
197Au-ActB
81BrUrd
127IdU

**Supplementary Figure 34. Newly synthesized *ActB* transcripts and global transcription were measured together by ion beam tomography in Nalm6 and Jurkat cells**.

(a and b) IBT images of DNA in the $^{127}$I channel (blue, cells were labeled for with $^{127}$I-dU for 23 hours prior to RNA labeling), newly synthesized RNAs in the $^{81}$Br channel (green, labeled with $^{81}$Br-rU for 1 hour after 1 hour of pulse), and *ActB* in the $^{197}$Au channel (magenta, labeled with smFISH). The 30$^{th}$ and 100$^{th}$ slices are presented from each channel, demonstrating that *ActB* mRNA preferentially localizes at the periphery of both a) a Nalm6 cell and b) a Jurkat cell.

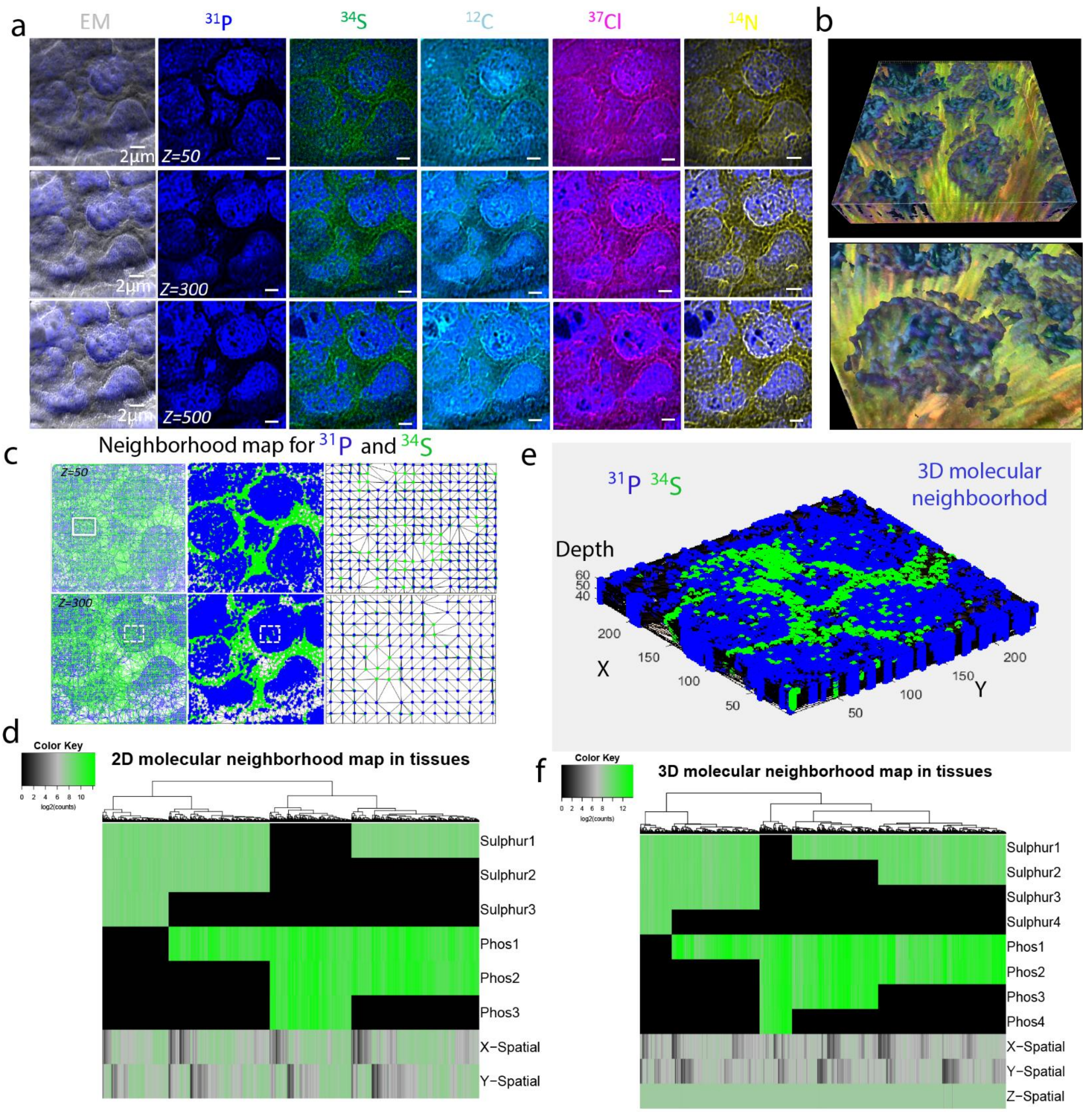

a
EM
31P
34S
12C
37Cl
14N
2µm
Z=50
Z=300
Z=500
b
c
Neighborhood map for 31P and 34S
Z=50
Z=300
e
31P 34S
3D molecular neighboorhod
Depth
X
Y
d
Color Key
log2(counts)
2D molecular neighborhood map in tissues
Sulphur1
Sulphur2
Sulphur3
Phos1
Phos2
Phos3
X−Spatial
Y−Spatial
f
Color Key
log2(counts)
3D molecular neighborhood map in tissues
Sulphur1
Sulphur2
Sulphur3
Sulphur4
Phos1
Phos2
Phos3
Phos4
X−Spatial
Y−Spatial
Z−Spatial

**Supplementary Figure 35. Molecular neighborhood analysis of chromatin and proteins in FFPE tissues using IBT.**

(a) A label-free ion imaging panel for $^{31}P$ (primarily chromatin, blue), $^{34}S$ (proteins, green), $^{12}C$ (cyan), $^{37}Cl$ (red), and $^{14}N$ (yellow) in individual $CD3^{+}$ T cells from a lymph node biopsy core of a T cell lymphoblastic lymphoma patient. Sub-100-nm resolution images were acquired across 500 depth sections. Scale bars, 2 µm. Images in the first column show secondary total ion images as a bright field contrast of the cells. Each row is a 2D representation for $50^{th}$, $300^{th}$ and $500^{th}$ slices from ion tomograms.

(b) IBT renders of single T cells with $^{31}P$ (blue), $^{34}S$ (green), and $^{37}Cl$ (red) co-localization resulting in yellow colors around chromatin islands.

(c) Molecular neighborhood map of DNA ($^{31}P$) and protein ($^{34}S$) interactions in tissues. Two distinct depth images for $50^{th}$ and $300^{th}$ slices were processed by Delaunay triangles that were cornered with blue dots ($^{31}P$) or green dots ($^{34}S$).

(d) Signals of each center-of-mass sub-regions were clustered by heatmap together with spatial positions (X-spatial and Y-spatial) of each triangle. Signals for $^{31}P$ were highly enriched in the nucleus, and other spatial sub-regions exhibit a combination of $^{31}P$ and $^{34}S$ signals. Delaunay triangles were cornered with a combination of phosphate (1-3) sulphur (2-3) signals.

(e) 3D neighborhood analysis of twenty sections with Delaunay tetrahedrons that were cornered by four unique green ($^{31}P$) or magenta ($^{34}S$) dots.

(f) Distinct neighborhoods were detected in the heatmap after clustering $^{31}P$ and $^{34}S$ with X-Y-Z spatial positions of tetrahedrons. Phosphate (1-4) and Sulphur (1-4) signals corner the tetrahedrons. Chromatin-protein interactions and protein-protein interactions exhibited spatial variations in different clusters.

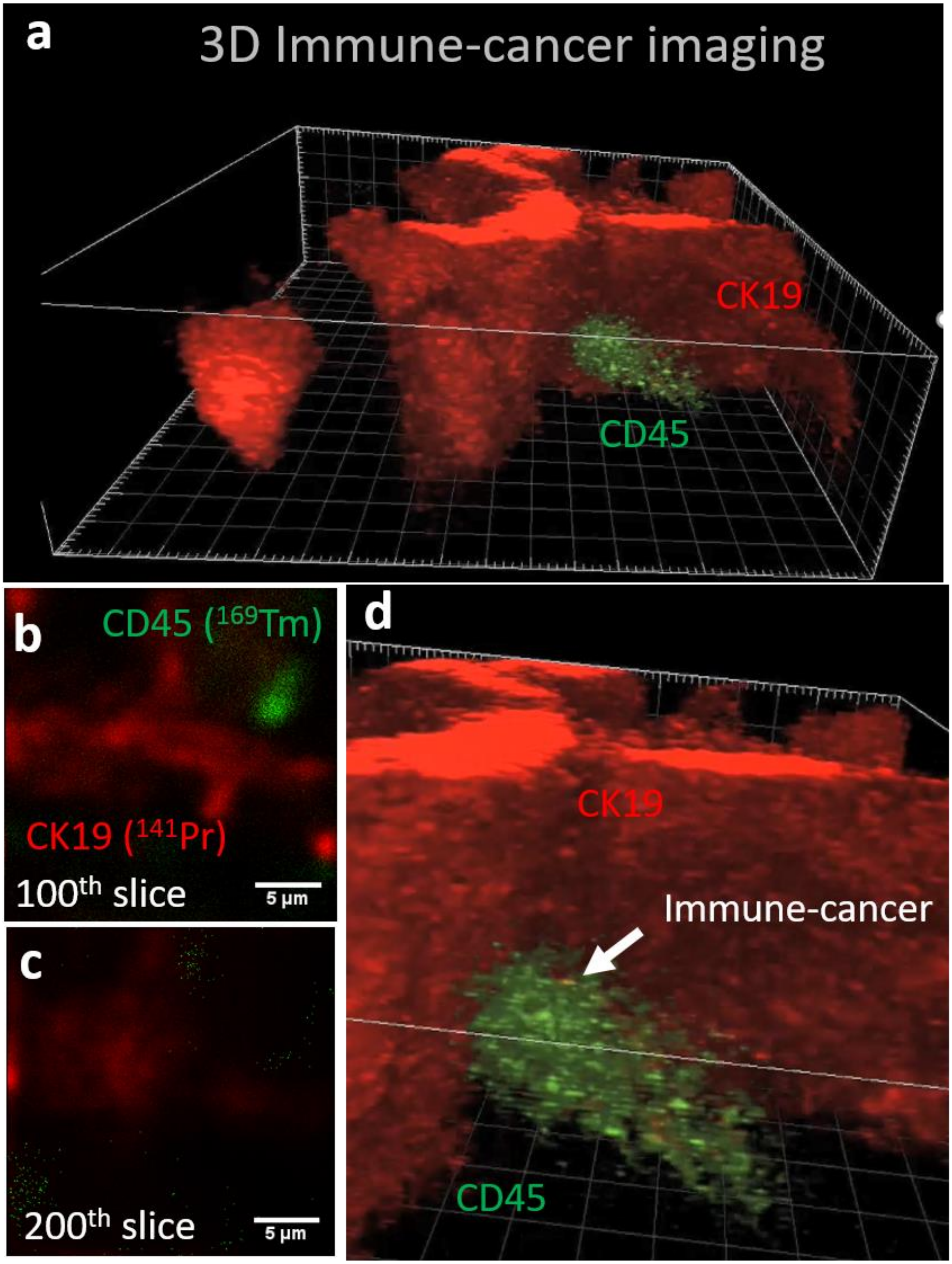
a
3D Immune-cancer imaging
CK19
CD45
b
CD45 (169Tm)
CK19 (141Pr)
100th slice
5 µm
c
200th slice
5 µm
d
CK19
Immune-cancer
CD45

**Supplementary Figure 36.** Volumetric histology by IBT.

(a) A sample of cholangiocellular carcinoma from a multi-tumor tissue microarray at 10-µm thickness was mounted on a silicon wafer for ion beam imaging by oxygen ion beam source.

(b) Immune cell ($CD45^+$, label: $^{169}Tm$) and cancer cells (CK19, label: $^{141}Pr$) were visualized.

(c) IBT collected at 420 depths allows digital focusing onto different layers ($100^{th}$ and $200^{th}$ slices shown).

(d) 3D renders of connected cancer cells (red) and an immune cell (green).

**Supplementary Methods**

**Tissues**

Paraffin blocks of 70 unique different tissues were obtained from the archive of the Institute of Pathology, University of Bern, Switzerland. Cancer and normal tissue regions were annotated on corresponding hematoxylin and eosin stained sections. A tissue microarray of 70 cores measuring 0.6 mm in diameter was assembled using an automated tissue microarrayer (TMA Grand Master, 3D Histech). Prior to slicing, silicon substrates were treated with Vectabond (Vector Laboratories) to improve attachment of tissues on the surface. The tissue microarray block was then sliced onto the silicon substrates (18 mm x 18 mm) at 5-, 10-, and 15-µm thickness. The final silicon samples with tissues were preserved in vacuum desiccator in the dark. To deparaffinize the tissue, the silicon substrate was manually treated sequentially with xylene (3×), 100% ethanol (2×), 95% ethanol (2×), 80% ethanol, 70% ethanol, and $H_2O$, each for 30 s. To perform antigen retrieval, the tissue was then placed in a plastic slide holder in target retrieval solution (Agilent, Dako) and loaded into a thermal cycler (PT Module, 97 $^{o}$C for 40 min and cool down to 65 $^{o}$C). After cooling to room temperature, the tissue was rinsed with a wash buffer (PBS 1× IHC with Tween 20 and 0.1% BSA). The sample then was dehydrated and directly imaged by IBT for label-free imaging of subcellular features (**Supplementary Fig. 35**).

**Tissue staining**

To enable labeling of mass reporters, after the antigen retrieval step, the tissue on silicon was treated with blocking buffer (TBS 1× IHC with Tween 20 and 3% donkey serum) for 1 h.  After removing blocking buffer, the antibody mix ($CD45^{+}$, label: $^{169}Tm$ and CK19, label: $^{141}Pr$) was loaded onto the sample and incubated overnight at 4 $^{o}$C. The next day, the sample was rinsed with wash buffer and post-fixed with 2% glutaraldehyde (Electron Microscopy Sciences) for 5 min. After rinsing the sample with low barium PBS, the sample was washed with DI water and dehydrated for ion beam tomographic experiments (**Supplementary Fig. 36)**.
